# Phylogenetic meta-analysis implicates large brains and our unusual posture in human handedness

**DOI:** 10.1101/2025.06.17.660131

**Authors:** Thomas A. Püschel, Rachel M. Hurwitz, Chris Venditti

**Affiliations:** Institute of Human Sciences, School of Anthropology and Museum Ethnography, University of Oxford, Oxford, UK; School of Biological Sciences, University of Reading, Reading, UK

## Abstract

Humans exhibit a striking and near-universal population-level right-hand preference, an evolutionary singularity unmatched among primates. Despite its pervasiveness, the origins of this lateralisation remain poorly understood. Here, we combine phylogenetic comparative methods with meta-analysis to investigate manual lateralisation across 41 anthropoid species (n = 2,025), testing longstanding eco-evolutionary hypotheses for handedness direction (MHI) and strength (MABSHI). Our models reveal significant phylogenetic signal for both traits and identify *Homo sapiens* as an evolutionary outlier, exhibiting exceptional rightward bias and strength relative to phylogenetic expectations. However, this outlier status disappears when brain size (endocranial volume, ECV) and intermembral index (IMI) are included, suggesting these factors are central to the emergence of human handedness. We also show that high MABSHI evolved early in hominin evolution, while MHI increased to unparalleled levels with the appearance of the genus *Homo*. Our findings implicate bipedalism and neuroanatomical expansion as key drivers of uniquely human lateralisation, while also revealing broader ecological patterns shaping handedness across primates. This work provides a framework for disentangling human-specific adaptations from general primate trends in the evolution of behavioural asymmetries.

---

In all human cultures across every corner of the globe about 90% of people favour their right hand ^1–4^. Some have argued that this has been true since the Neolithic ^5^, whilst others contend that it has been constant through the entire *Homo* lineage ^6–11^. Some level of directional manual lateralization is present in sub-populations of various primate species, but the level and consistency of handedness in humans is unmatched, and despite much interest, still represents an unexplained singularity ^4,11–15^.

The neurological basis of handedness is known to be rooted in specialised brain regions, but the evolutionary origins and dominance of right-handedness remain enigmatic ^4,16–20^. Several eco-evolutionary hypotheses involving tool use, diet, brain size, locomotor strategy, terrestriality, among many other potential factors, have been proposed to explain this pervasive preference (Figure 1; Supplementary Table 1). However, many of these have only been posited in only ambiguous and descriptive terms making them difficult to test and hampering progress in the area. This has resulted in many conflicting hypotheses concurrently existing in the literature without any clear tests of their applicability.

**Figure 1.**
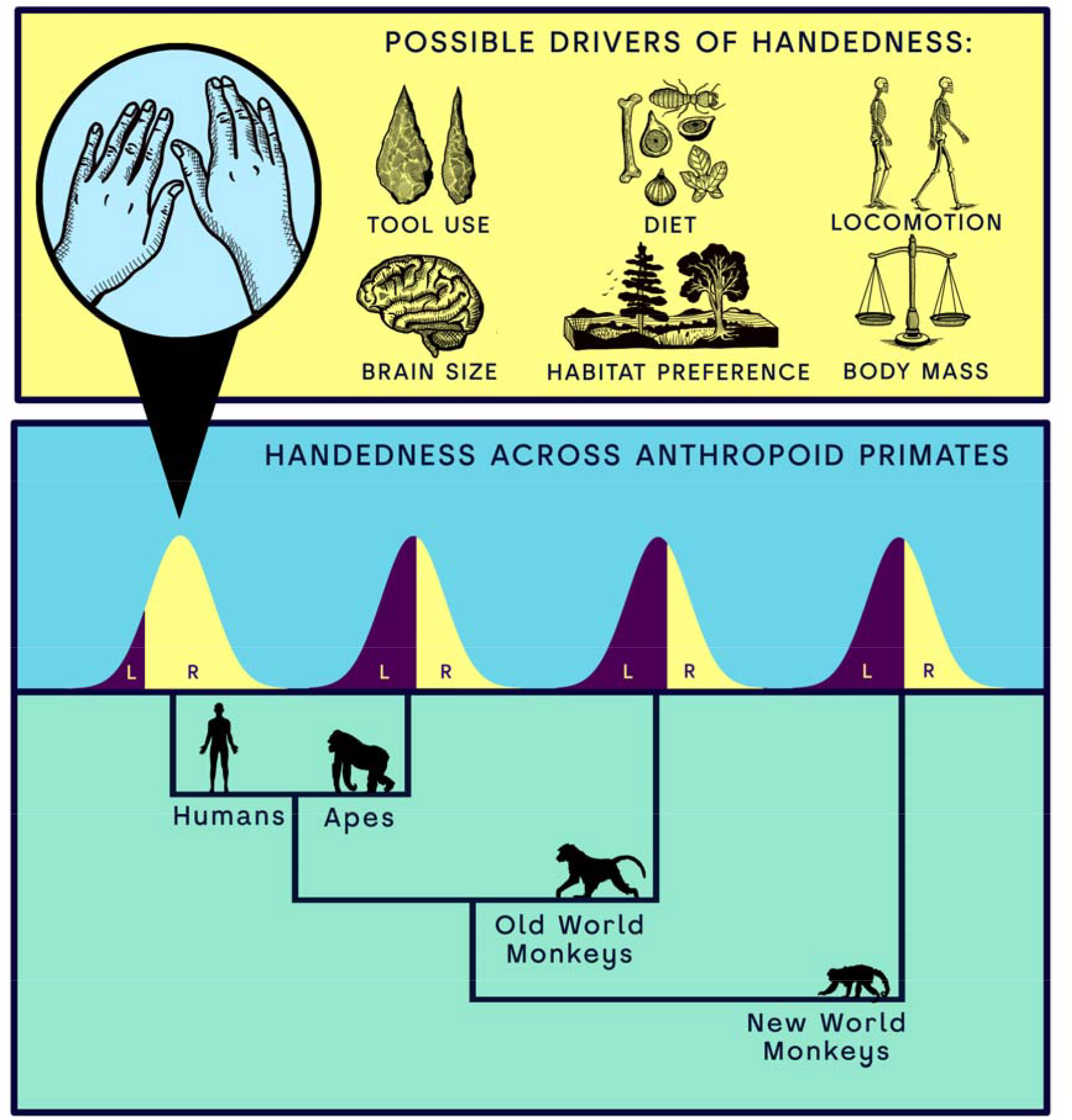
Drivers often proposed to explain the unique pattern of human handedness direction. Humans show an unparalleled level of handedness rightwards bias as compared to other anthropoid clades.

Handedness in non-human primates, has been understudied, especially from a comparative perspective. A recent meta-analysis found that, unlike humans, non-human anthropoids do not show population-level handedness but do exhibit strong manual preferences during bimanual tasks ^21^. A phylogenetic study confirmed that human right-handedness is an extreme case, with population-level handedness being rare in non-human primates ^14^. The latter also found no strong link between manual lateralization and factors like tool use, substrate preference, or brain size, but noted that terrestrial species had weaker hand preferences than arboreal ones.

No combined phylogenetic comparative and meta-analytical method has been used to study anthropoid handedness, leaving key eco-evolutionary hypotheses untested. This approach offers several advantages, including better identification of variation drivers, improved control of data biases, and more accurate phylogenetic conclusions .^22^ Although comparative and meta-analytical methods are rarely combined in evolutionary studies, they reduce sampling errors and help address research biases, such as the overrepresentation of certain charismatic species (e.g., chimpanzees)^23-25^ . This study integrates both methods to assess hypotheses on primate and human handedness.

We applied Bayesian Phylogenetic Comparative Meta-Analytical methods to assess handedness patterns across anthropoids. Handedness was evaluated based on two facets: direction (mean handedness index, MHI) and strength (mean absolute handedness index, MABSHI). Our data include a recent meta-analysis^21^ and recent experimental data^14^ (see Methods), resulting in a standardized dataset of 2,025 individuals across 41 anthropoid species. To test the influence of phylogeny and previously proposed hypotheses on handedness in humans and other primates, we compiled relevant covariates (e.g., body mass, brain size, tool use, intrasexual competition) from various literature sources (Fig. 2; Supplementary Data 1; see Methods). Multiple lines of evidence suggest that *Homo sapiens* is an evolutionary outlier in handedness, showing extreme rightward bias. We therefore conduct our hypothesis testing including and excluding *H. sapiens*. In addition, to explicitly assess our potential evolutionary distinctiveness we use a ‘Phylogenetic Outlier’ test^26^ . We also use our phylogenetic meta-analytical models to predict MHI and MABSHI values for extinct hominin species, based on their phylogenetic positions and associated predictor variables.

**Figure 2.**
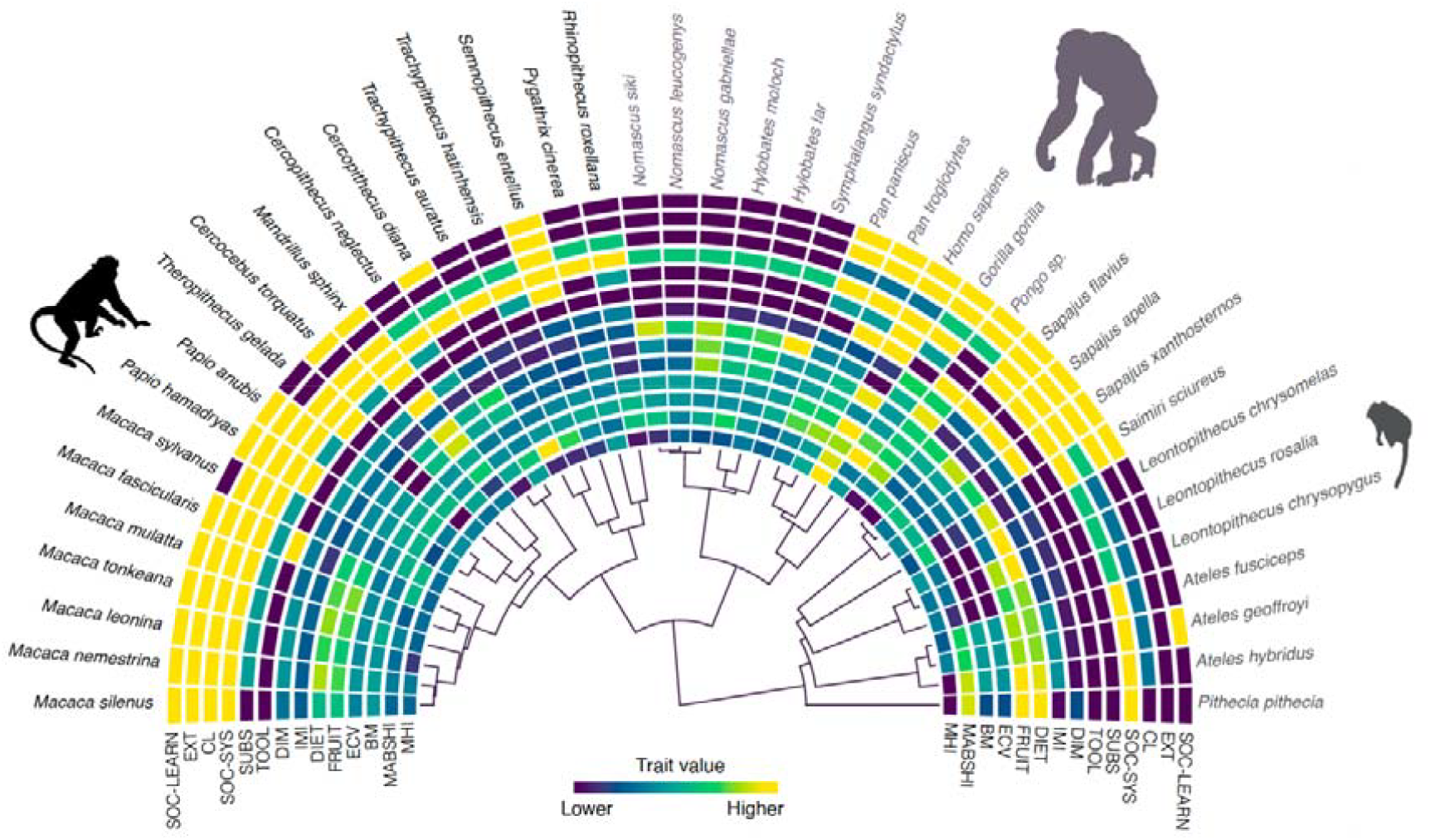
Traits analysed in the present study and maximum credibility clade tree for the analysed anthropoid species. The acronyms correspond to: MHI: mean handedness index; MABSHI: mean absolute handedness index; BM: body mass [kg]; ECV: endocranial volume [cm^3^]; FRUIT: percentage of fruits in diet; DIET: percentage of fruits and animals in diet; IMI: intermembral index; DIM: body mass sexual dimorphism; TOOL: tool use [0=absence, 1=presence); SUBS: substrate preference [0=arboreal, 1=both, 2=terrestrial]; SOC-SYS: social system [0=solitary, 1=pair, 2=group]; CL: intra-sexual competition levels sensu ^40^ [1,2,3,4]; EXT: extractive foraging [0=absence, 1=presence]; social learning sensu ^41^ [0=absence, 1=presence]. Traits are colour coded from lower to higher values. In the case of the discrete traits, colours also go from lower to higher values based on the categorisation provided in this legend. Cercopithecoidea, Hominoidea and Platyrrhini are represented by different silhouettes, as well as colours.

## Results

### Directionality of handedness

Contrary to previous studies we demonstrate that MHI shows significant phylogenetic signal (*h*^*2*^ = 0.48). This discrepancy likely stems from our approach, which adjusts for sampling error rather than relying on species means. There is no directional preference in MHI value across anthropoid species (MHI = -0.03, 95% CI: -0.32, 0.22) (Supplementary Figure 1). As expected, the highest MHI of 0.76 was found in *H. sapiens*, and although population-level handedness direction-biases were uncommon, other species including *Pan troglodytes, Gorilla gorilla*, and *Cercopithecus diana* showed a weak-moderate rightward preference (MHI > 0.17) (Supplementary Figure 1). Interestingly, there were more species displaying a stronger leftwards bias (MHI < -0.3) (e.g., *Pithecia pithecia, Ateles hybridus, Sapajus flavius, Pongo s*p., *Rhinopithecus roxellana, Cercopithecus neglectus)* (Supplementary Figure 1). However, it is important to keep in mind that in most cases, confidence intervals overlap with zero, indicating higher variability. In fact, *Pongo s*p. (MHI= -0.32) and *Rhinopithecus roxellana (*MHI= -0.32) are the only two taxa that show significant leftward preference, whereas humans are the only species that significantly favour the right hand.

When testing all the hypotheses listed in Supplementary Table 1 using MHI as the dependent variable, we found that no single hypothesis performed meaningfully better than any other, which was true both with and without humans (Supplementary Table 2). Interestingly, the inclusion of humans in the model routinely changed the significance of predictors in the model highlighting our species as evolutionary outliers See Supplementary Section 1). This was the case in every hypothesis that included brain size (measured by endocranial volume, ECV) and locomotor adaptations (measured by the intermembral index, IMI). The human IMI is extremely low, 72, reflecting that our hindlimbs (legs) are significantly longer than the forelimbs (arms), which is a key adaptation for bipedal locomotion. This provides evidence that that brain size and bipedal location have driven our exceptional MHI. After excluding *H. sapiens*, only ‘social system’, when testing the TU-SH-SSH hypothesis, is a significant predictor (which is explained by the orangutan) and BM in the SP-SS-B-TUH.

To better evaluate the role of the predictors identified as ‘significant’ for MHI, we created two new models, following the same modelling procedures as described before. Each model used only the predictors that were deemed relevant across all tested hypotheses. The first model, which excluded *H. sapiens*, included IMI, BM and SOC-SYS as covariates. The second model included *H. sapiens*, and featured DIET, ECV, IMI, TOOL, SUBS, BM, and SOC-SYS as fixed effects. We then used a model reduction process to iteratively remove those predictors that were not ‘significant’. SOC-SYS (social system: pair) was the only predictor that was ‘significant’ in the first model excluding *H. sapiens* after model reduction (Fig. 3a). The R^2^ value for this model was 0.26, indicating that social system only explains a limited amount of the variation in MHI patterns. The second model, resulted in only three predictors continuing to be ‘significant’, ECV, IMI and SOC-SYS (social system: pair) after model reduction (Fig. 3b). This model displayed a R^2^ of 0.42, thus showcasing the relevance of these three covariates.

**Figure 3.**
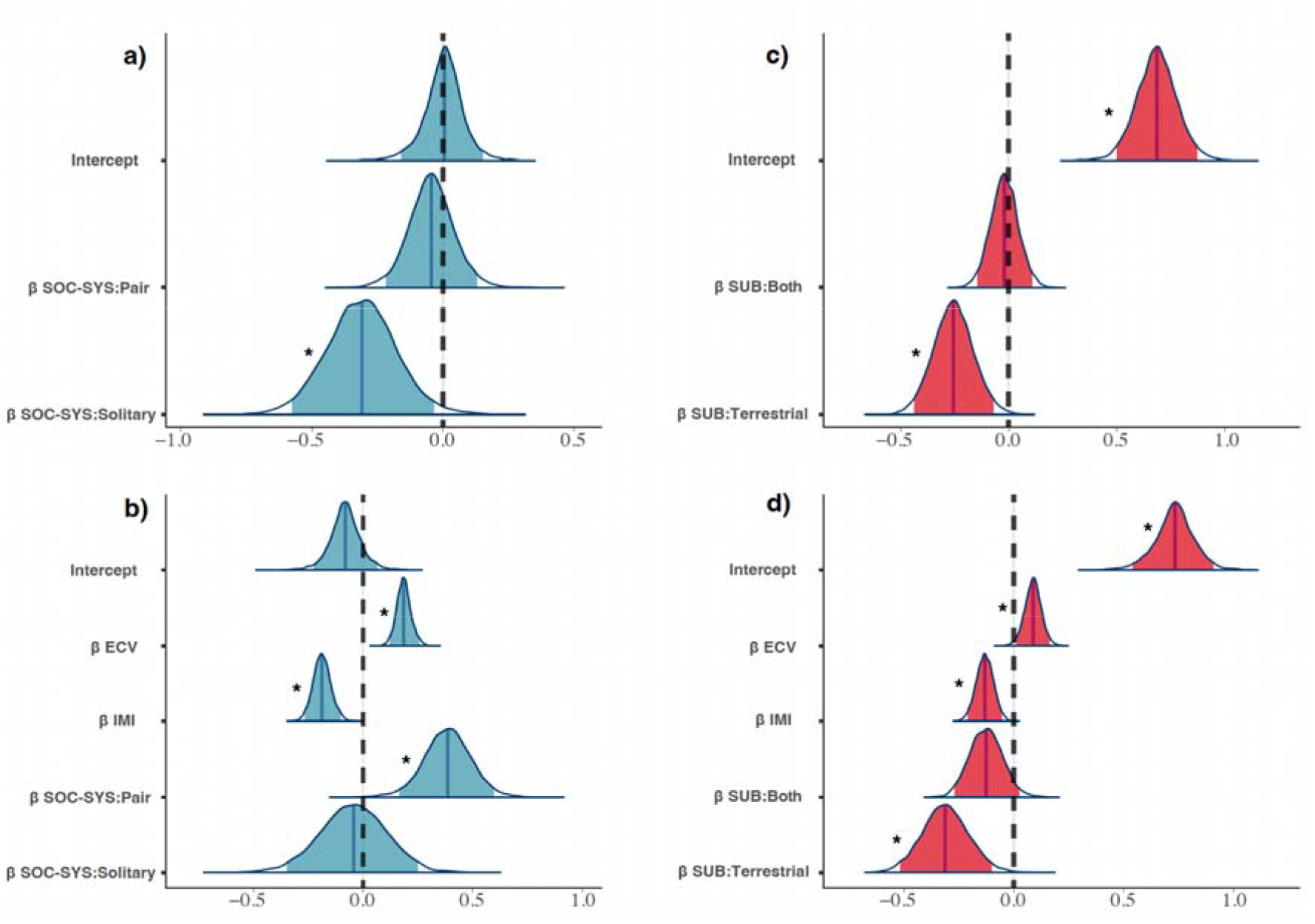
Coefficient Estimate Distributions for the Reduced Models. This figure shows the coefficient estimate distributions for the reduced models: (a) MHI excluding H. sapiens, (b) MHI including H. sapiens, (c) MABSHI excluding H. sapiens, and (d) MABSHI including H. sapiens. Significant effects are identified when the 95% credibility intervals (light coloured areas) do not overlap with zero (dotted line). The median for each coefficient distribution is represented by darker solid lines. Significant predictors are marked with asterisks.

These results suggest that *H. sapiens* are an outlier relative to the general primate trend. Therefore, we explicitly tested the status of humans as an evolutionary singularity using a phylogenetic outlier test (methods). Figure 4a displays both the observed MHI value for *H. sapiens* and the posterior distribution of MHI values derived from a model in which humans were excluded. The MHI prediction for the model using the predictor that was significant when excluding humans is 0.0, whereas the observed value is 0.76, highlighting the exceptional nature of human handedness direction relative to phylogenetic expectations.

**Figure 4.**
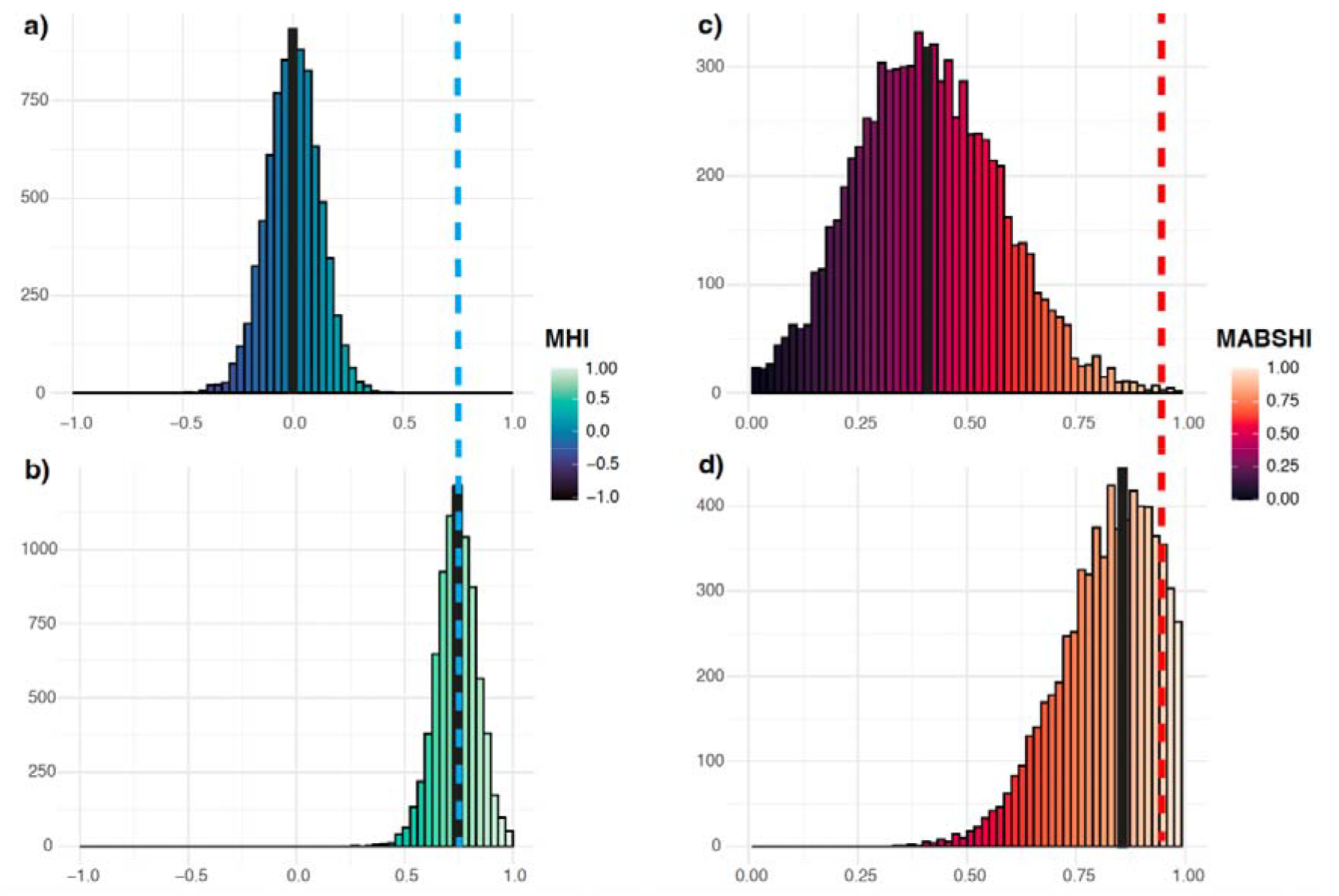
Phylogenetic Outlier Tests for MHI and MABSHI. This figure presents the posterior distribution of predictions for MHI and MABSHI for a new Homo sapiens observation. It illustrates these predictions based on the reduced models: (a) MHI excluding humans, (b) MHI including humans, (c) MABSHI excluding humans, and (d) MABSHI including humans. All predictions are marginalised over meta-analytical and non-phylogenetic species random effects.

This divergence implies that strong selective pressures would have been necessary to produce such a striking difference. Interestingly, *H. sapiens’* position as an outlier is no longer evident when ECV, IMI, and SOC-SYS (social system: pair) are included in the model, as the obtained MHI value (0.74) is almost identical to the observed one (0.76) (Fig. 4b). This suggests that selection pressures associated with these traits may explain the uniquely high MHI value observed in humans.

When using the reduced model including humans to predict hominin MHI values we found a clear trend of increasing MHI from older to more recent hominin species (Fig. 5): *Ardipithecus ramidus (*0.16), *Australopithecus afarensis* (0.32), *Homo ergaster* (0.50), *Homo erectus* (0.54), and *Homo neanderthalensis* (0.64). The notable exception is *Homo floresiensis*, which exhibits a comparatively weaker handedness directionality (MHI = 0.28). These results suggest that earlier hominins displayed weaker or less consistent hand preferences, whereas more recent species demonstrate stronger and more consistent lateralization.

**Figure 5.**
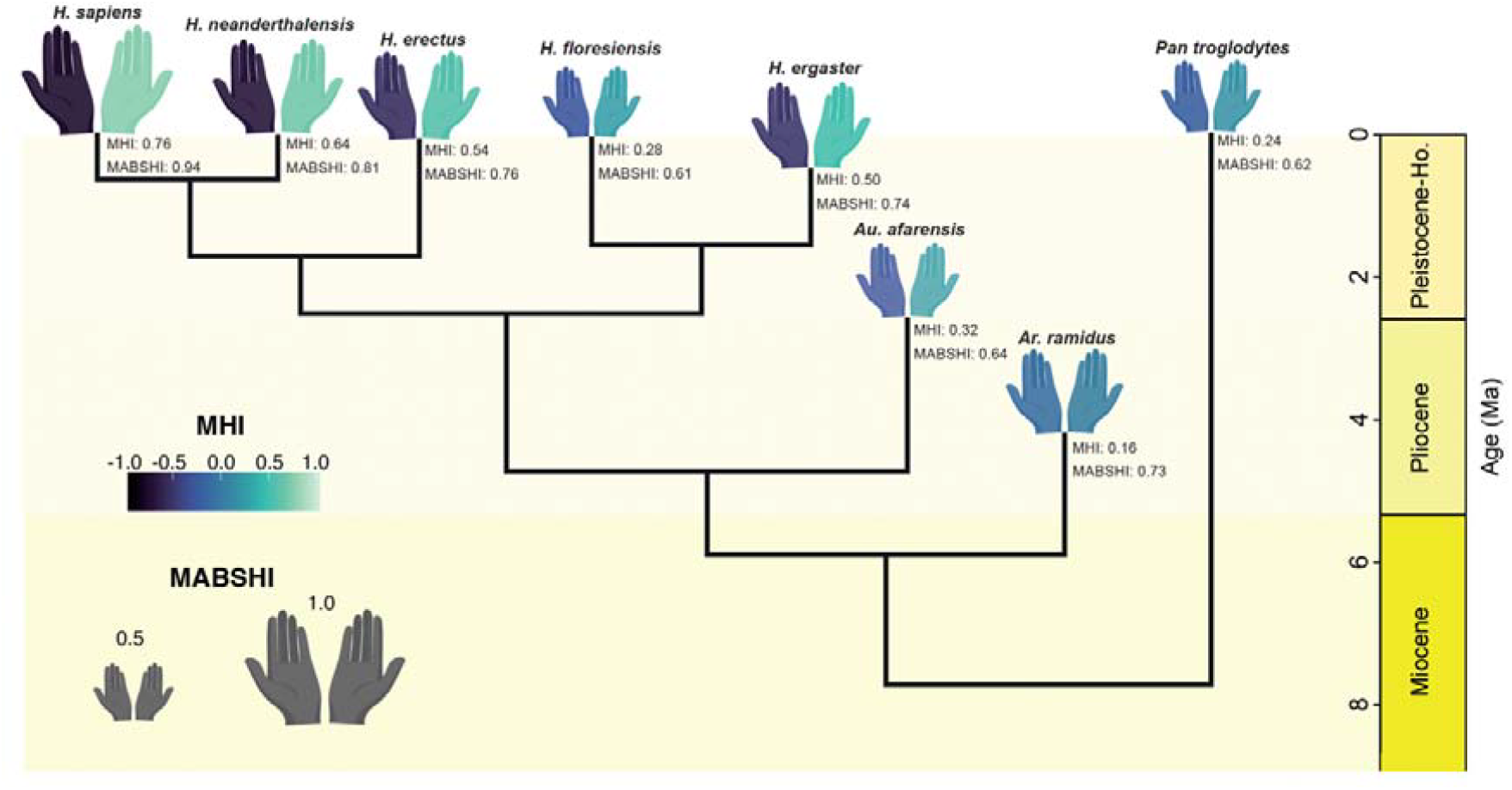
Predicted handedness in hominin species based on the reduced models including Homo sapiens. Right-hand colours represent the predicted magnitude of MHI; left-hand colours show the corresponding negative values (i.e., greater colour differences between hands reflect stronger handedness direction bias). Hand size is proportional to handedness strength, MABSHI. The phylogeny is the maximum a posteriori tree from^34^. Only hominin species with complete relevant covariate data were included in the analysis and are shown here. The handedness values for Pan troglodytes and Homo sapiens correspond to observed data.

### Strength of handedness

MABSHI showed a stronger phylogenetic signal (h^2^ = 0.66) than MHI, though lower than previously reported ^14^. Again, this difference likely stems from our meta-analytic and multilevel approach, which allowed us to include multiple measurements per species and apply differential weighting based on study standard errors. The mean MABSHI for anthropoids was significantly above zero (MABSHI = 0.66, 95% CI: 0.44, 0.91; Supplementary Figure 2), indicating strong individual hand preferences across the clade, despite a lack of directional bias in most species. As expected, *H. sapiens* showed one of the highest MABSHI values (0.94), though East Javan langurs (*Trachypithecus auratus)* exhibited the strongest handedness strength among anthropoids (MABSHI = 0.98). Overall, humans and primarily arboreal species, such as langurs (*Trachypithecus)* and spider monkeys (*Ateles)*, displayed the highest MABSHI values, while predominantly terrestrial species, such as geladas (*Theropithecus)* and baboons (*Papio)*, showed weaker hand preferences (Supplementary Figure 2).

Similar to MHI, testing all hypotheses listed in Supplementary Table 1 with MABSHI as the dependent variable revealed no hypothesis performed significantly better than others, regardless of whether humans were included (See Supplementary Section 2; Supplementary Table 3). The exclusion of humans did, however, alter the significance of some predictors. As with MHI, predictors such as ECV and IMI were significant when humans were included, though for MABSHI, BM was highlighted as significant across multiple hypotheses. When we excluded humans from our dataset, we observed that most of the tested hypotheses showed SUB (substrate: terrestrial) as the only ‘significant’ predictor. The only other covariate that was ‘significant’ corresponded to DIM (i.e., body mass sexual dimorphism), which did not cross zero in the FH.

To further assess the relevance of the predictors that were found to be ‘‘significant’ for MABSHI, we ran two new models that exclusively considered all the predictors that were found to be relevant in all the tested hypotheses. The same modelling steps used in the previous models were followed. Therefore, we tested one model excluding *H. sapiens* that had as covariates SUBS and DIM, and another including humans that had as fixed effects DIET, ECV, IMI, SUBS, BM and DIM, and applied an iterative model reduction approach to only keep those predictors that were ‘significant’. SUBS (substrate: terrestrial) was the only predictors that was ‘significant’ in the first model without *H. sapiens* (Fig. 3c). The R^2^ value for this model was 0.79, which indicates again the crucial role that locomotor differences seem to play in MABSHI patterns. The second model that included *H. sapiens*, resulted in only three ‘significant’ predictors after model reduction, ECV, IMI and SUBS (substrate: terrestrial) (Fig. 3d). The R^2^ for this model was 0.8, which indicates the key role that locomotion and brain size plays in MABSHI pattern.

As with MHI, we explicitly tested the status of *H. sapiens* as an evolutionary singularity for MABSHI using a phylogenetic outlier test (see Methods). Figure 4c shows both the observed MABSHI value for *H. sapiens* and the posterior distribution of MABSHI values predicted by a model in which humans were excluded. The phylogenetic prediction for MABSHI is 0.43, whereas the observed value is 0.94, further highlighting the exceptional nature of human MABSHI relative to phylogenetic expectations. However, when IMI is included in the model, *H. sapiens* is no longer identified as an outlier (MABSHI: 0.86; Fig. 4d). This suggests that IMI is a critical factor in explaining the distinctiveness of human handedness as captured by MABSHI.

Using the reduced model to predict hominin MABSHI values, we observed consistently high handedness strength across species: *Ar. ramidu*s (0.73), *Au. afarensis* (0.64), *H. ergaster (*0.74), *H. erectus (*0.76), *H. floresiensis* (0.61), and *H. neanderthalensis* (0.81) (Fig. 5). These results suggest that handedness strength has remained consistently high since the last common ancestor of hominins and panins.

## Discussion

We present a comprehensive phylogenetic comparative meta-analysis of handedness in anthropoids that accounts for shared evolutionary history, effect variation across multiple sources, and potential publication bias, thus addressing previous limitations. By testing multiple hypotheses about the evolution of handedness simultaneously, we offer new insights into the role of evolutionary selection in shaping both the direction (MHI) and strength (MABSHI) of manual lateralisation.

Our results highlight significant phylogenetic signal in both MHI and MABSHI, indicating that handedness traits have evolved with descent along the branches of the primate phylogenetic tree. However, the data do not fully support any single existing hypothesis, with most predictors only showing significance when *H. sapiens* is included. This suggests that many hypotheses regarding handedness may be overly anthropocentric. As such, future research should seek to clearly distinguish between human-specific and broader primate explanations for manual lateralization.

A striking finding of this study is the identification of *H. sapiens* as a significant outlier in MABSHI but especially MHI. Humans display a pronounced right-handed bias (MHI = 0.76), which contrasts sharply with the phylogenetic prediction of from the reduced model excluding humans (MHI = 0.0) (Fig. 4a). Likewise, humans show extreme handedness strength (MABSHI = 0.94), with MABSHI values near the highest observed among anthropoids, although some arboreal species, such as spider monkeys and langurs, also exhibit high levels of lateralization. These results strongly support the hypothesis that human exceptionalism in handedness is likely owing to strong, human-specific selective pressures.

When considering the broader primate phylogeny, significant predictors for both MHI and MABSHI include factors such as IMI, ECV, DIET, IMI, and SOC SYS. However, when *H. sapiens* is excluded from the analysis, only a few predictors remain significant. Notably, IMI, substrate use (SUBS) emerged as key factor. Lower IMI (indicative of more quadrupedal or leaping locomotion) correlate with leftward bias in MHI, and arboreal species generally exhibit higher MABSHI values. This highlights the importance of locomotor and ecological factors in shaping handedness across non-human primates, whereas our large brain size emerges as an important link to human’s strong directionality in handedness. Taken together these results imply that our unusual gait was the main initial driver of our exceptional handedness with our large brain more linked to the directionality. In humans, the evolution of bipedalism and the subsequent freeing of the hands may have intensified selective pressures for stronger hand preferences. This finding aligns with previous research linking bipedal posture to the evolution of more pronounced handedness in fossil hominins, further suggesting that the evolutionary trajectory of human handedness is rooted in our unique locomotor adaptations ^27–30^.

Our predictions for other hominin species using our reduced models provide a more precise temporal perspective on the evolution of handedness direction and strength in human evolution (Fig. 5). Our results indicate consistently high handedness strength (MABSHI) among hominins from early on in the lineage. Although hominins became increasingly terrestrial throughout their evolution ^31^, a locomotor behaviour that reduces MABSHI among other anthropoids, the persistence of strong handedness strength is likely explained by our unique mode of locomotion -bipedalism-which freed the upper limbs and enabled specialised manual behaviours ^32^, maintaining selective pressure for lateralised hand use. This consistently high MABSHI level may also offer some support for those views that consider that that adaptations for bipedalism arose in an arboreal context ^33^, as more arboreal species tend to show higher handedness strength values.

In contrast, handedness direction (MHI) follows a different evolutionary pattern (Fig. 5). Early hominins such as *Ar. ramidus* and *Au. afarensis* exhibit MHI values relatively similar to other great apes. It is with the emergence of the genus *Homo*, and particularly the onset of significant encephalization, that we observe a marked increase in MHI values, reaching levels that are highly unusual among anthropoids. This pattern suggests a novel link between the evolution of directional handedness and increasing brain size ^34^, highlighting a link between these two human evolutionary hallmarks. In this context, the emergence of pronounced right-handedness bias in humans may be viewed as part of a broader suite of neurological and behavioural specialisations tied to the unique cognitive trajectory of our lineage. Importantly, this connection between encephalization and handedness direction has not been demonstrated in such a temporally resolved, phylogenetically informed framework before, offering new insights into the deep evolutionary roots of human lateralization.

The intriguing result for *H. floresiensis*, which shows relatively low MHI despite its placement within the genus *Homo*, may be explained by its unusual combination of small brain size and a unique locomotor repertoire blending bipedalism with arboreality. While the pelvis and lower limbs of *H. floresiensis* exhibit clear adaptations for upright walking, features such as long feet, an elongated forefoot, and curved phalanges suggest a locomotor pattern more similar to *Australopithecus*, including climbing behaviours ^35^. This unique morphology may have reduced handedness direction evolution in this lineage. However, further analyses are necessary to confirm this hypothesis.

While our analysis provides substantial evidence for the role of both ecological and anatomical factors in the evolution of handedness, it also raises important questions for future research. Notably, our findings underscore the need to refine hypotheses that distinguish human-specific factors from broader primate trends in terms of handedness. Additionally, expanding this analysis to include non-primate taxa, such as parrots or kangaroos, would be valuable in investigating the potential for convergent evolution in handedness across species ^36–38^. Finally, expanding the fossil sample size, by e.g., using other handedness proxies such as dental wear analyses, could significantly enhance our understanding of handedness evolution in extinct hominins and contribute to more refined and robust phylogenetic models.

In conclusion, this study highlights the complexity of handedness evolution and demonstrates the importance of considering both ecological and anatomical factors in understanding the selective pressures that have shaped human lateralization. Future research will benefit from a more comprehensive approach that considers both the primate and broader animal phylogenies, as well as the distinct evolutionary trajectory of *H. sapiens*.

## Materials and Methods

We combined the two most recent and comprehensive datasets available for primate handedness to carry out our subsequent analyses (Supplementary Data 1). ^21^ corresponds to a meta-analysis of hand preferences in coordinated bimanual tasks in non-human primates that compiled multiple sources on non-human handedness, whilst ^14^ consists of a phylogenetic comparative analysis that combined both new experimental data, as well as published sources on hand preference. To enable meaningful comparisons, we followed ^14^ and only considered studies that used the so-called ‘tube task’, as this experimental paradigm corresponds to a simple and widely applicable test to determine primate hand preferences. We only included data on MHI and MABSHI, as these two variables captured the main aspects of our phenomena of interest, namely the direction and strength of the manual asymmetries. The MHI is calculated by averaging the handedness scores of all individuals in a group, where handedness scores range from -1 (indicating complete left-handedness) to +1 (indicating complete right-handedness), with 0 representing ambidexterity. The MHI provides insight into the overall direction of hand preference within a population or sample. The MABSHI quantifies the strength of hand preference in a group, regardless of the direction of the preference (i.e., left, or right). It is calculated by taking the absolute value of the handedness scores of all individuals in the group, then averaging these absolute values. This is useful for determining the consistency or strength of hand preference within a population, irrespective of whether the preference is for the left or right hand. Therefore, we extracted all the information available for these two variables (i.e., effect sizes, variance measures and sample sizes) from ^21^ and ^14^ being careful of not duplicating sources that were referenced in both studies. The final dataset included information from 2,025 individuals across 41 species of anthropoid primates. It is important to bear in mind that in the present study we exclusively focus on the evolutionary underpinnings of population-level hand preferences, as opposed to studies or analyses that have focused on the individual-level of variation within species.

We downloaded 100 trees from http://vertlife.org/ comprising the 41 anthropoid species for which we have data to be used in throughout our analyses. These phylogenies correspond to Bayesian-inferred trees built using a ‘backbone-and-patch’ approach based on a 31-gene super-matrix ^39^. These phylogenies were used in our subsequent comparative analyses. To assess the hypotheses proposed to explain primate and/or human hand preference (Supplementary table 1) we compiled diverse relevant covariates from the literature. These variables are BM: body mass [kg]; ECV: endocranial volume [cm^3^]; FRUIT: percentage of fruits in diet; DIET: percentage of fruits and animals in diet; IMI: intermembral index; DIM: body mass sexual dimorphism; TOOL: tool use [0=absence, 1=presence); SUBS: substrate preference [0=arboreal, 1=both, 2=terrestrial]; SOC-SYS: social system [0=solitary, 1=pair, 2=group]; CL: intra-sexual competition levels sensu ^40^ [1, 2,3, 4]; EXT: extractive foraging [0=absence, 1=presence]; social learning sensu ^41^ [0=absence, 1=presence] (Fig.1; Table 1; Supplementary Data 1). In the case of intermembral index (IMI) and body mass sexual dimorphism (DIM), there were a limited number of species for which we did not find adequate data in the literature (See Supplementary Data 1 for more details). Therefore, we applied a phylogenetic imputation procedure known as ‘PhyloPars’ to deal with this missing data. Phenotypic covariance was assumed to be equivalent among species, and we also assumed Brownian motion in our imputation procedure. We used the maximum credibility clade phylogeny from our 100 trees when carrying out this procedure. Missing observations were incorporated by maximizing the log-likelihood of the covariance parameters using relevant covariates (i.e., BM for IMI and DIM), thus allowing us to predict means and covariances for missing values at the tips of the phylogenetic tree ^42^. This phylogenetic imputation procedure was carried out using the R package ‘Rphylopars’ v.0.3.9 ^43^. All the covariates that were considered in our subsequent modelling procedures were used as shown in Supplementary table 1 to test the 10 hypotheses proposed to explain handedness. Continuous predictors were logged_10_ and scaled prior to the modelling steps, while the few proportion variables were logit-transformed to ensure numerical stability.

All our Bayesian meta-analytical phylogenetic comparative analyses were done using the ‘brms’ package, which provides an R interface to fit Bayesian generalised linear and non-linear multivariate models using Stan ^44^. We applied generic weakly informative priors as all our continuous predictors were standardised (i.e., scaled, and centred). Continuous and categorical predictors were modelled using a normal distribution with mean 0 and variance 1. Prior predictive checks show that using wider priors (e.g., mean 0 and variance 5 or 10) did not influence our obtained results, and therefore narrower priors were preferred for computational efficiency. All predictors shown in Supplementary Table 1 were treated as fixed effects. Our random effects comprised non-phylogenetic species-specific effects (i.e., specific effects that would be independent of the phylogenetic relationship between species such as environmental or niche effects), as well as phylogenetic species-effects assumed to be sampled from a normal distribution with a mean of 0 and covariance matrix proportional to the phylogenetic correlation matrix among taxa obtained from our phylogenetic trees. Between-study heterogeneity standard deviation (τ) was explicitly modelled by via the group-level effects per study. Hence, we used (τ^2^), which corresponds the variance of true effects, as a measure of between-study heterogeneity. All random effects were modelled using half-Cauchy distributions with location 0 and scale 0.05, as our random-effect parameters should always be non-negative but can be close to zero ^45^. As our models were also meta-analytical, each dependent variable (i.e., MHI or MABSHI) had an associated measurement error defined by the standard error of every specific study. We considered an effect to be significant on handedness patterns when the model’s 95% credibility intervals of intercept and respective slopes did not overlap with zero. Phylogenetic signal was measured using Lynch’s *h*^*2*^, which is equivalent to Pagel’s λ in the context of phylogenetic generalised linear mixed models ^46,47^. We ran two independent chains per model using a warmup period of 4,000 out of a total of 12,000 iterations. Convergence was assessed both visually, by looking at the obtained trace plots, as well as by using rstan’s standard convergence and efficiency diagnostic metrics for Markov chains. The Gelman-Rubin 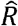 values were always one for all the estimated effects in all our models ^48^, thus confirming that our chains were well-mixed. Bulk and Tail Effective Sample Sizes were > 1,000 for all the effects estimated by all our models, hence indicating good sampling efficiency, as well as that all our estimates were reliable.

We repeated all our analyses separately for both variables, MHI and MABSHI, as different hypotheses may disparately apply to manual strength and/or direction asymmetry. In addition, we also repeated all our analyses excluding *H. sapiens*, to assess how our highly derived handedness patterns may influence the overall pattern observed across anthropoids. To account for topological differences, we repeated all our analyses using the 100 phylogenies previously mentioned. Values reported correspond to the average of the 100 models ran for each one of our analyses. We compared those models showing at least one ‘significant’ predictor using an efficient approximate leave-one-out cross-validation approach (LOO-CV) ^49^. Observations that were too influential in each model and that could not be accurately approximated were estimated using an exact cross-validation. To assess the fitness of our models we carried out posterior predictive checks by comparing 1,000 datasets simulated from each one of our models with our original data (i.e., MHI and MABSHI) (Supplementary section 1,2). All our models showed simulated datasets that resembled our original data, thus indicating the good fit of our models. We also ran four additional models (i.e., two for MABSHI ad two for MHI, including and excluding *H. sapiens)* that used as fixed effects all the predictors that were found to be ‘significant’ at least in one of the respective tested hypotheses. The idea was to assess what covariates played a key role in the observed handedness patterns. To evaluate the contribution of these covariates on the variance of MHI and MABSHI, we used an R^2^ metric that is suitable for Bayesian multilevel models ^50^.

To assess *H. sapiens’ a*pparent singularity, we used our intercept-only models to predict the expected levels MHI and MABSHI for our own species using a ‘phylogenetic outlier test’ ^26^, while also considering a meta-analytical component. We computed a posterior distribution of predictions for the dependent variables (i.e., MHI or MABSHI) for a new observation (i.e., in our case *H. sapiens)* given the posterior distribution of intercept-only models, the standard error for our species, and the phylogenetic effects. Predictive distributions that deviate strongly from the known value (i.e., outliers) provide evidence that the species has undergone a substantial amount of evolutionary change which cannot be accounted for by its phylogenetic position, branch lengths, and evolutionary change in the independent variable. The implication is that the trait has adaptive value for the species in ways not shared by its close relatives. This test was used to evaluate the idea that human handedness levels correspond to an evolutionary singularity (i.e., a derived character unique to us). To further explore the role of our eco-evolutionary predictors, we repeated the above process using the most supported models for MHI and MABSHI. This allowed us to assess the relative contribution of the covariates present in the most supported model when explaining *H. sapiens* handedness singularity.

To predict hominin MHI and MABSHI values, we used the reduced human models to generate predictions for hominins, while marginalising over measurement error and non-phylogenetic species effects. We assumed human-level standard errors for our meta-analytical component when computing our predictions. Data on hominin IMI was obtained from ^51^, whilst ECVs were obtained from ^34^. Hominins were categorised as ‘group’ in social system, whereas for substrate we classified *Ardipithecus* as ‘both’, whilst all the remaining hominin species were classified as ‘terrestrial’. We grafted the maximum credibility clade phylogeny from ^34^ to a consensus phylogeny obtained from the previously used 100 anthropoid trees. We removed the hominin species for which we did not have any data, which left us with the following species: *Ar. ramidus, Au. afarensis, H. ergaster, H. erectus, H. floresiensis* and *H. neanderthalensis*.

## Supporting information

Supplementary Section 1

Supplementary Section 2

## Acknowledgments

We are grateful to Elodie Freymann for her work with Figure 1 in this article. This work was funded by a Leverhulme Trust Research Leadership Award to CV (RL-2019-012).

