## Supplementary Figure 1 for "Phylogenetic meta-analysis implicates large brains and our unusual posture in human handedness"

| Reference | Species | Sample size | Mean [95% CI] |
|---|---|---|---|
| Caspar et al. 2022 | *Ateles fusciceps* | 37 | 0.16 [−0.10, 0.42] |
| Nelson & Boeving 2015 | *Ateles fusciceps* | 9 | −0.33 [−0.94, 0.28] |
| Nelson et al., 2015 | *Ateles fusciceps* | 10 | −0.12 [−0.60, 0.36] |
| Caspar et al. 2022 | *Ateles geoffroyi* | 9 | −0.00 [−0.57, 0.57] |
| Motes Rodrigo et al. 2018 | *Ateles geoffroyi* | 14 | 0.10 [−0.39, 0.59] |
| Caspar et al. 2022 | *Ateles hybridus* | 18 | −0.38 [−0.78, 0.03] |
| Caspar et al. 2022 | *Cercocebus torquatus* | 18 | −0.18 [−0.56, 0.20] |
| Maille et al. 2013 | *Cercocebus torquatus* | 13 | 0.18 [−0.15, 0.51] |
| Caspar et al. 2022 | *Cercopithecus diana* | 20 | 0.18 [−0.18, 0.53] |
| Caspar et al. 2022 | *Cercopithecus neglectus* | 12 | −0.04 [−0.41, 0.32] |
| Maille et al. 2013 | *Cercopithecus neglectus* | 12 | −0.44 [−0.81, −0.08] |
| Schweitzer et al., 2007 | *Cercopithecus neglectus* | 12 | −0.42 [−0.78, −0.06] |
| Hopkins et al., 2003 | *Gorilla gorilla* | 31 | 0.10 [−0.11, 0.31] |
| Hopkins et al., 2011 | *Gorilla gorilla* | 76 | 0.25 [ 0.12, 0.38] |
| Cochet & Vauclair, 2012 | *Homo sapiens* | 127 | 0.76 [ 0.66, 0.86] |
| Caspar et al. 2022 | *Hylobates lar* | 16 | −0.06 [−0.42, 0.30] |
| Caspar et al., 2018 | *Hylobates lar* | 3 | 0.13 [−0.72, 0.98] |
| Morino et al. 2017 | *Hylobates lar* | 6 | 0.13 [−0.39, 0.65] |
| Spoelstra 2021 | *Hylobates lar* | 11 | −0.06 [−0.47, 0.35] |
| Caspar et al. 2022 | *Hylobates moloch* | 22 | −0.12 [−0.48, 0.25] |
| Caspar et al. 2022 | *Leontopithecus chrysomelas* | 30 | 0.15 [−0.06, 0.36] |
| Caspar et al. 2022 | *Leontopithecus chrysopygus* | 15 | 0.04 [−0.19, 0.27] |
| Caspar et al. 2022 | *Leontopithecus rosalia* | 28 | 0.02 [−0.20, 0.25] |
| Caspar et al. 2022 | *Macaca fascicularis* | 12 | 0.02 [−0.39, 0.44] |
| Chatagny et al., 2013 | *Macaca fascicularis* | 8 | −0.12 [−0.71, 0.46] |
| Zhao et al., 2016 | *Macaca leonina* | 9 | −0.21 [−0.56, 0.14] |
| Nelson et al., 2011 | *Macaca mulatta* | 16 | −0.02 [−0.30, 0.26] |
| Caspar et al. 2022 | *Macaca nemestrina* | 29 | 0.04 [−0.20, 0.27] |
| Caspar et al. 2022 | *Macaca silenus* | 35 | −0.05 [−0.24, 0.13] |
| Caspar et al. 2022 | *Macaca sylvanus* | 15 | −0.14 [−0.49, 0.21] |
| Regaiolli et al., 2018 | *Macaca sylvanus* | 6 | 0.42 [−0.20, 1.04] |
| Schmitt et al., 2008 | *Macaca sylvanus* | 3 | −0.33 [−1.26, 0.60] |
| Canteloup et al., 2013 | *Macaca tonkeana* | 14 | −0.06 [−0.41, 0.29] |
| Caspar et al. 2022 | *Mandrillus sphinx* | 32 | 0.03 [−0.14, 0.21] |
| Caspar et al. 2022 | *Nomascus gabriellae* | 6 | −0.31 [−0.74, 0.12] |
| Caspar et al., 2018 | *Nomascus gabriellae* | 4 | 0.03 [−0.94, 1.00] |
| Caspar et al. 2022 | *Nomascus leucogenys* | 7 | 0.05 [−0.51, 0.61] |
| Caspar et al., 2018 | *Nomascus leucogenys* | 9 | −0.21 [−0.68, 0.27] |
| Fan et al., 2017 | *Nomascus leucogenys* | 9 | −0.03 [−0.39, 0.33] |
| Caspar et al., 2018 | *Nomascus siki* | 4 | −0.24 [−1.15, 0.67] |
| Chapelain & Hogervorst, 2009 | *Pan paniscus* | 29 | 0.05 [−0.19, 0.29] |
| Hopkins et al., 2011 | *Pan paniscus* | 118 | 0.04 [−0.07, 0.15] |
| Hopkins et al., 2011 | *Pan troglodytes* | 536 | 0.13 [ 0.08, 0.18] |
| Llorente et al., 2009 | *Pan troglodytes* | 14 | 0.37 [−0.05, 0.79] |
| Padrell et al., 2019 | *Pan troglodytes* | 14 | 0.35 [−0.06, 0.76] |
| Phillips & Hopkins, 2007 | *Pan troglodytes* | 16 | 0.11 [−0.11, 0.33] |
| Vauclair et al. 2005 | *Papio anubis* | 84 | 0.11 [−0.02, 0.24] |
| Caspar et al. 2022 | *Papio hamadryas* | 24 | 0.07 [−0.14, 0.27] |
| Caspar et al. 2022 | *Pithecia pithecia* | 7 | −0.39 [−1.07, 0.30] |
| Hopkins et al., 2003 | *Pongo* sp. | 19 | −0.42 [−0.77, −0.07] |
| Hopkins et al., 2011 | *Pongo* sp. | 47 | −0.22 [−0.38, −0.07] |
| Cubi & Llorente, 2021 | *Pygathrix cinerea* | 18 | 0.17 [−0.12, 0.45] |
| Zhao et al., 2012 | *Rhinopithecus roxellana* | 24 | −0.32 [−0.61, −0.03] |
| Meguerditchian et al., 2012 | *Saimiri sciureus* | 36 | −0.12 [−0.38, 0.14] |
| de Andrade & de Sousa, 2018 | *Sapajus apella* | 12 | −0.17 [−0.57, 0.23] |
| Lilak & Phillips, 2008 | *Sapajus apella* | 11 | −0.10 [−0.53, 0.33] |
| Phillips & Hopkins, 2007 | *Sapajus apella* | 11 | −0.02 [−0.52, 0.48] |
| Phillips & Sherwood, 2005 | *Sapajus apella* | 7 | −0.07 [−0.68, 0.54] |
| Phillips & Sherwood, 2007 | *Sapajus apella* | 13 | 0.10 [−0.35, 0.55] |
| Phillips, Sherwood & Lilak, 2007 | *Sapajus apella* | 13 | 0.10 [−0.35, 0.55] |
| Spinozzi et al., 1998 | *Sapajus apella* | 26 | 0.33 [ 0.03, 0.63] |
| Caspar et al. 2022 | *Sapajus flavius* | 3 | −0.65 [−1.34, 0.04] |
| de Andrade & de Sousa, 2018 | *Sapajus flavius* | 18 | −0.04 [−0.45, 0.36] |
| Caspar et al. 2022 | *Sapajus xanthosternos* | 16 | 0.05 [−0.31, 0.41] |
| de Andrade & de Sousa, 2018 | *Sapajus xanthosternos* | 18 | 0.13 [−0.23, 0.48] |
| Caspar et al. 2022 | *Semnopithecus entellus* | 30 | −0.18 [−0.40, 0.03] |
| Caspar et al. 2022 | *Symphalangus syndactylus* | 14 | 0.09 [−0.26, 0.45] |
| Morino et al. 2017 | *Symphalangus syndactylus* | 16 | −0.23 [−0.46, −0.01] |
| Caspar et al. 2022 | *Theropithecus gelada* | 38 | 0.05 [−0.05, 0.16] |
| Caspar et al. 2022 | *Trachypithecus auratus* | 8 | −0.26 [−0.96, 0.45] |
| Cubi & Llorente, 2021 | *Trachypithecus hatinhensis* | 18 | −0.25 [−0.63, 0.13] |
| | | | **−0.03 [−0.31, 0.22]** |

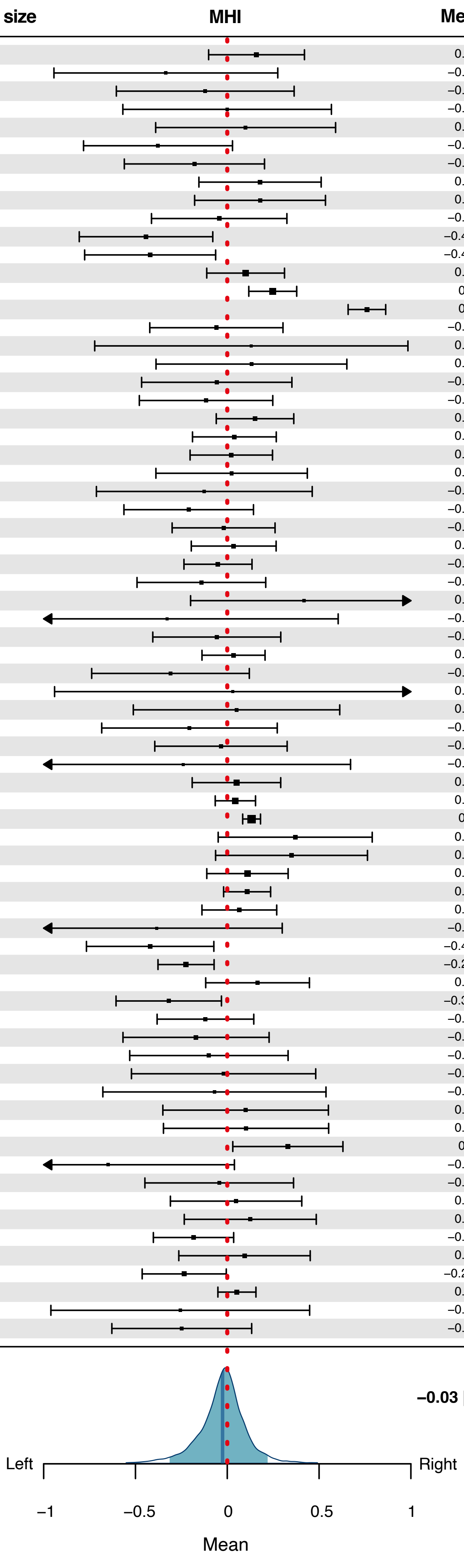
