## Supplementary Figure 2 for "Phylogenetic meta-analysis implicates large brains and our unusual posture in human handedness"

| Reference | Species | Sample size | Mean [95% CI] |
|---|---|---|---|
| Caspar et al. 2022 | *Ateles fusciceps* | 37 | 0.76 [0.67, 0.86] |
| Nelson & Boeving 2015 | *Ateles fusciceps* | 9 | 0.94 [0.87, 1.00] |
| Nelson et al., 2015 | *Ateles fusciceps* | 10 | 0.70 [0.51, 0.89] |
| Caspar et al. 2022 | *Ateles geoffroyi* | 9 | 0.73 [0.47, 0.99] |
| Motes Rodrigo et al. 2018 | *Ateles geoffroyi* | 14 | 0.89 [0.81, 0.98] |
| Caspar et al. 2022 | *Ateles hybridus* | 18 | 0.92 [0.84, 1.00] |
| Caspar et al. 2022 | *Cercocebus torquatus* | 18 | 0.79 [0.68, 0.90] |
| Maille et al. 2013 | *Cercocebus torquatus* | 13 | 0.49 [0.28, 0.70] |
| Caspar et al. 2022 | *Cercopithecus diana* | 20 | 0.75 [0.62, 0.89] |
| Caspar et al. 2022 | *Cercopithecus neglectus* | 12 | 0.52 [0.32, 0.73] |
| Maille et al. 2013 | *Cercopithecus neglectus* | 12 | 0.72 [0.58, 0.86] |
| Schweitzer et al., 2007 | *Cercopithecus neglectus* | 12 | 0.70 [0.56, 0.84] |
| Hopkins et al., 2003 | *Gorilla gorilla* | 31 | 0.50 [0.38, 0.62] |
| Hopkins et al., 2011 | *Gorilla gorilla* | 76 | 0.54 [0.47, 0.61] |
| Cochet & Vauclair, 2012 | *Homo sapiens* | 127 | 0.94 [0.91, 0.97] |
| Caspar et al. 2022 | *Hylobates lar* | 16 | 0.64 [0.46, 0.81] |
| Caspar et al., 2018 | *Hylobates lar* | 3 | 0.54 [0.08, 0.99] |
| Morino et al. 2017 | *Hylobates lar* | 6 | 0.59 [0.46, 0.72] |
| Spoelstra 2021 | *Hylobates lar* | 11 | 0.62 [0.48, 0.76] |
| Caspar et al. 2022 | *Hylobates moloch* | 22 | 0.80 [0.67, 0.93] |
| Caspar et al. 2022 | *Leontopithecus chrysomelas* | 30 | 0.51 [0.40, 0.63] |
| Caspar et al. 2022 | *Leontopithecus chrysopygus* | 15 | 0.35 [0.21, 0.49] |
| Caspar et al. 2022 | *Leontopithecus rosalia* | 28 | 0.50 [0.38, 0.62] |
| Caspar et al. 2022 | *Macaca fascicularis* | 12 | 0.64 [0.48, 0.81] |
| Chatagny et al., 2013 | *Macaca fascicularis* | 8 | 0.75 [0.54, 0.96] |
| Zhao et al., 2016 | *Macaca leonina* | 9 | 0.49 [0.31, 0.67] |
| Nelson et al., 2011 | *Macaca mulatta* | 16 | 0.45 [0.29, 0.61] |
| Caspar et al. 2022 | *Macaca nemestrina* | 29 | 0.53 [0.40, 0.65] |
| Caspar et al. 2022 | *Macaca silenus* | 35 | 0.47 [0.37, 0.57] |
| Caspar et al. 2022 | *Macaca sylvanus* | 15 | 0.63 [0.48, 0.77] |
| Regaiolli et al., 2018 | *Macaca sylvanus* | 6 | 0.76 [0.47, 1.04] |
| Schmitt et al., 2008 | *Macaca sylvanus* | 3 | 0.72 [0.45, 0.99] |
| Canteloup et al., 2013 | *Macaca tonkeana* | 14 | 0.54 [0.35, 0.73] |
| Caspar et al. 2022 | *Mandrillus sphinx* | 32 | 0.39 [0.28, 0.49] |
| Caspar et al. 2022 | *Nomascus gabriellae* | 6 | 0.47 [0.16, 0.77] |
| Caspar et al., 2018 | *Nomascus gabriellae* | 4 | 0.85 [0.71, 0.99] |
| Caspar et al. 2022 | *Nomascus leucogenys* | 7 | 0.62 [0.36, 0.89] |
| Caspar et al., 2018 | *Nomascus leucogenys* | 9 | 0.62 [0.36, 0.87] |
| Fan et al., 2017 | *Nomascus leucogenys* | 9 | 0.39 [0.16, 0.63] |
| Caspar et al., 2018 | *Nomascus siki* | 4 | 0.73 [0.27, 1.19] |
| Chapelain & Hogervorst, 2009 | *Pan paniscus* | 29 | 0.57 [0.46, 0.68] |
| Hopkins et al., 2011 | *Pan paniscus* | 118 | 0.53 [0.48, 0.58] |
| Hopkins et al., 2011 | *Pan troglodytes* | 536 | 0.51 [0.48, 0.53] |
| Llorente et al., 2009 | *Pan troglodytes* | 14 | 0.80 [0.68, 0.92] |
| Padrell et al., 2019 | *Pan troglodytes* | 14 | 0.81 [0.69, 0.93] |
| Phillips & Hopkins, 2007 | *Pan troglodytes* | 16 | 0.35 [0.21, 0.49] |
| Vauclair et al. 2005 | *Papio anubis* | 84 | 0.53 [0.46, 0.59] |
| Caspar et al. 2022 | *Papio hamadryas* | 24 | 0.41 [0.29, 0.53] |
| Caspar et al. 2022 | *Pithecia pithecia* | 7 | 0.93 [0.86, 1.00] |
| Hopkins et al., 2003 | *Pongo* sp. | 19 | 0.63 [0.36, 0.90] |
| Hopkins et al., 2011 | *Pongo* sp. | 47 | 0.49 [0.40, 0.57] |
| Cubi & Llorente, 2021 | *Pygathrix cinerea* | 18 | 0.50 [0.33, 0.67] |
| Zhao et al., 2012 | *Rhinopithecus roxellana* | 24 | 0.73 [0.63, 0.83] |
| Meguerditchian et al., 2012 | *Saimiri sciureus* | 36 | 0.76 [0.67, 0.85] |
| de Andrade & de Sousa, 2018 | *Sapajus apella* | 12 | 0.61 [0.41, 0.81] |
| Lilak & Phillips, 2008 | *Sapajus apella* | 11 | 0.67 [0.53, 0.81] |
| Phillips & Hopkins, 2007 | *Sapajus apella* | 11 | 0.75 [0.59, 0.91] |
| Phillips & Sherwood, 2005 | *Sapajus apella* | 7 | 0.70 [0.46, 0.94] |
| Phillips & Sherwood, 2007 | *Sapajus apella* | 13 | 0.76 [0.62, 0.90] |
| Phillips, Sherwood & Lilak, 2007 | *Sapajus apella* | 13 | 0.76 [0.62, 0.90] |
| Spinozzi et al., 1998 | *Sapajus apella* | 26 | 0.82 [0.75, 0.89] |
| Caspar et al. 2022 | *Sapajus flavius* | 3 | 0.68 [0.06, 1.30] |
| de Andrade & de Sousa, 2018 | *Sapajus flavius* | 18 | 0.78 [0.62, 0.94] |
| Caspar et al. 2022 | *Sapajus xanthosternos* | 16 | 0.64 [0.49, 0.79] |
| de Andrade & de Sousa, 2018 | *Sapajus xanthosternos* | 18 | 0.71 [0.57, 0.85] |
| Caspar et al. 2022 | *Semnopithecus entellus* | 30 | 0.56 [0.46, 0.66] |
| Caspar et al. 2022 | *Symphalangus syndactylus* | 14 | 0.57 [0.38, 0.76] |
| Morino et al. 2017 | *Symphalangus syndactylus* | 16 | 0.38 [0.21, 0.55] |
| Caspar et al. 2022 | *Theropithecus gelada* | 38 | 0.26 [0.19, 0.32] |
| Caspar et al. 2022 | *Trachypithecus auratus* | 8 | 0.98 [0.96, 1.01] |
| Cubi & Llorente, 2021 | *Trachypithecus hatinhensis* | 18 | 0.82 [0.73, 0.91] |
| | | | **0.67 [0.44, 0.91]** |

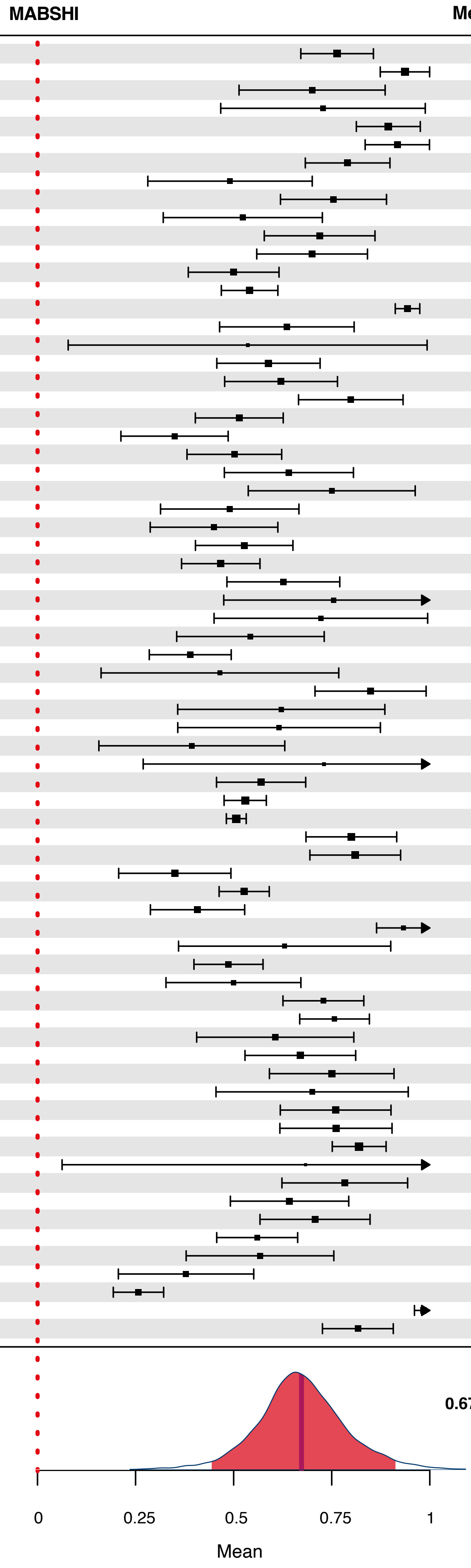
