## Supplementary Section 2 for "Phylogenetic meta-analysis implicates large brains and our unusual posture in human handedness"

**MABSHI results excluding *Homo sapiens***

Numerical results for all the hypotheses tested, using MABSHI as the dependent variable, are presented here (see Supplementary Table 1 for further details). *Homo sapiens* was excluded from these analyses. In addition, the final reduced model is also presented after all the hypotheses tested. Posterior predictive checks for each of the tested hypotheses are shown, along with plots illustrating the estimates for the fixed effects included in each of the tested models.

#### 0) Intercepts-only model

### Summary

| effect | component | term | estimate | std.error | conf.low | conf.high | rhat |
|---|---|---|---|---|---|---|---|
| fixed | cond | (Intercept) | 0.67 | 0.11 | 0.46 | 0.88 | 1 |

| effect | component | group | term | estimate | std.error | conf.low | conf.high | rhat |
|---|---|---|---|---|---|---|---|---|
| ran_pars | cond | obs | sd__(Intercept) | 0.10 | 0.02 | 0.07 | 0.14 | 1 |
| ran_pars | cond | phylo | sd__(Intercept) | 0.03 | 0.01 | 0.01 | 0.05 | 1 |
| ran_pars | cond | species | sd__(Intercept) | 0.04 | 0.03 | 0.00 | 0.10 | 1 |

**Posterior predictive check; 100 simulated datasets**

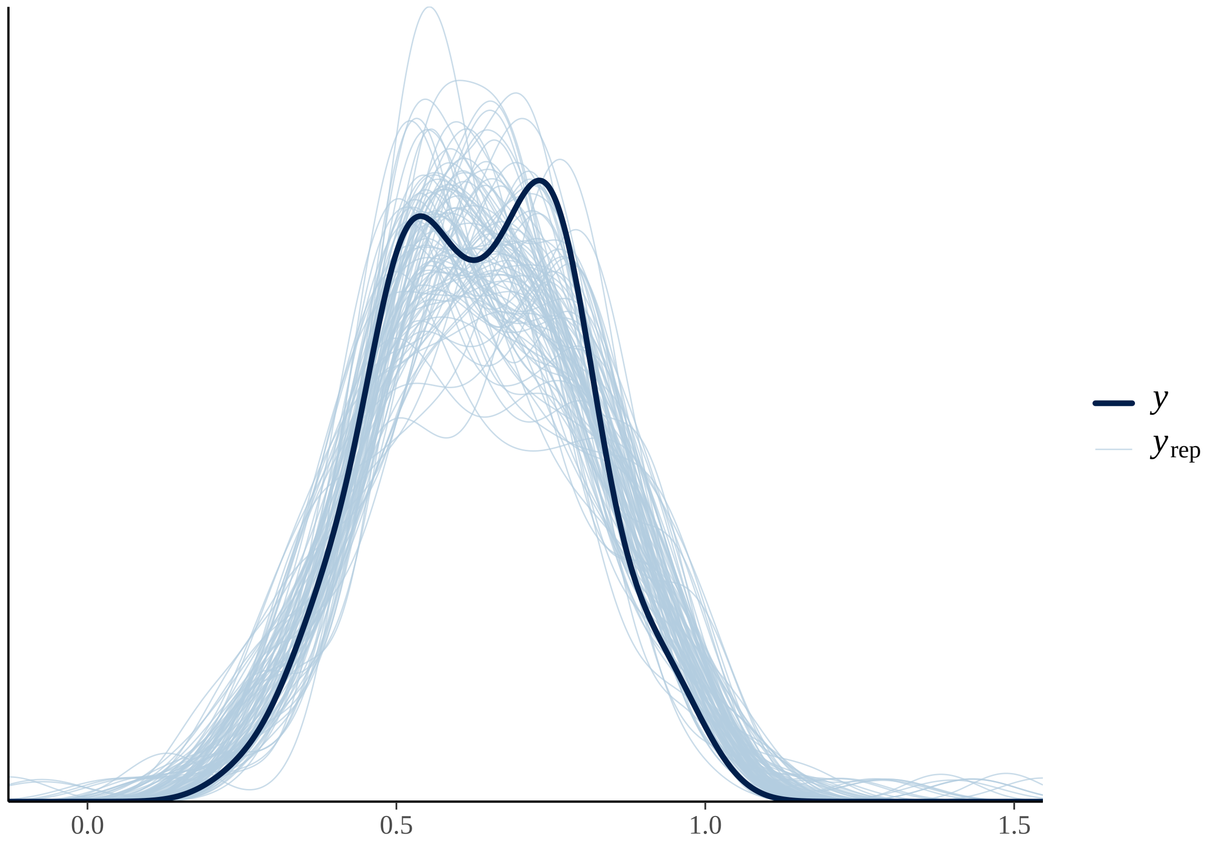

### Fixed effects results

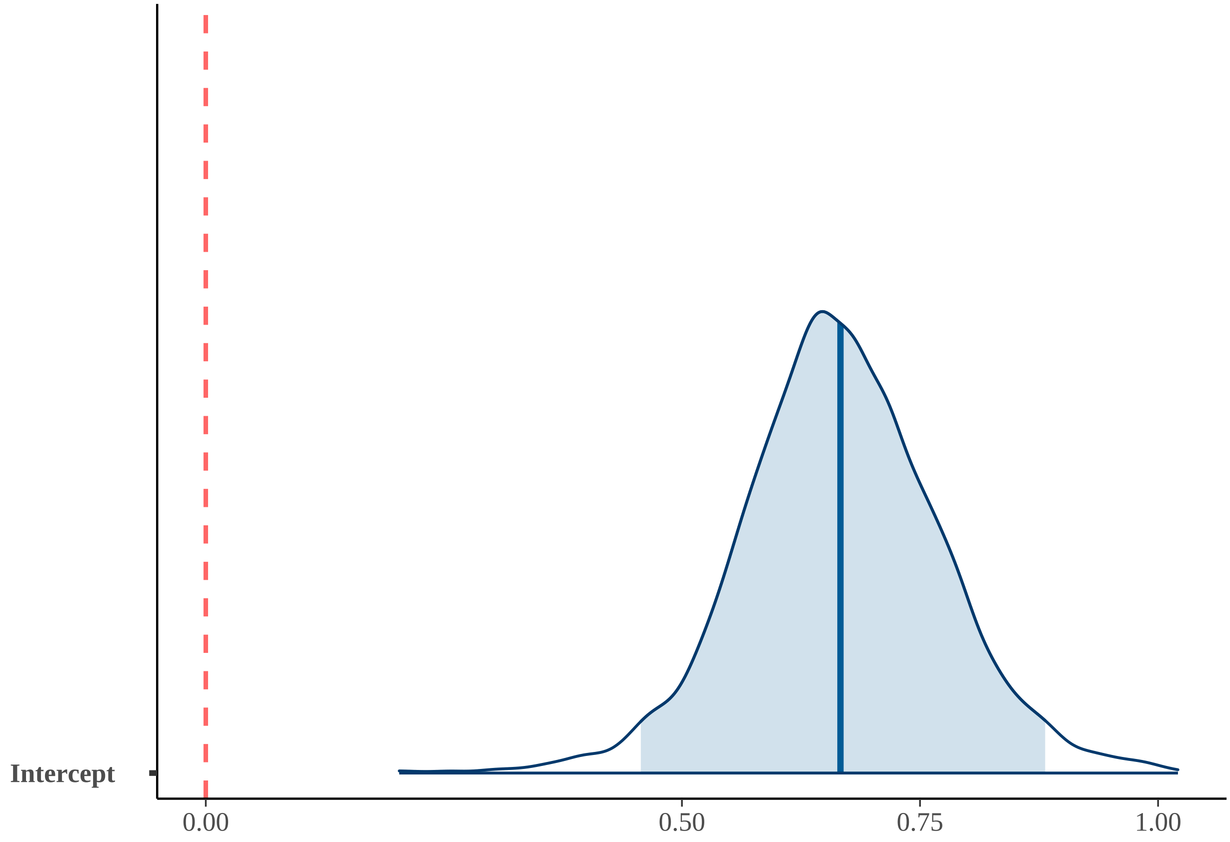


##### 1) POH1

#### Summary POH1

| effect | component | term | estimate | std.error | conf.low | conf.high | rhat |
|---|---|---|---|---|---|---|---|
| fixed | cond | (Intercept) | 0.68 | 0.10 | 0.48 | 0.88 | 1 |
| fixed | cond | fruit | 0.02 | 0.02 | -0.02 | 0.06 | 1 |
| fixed | cond | substrateBOTH | -0.02 | 0.06 | -0.14 | 0.11 | 1 |
| fixed | cond | substrateT | -0.23 | 0.10 | -0.42 | -0.04 | 1 |

| effect | component | group | term | estimate | std.error | conf.low | conf.high | rhat |
|---|---|---|---|---|---|---|---|---|
| ran_pars | cond | obs | sd___(Intercept) | 0.09 | 0.02 | 0.06 | 0.13 | 1 |
| ran_pars | cond | phylo | sd___(Intercept) | 0.03 | 0.01 | 0.01 | 0.05 | 1 |
| ran_pars | cond | species | sd___(Intercept) | 0.03 | 0.02 | 0.00 | 0.09 | 1 |

**Posterior predictive check POH1; 100 simulated datasets**

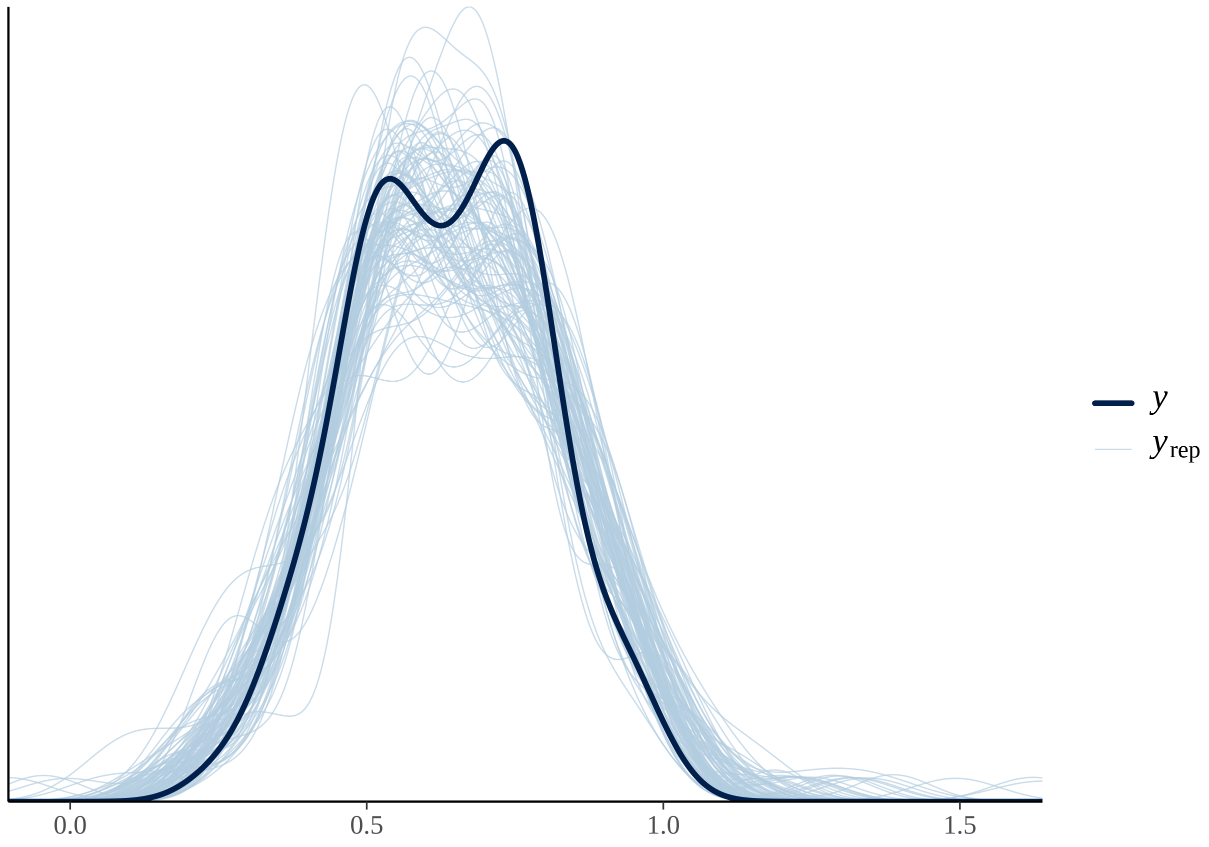

#### Fixed effects results POH1

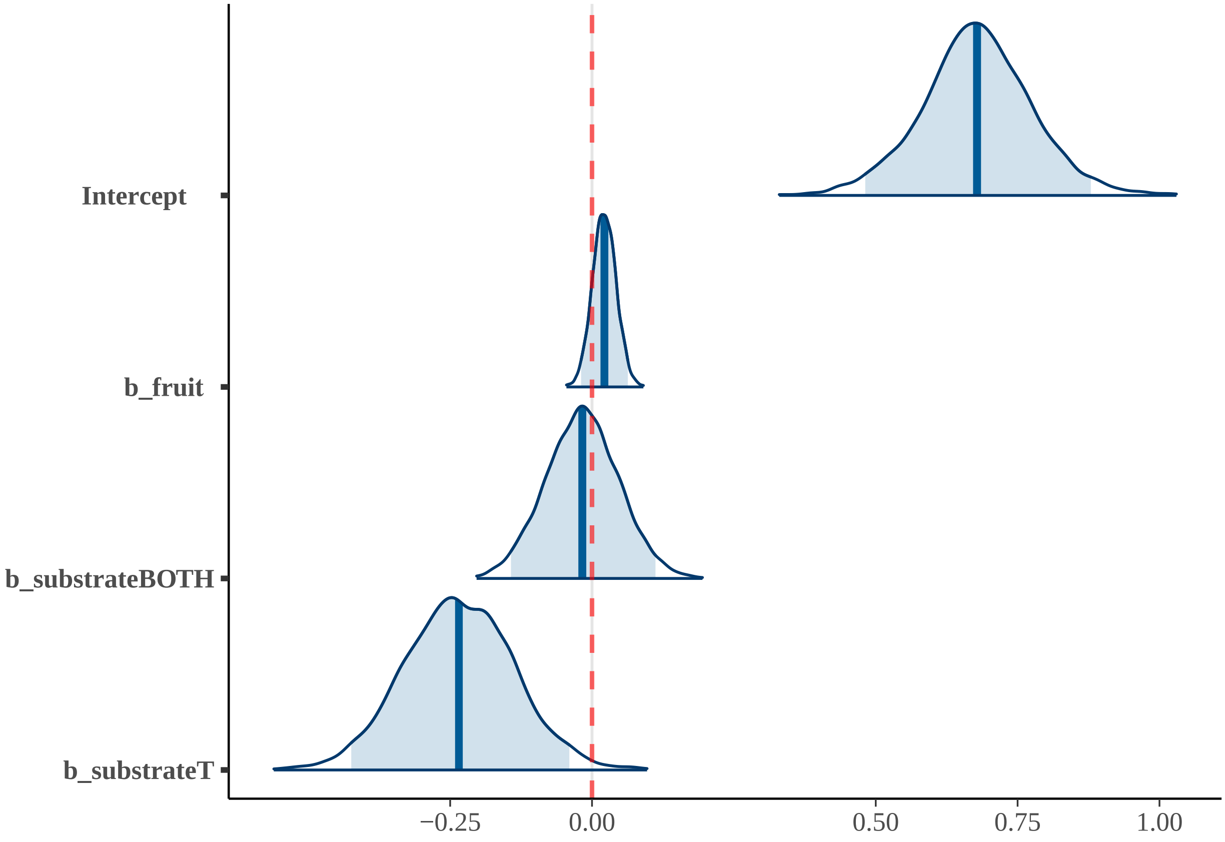


##### 2) POH2

#### Summary POH2

| effect | component | term | estimate | std.error | conf.low | conf.high | rhat |
|---|---|---|---|---|---|---|---|
| fixed | cond | (Intercept) | 0.67 | 0.10 | 0.46 | 0.88 | 1 |
| fixed | cond | diet | 0.01 | 0.02 | -0.03 | 0.06 | 1 |
| fixed | cond | substrateBOTH | -0.02 | 0.07 | -0.15 | 0.11 | 1 |
| fixed | cond | substrateT | -0.24 | 0.10 | -0.44 | -0.05 | 1 |

| effect | component | group | term | estimate | std.error | conf.low | conf.high | rhat |
|---|---|---|---|---|---|---|---|---|
| ran_pars | cond | obs | sd__(Intercept) | 0.09 | 0.02 | 0.06 | 0.13 | 1 |
| ran_pars | cond | phylo | sd__(Intercept) | 0.03 | 0.01 | 0.01 | 0.05 | 1 |
| ran_pars | cond | species | sd__(Intercept) | 0.03 | 0.02 | 0.00 | 0.09 | 1 |

**Posterior predictive check POH2; 100 simulated datasets**

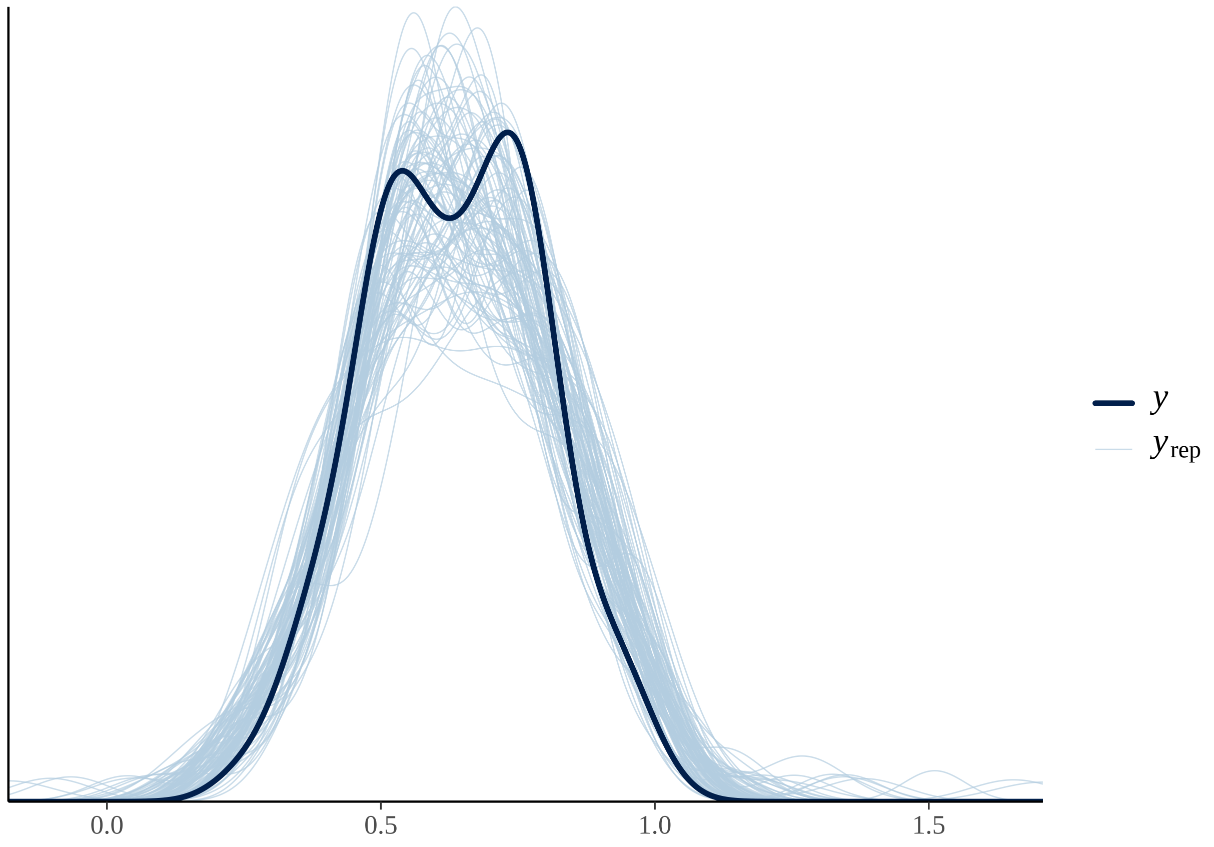

#### Fixed effects results POH2

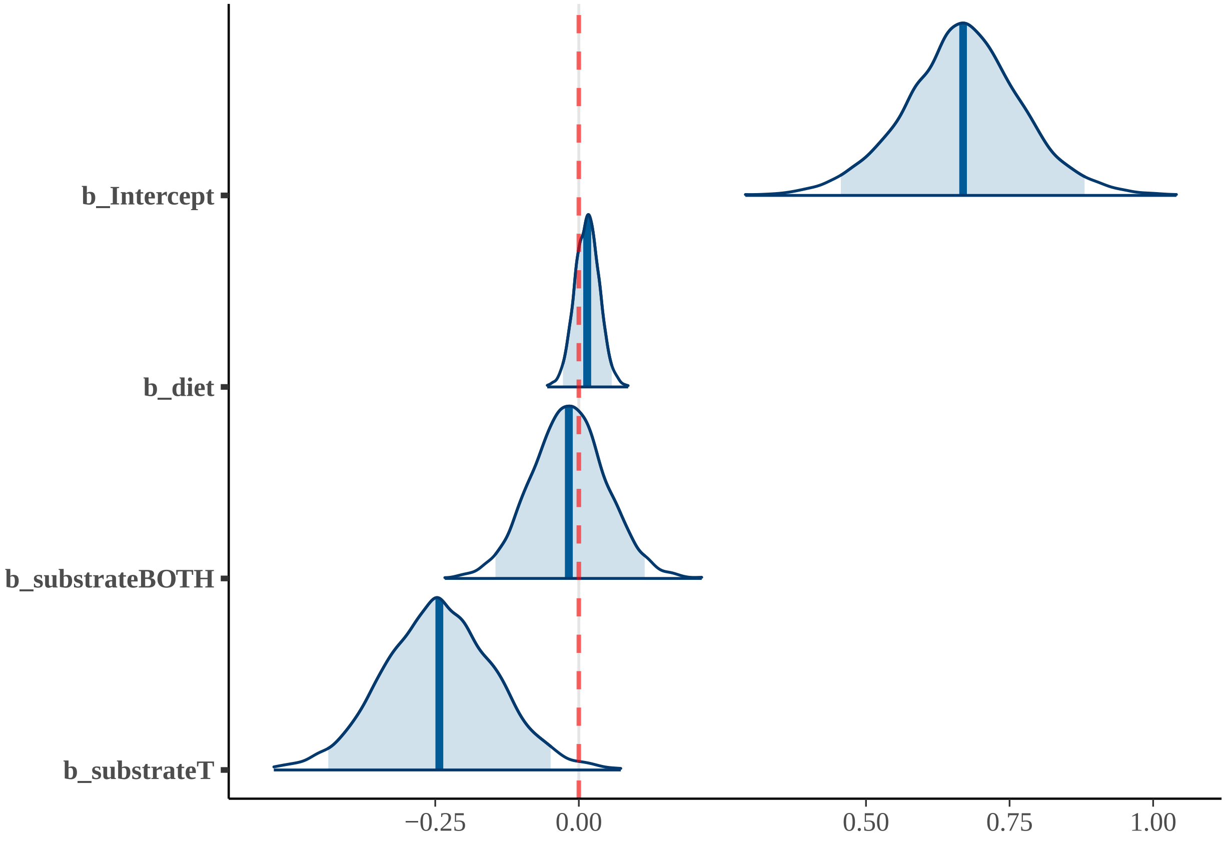


### 3) BH

#### Summary BH

| effect | component | term | estimate | std.error | conf.low | conf.high | rhat |
|---|---|---|---|---|---|---|---|
| fixed | cond | (Intercept) | 0.70 | 0.09 | 0.51 | 0.89 | 1 |
| fixed | cond | imi | -0.08 | 0.05 | -0.18 | 0.03 | 1 |
| fixed | cond | ecv | 0.08 | 0.10 | -0.11 | 0.27 | 1 |
| fixed | cond | bm | -0.05 | 0.10 | -0.24 | 0.14 | 1 |
| fixed | cond | substrateBOTH | -0.08 | 0.08 | -0.22 | 0.08 | 1 |
| fixed | cond | substrateT | -0.30 | 0.11 | -0.51 | -0.09 | 1 |

| effect | component | group | term | estimate | std.error | conf.low | conf.high | rhat |
|---|---|---|---|---|---|---|---|---|
| ran_pars | cond | obs | sd__(Intercept) | 0.10 | 0.02 | 0.06 | 0.13 | 1 |
| ran_pars | cond | phylo | sd__(Intercept) | 0.03 | 0.01 | 0.01 | 0.05 | 1 |
| ran_pars | cond | species | sd__(Intercept) | 0.03 | 0.03 | 0.00 | 0.10 | 1 |

**Posterior predictive check BH; 100 simulated datasets**

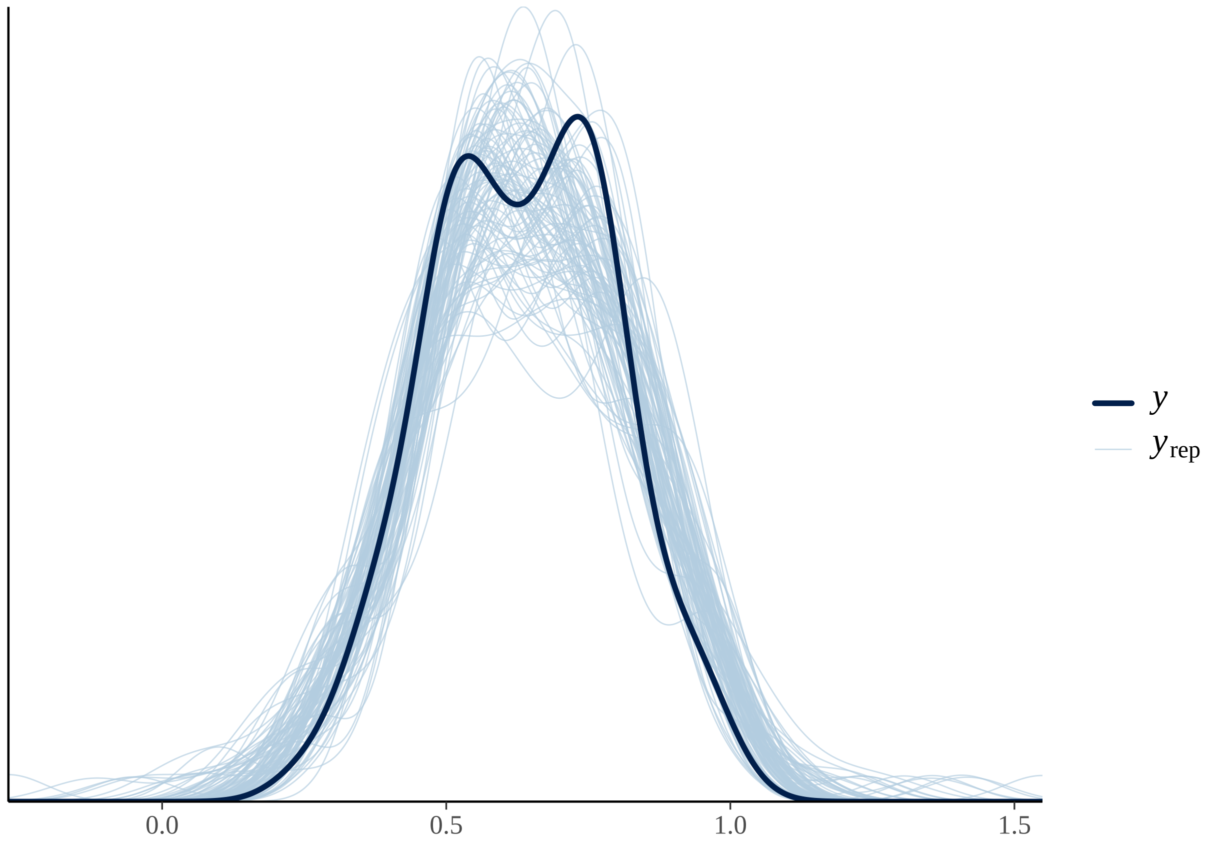

#### Fixed effects results BH

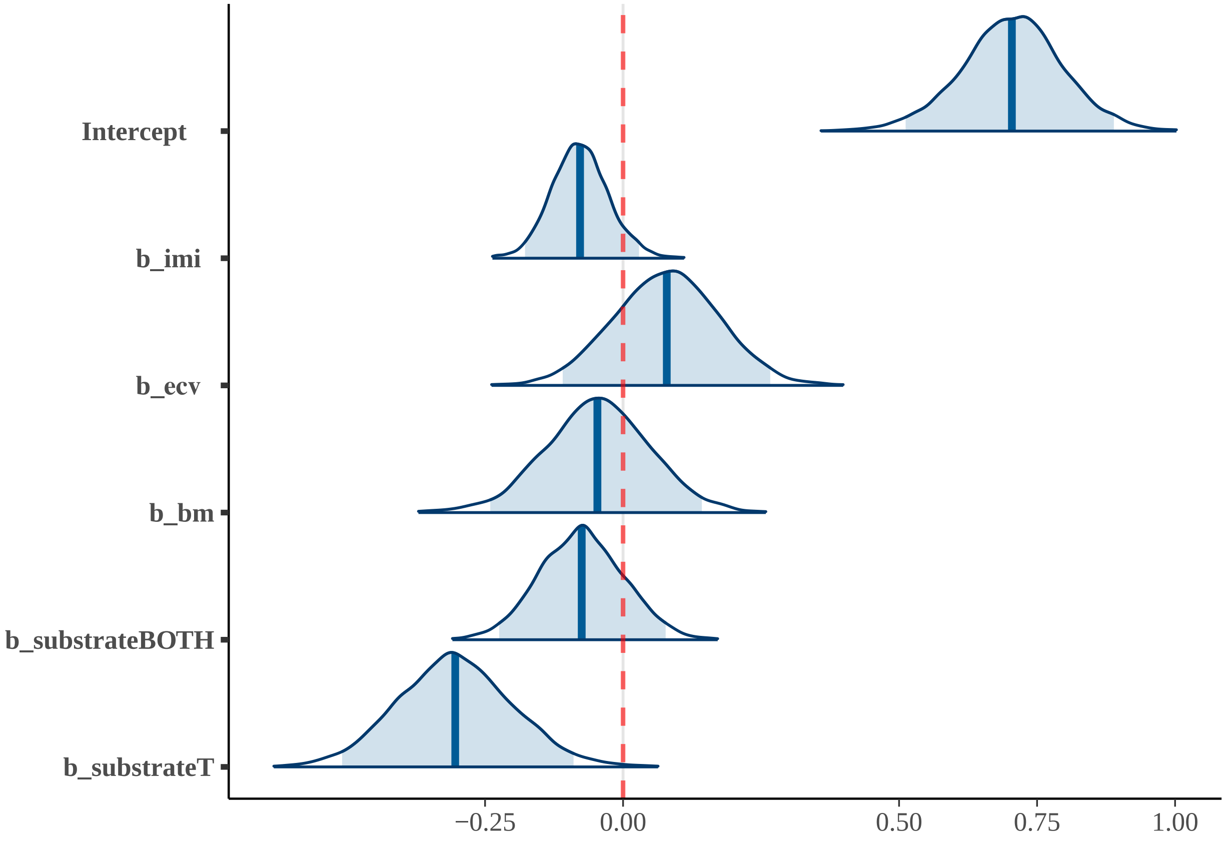


#### 4) TUH

#### Summary TUH

| effect | component | term | estimate | std.error | conf.low | conf.high | rhat |
|---|---|---|---|---|---|---|---|
| fixed | cond | (Intercept) | 0.65 | 0.12 | 0.41 | 0.90 | 1 |
| fixed | cond | tool_useYes | 0.06 | 0.08 | -0.09 | 0.21 | 1 |
| fixed | cond | ecv | 0.05 | 0.12 | -0.18 | 0.28 | 1 |
| fixed | cond | bm | -0.09 | 0.12 | -0.32 | 0.14 | 1 |

| effect | component | group | term | estimate | std.error | conf.low | conf.high | rhat |
|---|---|---|---|---|---|---|---|---|
| ran_pars | cond | obs | sd__(Intercept) | 0.10 | 0.02 | 0.06 | 0.14 | 1 |
| ran_pars | cond | phylo | sd__(Intercept) | 0.04 | 0.01 | 0.02 | 0.06 | 1 |
| ran_pars | cond | species | sd__(Intercept) | 0.04 | 0.03 | 0.00 | 0.10 | 1 |

**Posterior predictive check TUH; 100 simulated datasets**

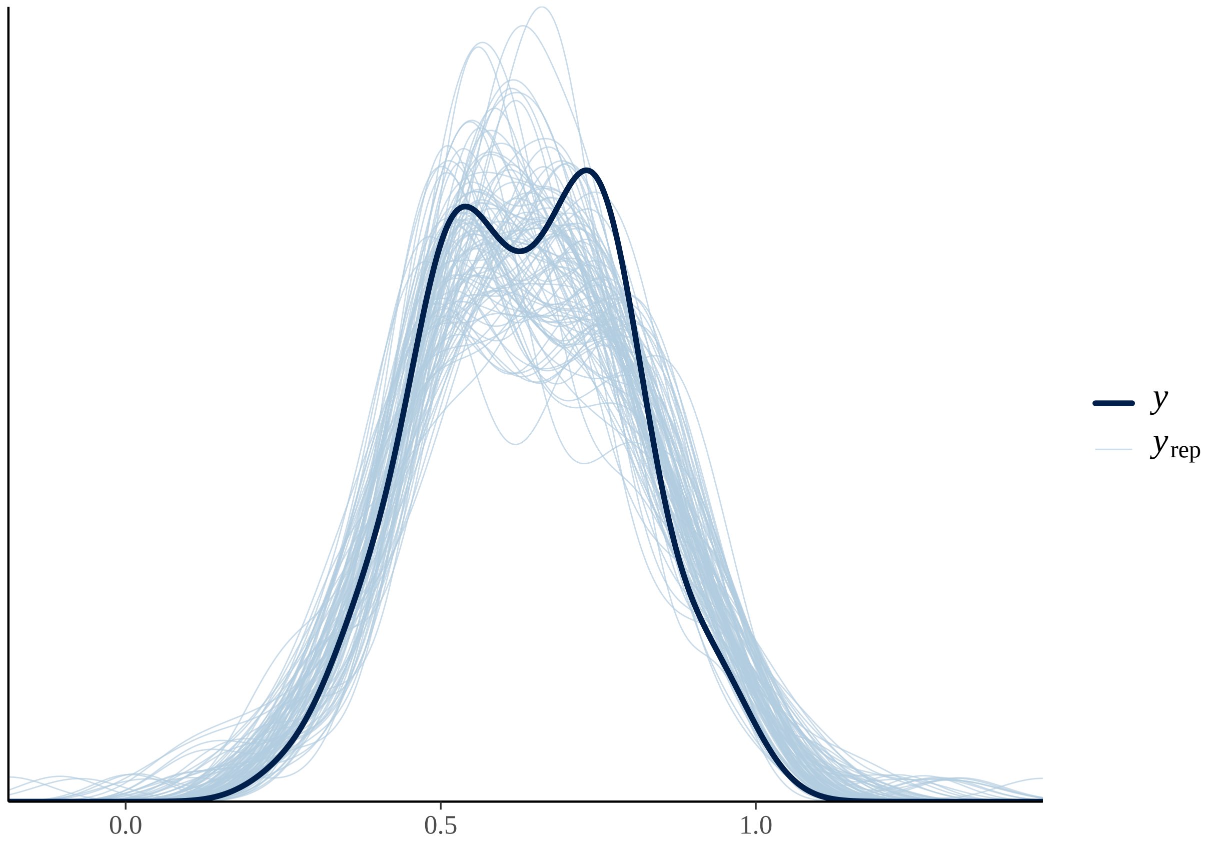

#### Fixed effects results TUH

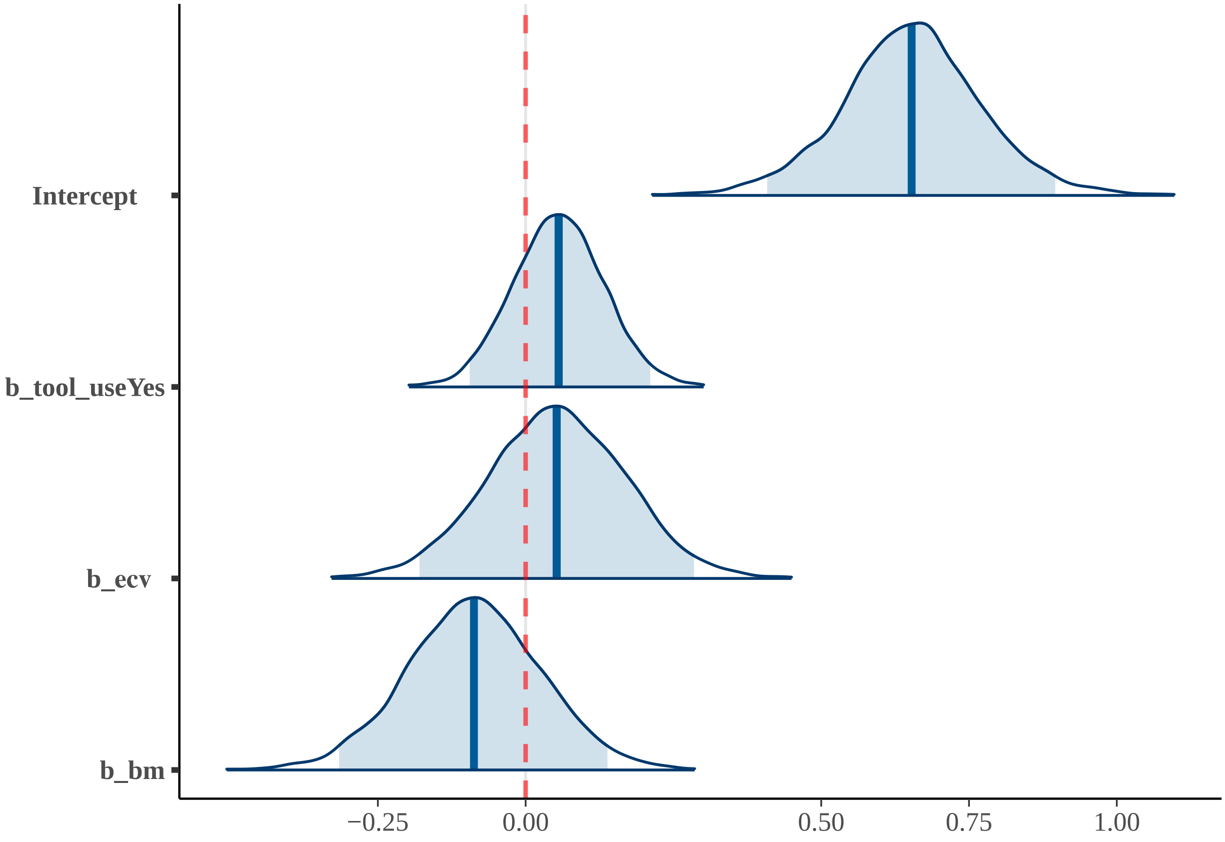


### 5) TU-SH

#### Summary TU-SH

| effect | component | term | estimate | std.error | conf.low | conf.high | rhat |
|---|---|---|---|---|---|---|---|
| fixed | cond | (Intercept) | 0.68 | 0.11 | 0.46 | 0.90 | 1 |
| fixed | cond | tool_useYes | 0.04 | 0.07 | -0.10 | 0.19 | 1 |
| fixed | cond | ecv | 0.04 | 0.11 | -0.17 | 0.25 | 1 |
| fixed | cond | bm | -0.05 | 0.11 | -0.26 | 0.16 | 1 |
| fixed | cond | substrateBOTH | -0.02 | 0.07 | -0.16 | 0.12 | 1 |
| fixed | cond | substrateT | -0.24 | 0.10 | -0.45 | -0.04 | 1 |

| effect | component | group | term | estimate | std.error | conf.low | conf.high | rhat |
|---|---|---|---|---|---|---|---|---|
| ran_pars | cond | obs | sd__(Intercept) | 0.09 | 0.02 | 0.06 | 0.13 | 1 |
| ran_pars | cond | phylo | sd__(Intercept) | 0.03 | 0.01 | 0.02 | 0.05 | 1 |
| ran_pars | cond | species | sd__(Intercept) | 0.03 | 0.02 | 0.00 | 0.09 | 1 |

### Posterior predictive check TU-SH; 100 simulated datasets

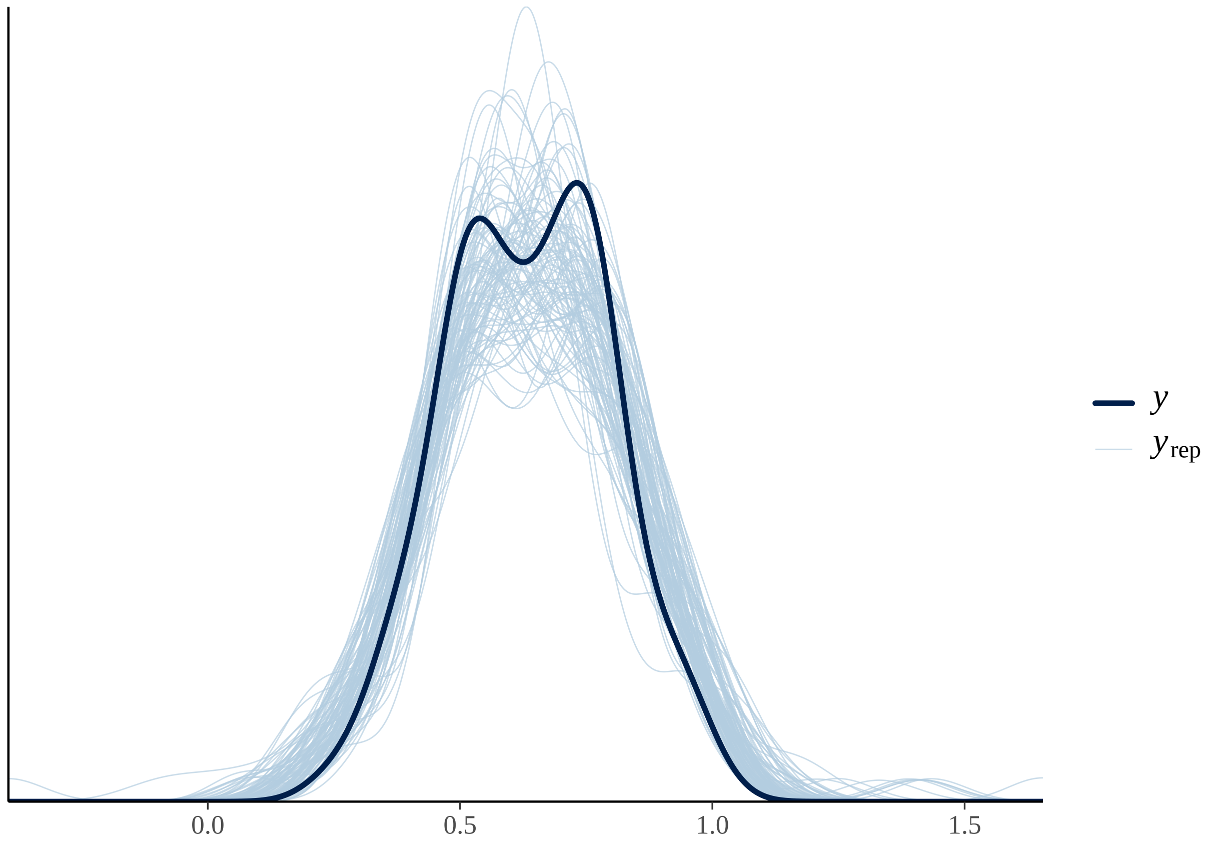

#### Fixed effects results TU-SH

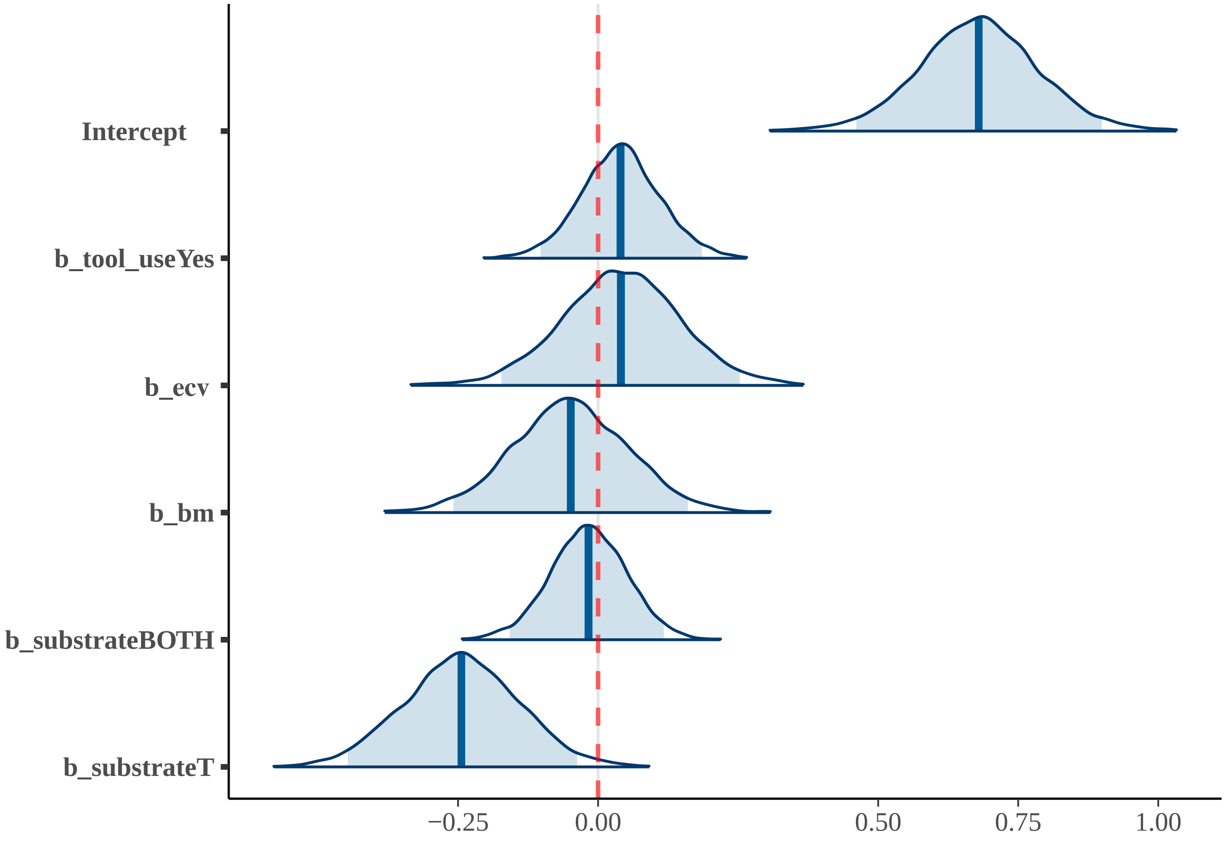


### 6) TU-BH

#### Summary TU-BH

| effect | component | term | estimate | std.error | conf.low | conf.high | rhat |
|---|---|---|---|---|---|---|---|
| fixed | cond | (Intercept) | 0.70 | 0.10 | 0.48 | 0.90 | 1 |
| fixed | cond | tool_useYes | 0.01 | 0.07 | -0.13 | 0.16 | 1 |
| fixed | cond | ecv | 0.08 | 0.10 | -0.13 | 0.28 | 1 |
| fixed | cond | bm | -0.05 | 0.10 | -0.25 | 0.15 | 1 |
| fixed | cond | substrateBOTH | -0.07 | 0.08 | -0.23 | 0.09 | 1 |
| fixed | cond | substrateT | -0.30 | 0.11 | -0.51 | -0.08 | 1 |
| fixed | cond | imi | -0.08 | 0.05 | -0.17 | 0.03 | 1 |

| effect | component | group | term | estimate | std.error | conf.low | conf.high | rhat |
|---|---|---|---|---|---|---|---|---|
| ran_pars | cond | obs | sd___(Intercept) | 0.10 | 0.02 | 0.06 | 0.13 | 1 |
| ran_pars | cond | phylo | sd___(Intercept) | 0.03 | 0.01 | 0.01 | 0.05 | 1 |
| ran_pars | cond | species | sd___(Intercept) | 0.03 | 0.03 | 0.00 | 0.10 | 1 |

#### Posterior predictive check TU-BH; 100 simulated datasets

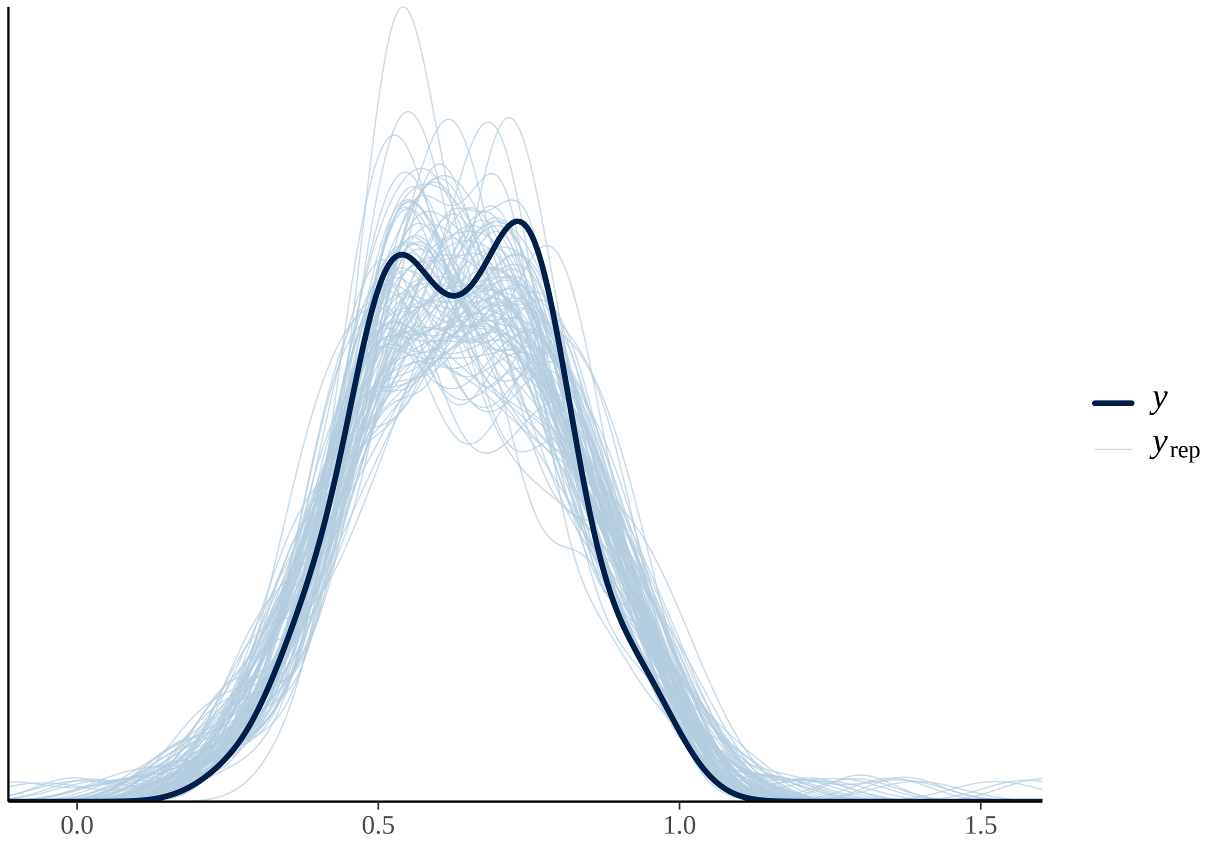

#### Fixed effects results TU-BH

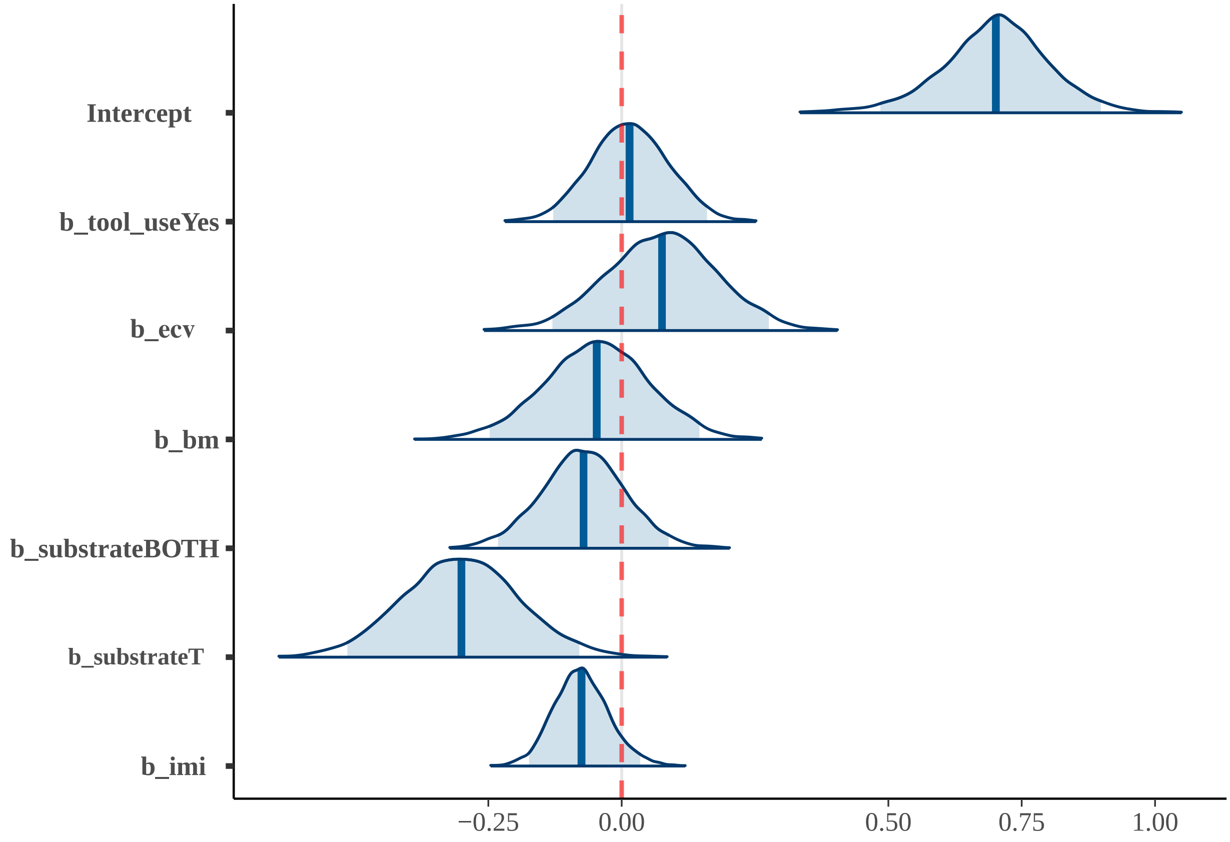


##### 7) TU-SH-SSH

#### Summary TU-SH-SSH

| effect | component | term | estimate | std.error | conf.low | conf.high | rhat |
|---|---|---|---|---|---|---|---|
| fixed | cond | (Intercept) | 0.75 | 0.09 | 0.55 | 0.92 | 1 |
| fixed | cond | tool_useYes | 0.03 | 0.07 | -0.10 | 0.17 | 1 |
| fixed | cond | substrateBOTH | -0.09 | 0.07 | -0.23 | 0.05 | 1 |
| fixed | cond | substrateT | -0.32 | 0.10 | -0.51 | -0.13 | 1 |
| fixed | cond | social_systemPair | -0.19 | 0.10 | -0.38 | 0.02 | 1 |
| fixed | cond | social_systemSolitary | -0.21 | 0.16 | -0.53 | 0.10 | 1 |

| effect | component | group | term | estimate | std.error | conf.low | conf.high | rhat |
|---|---|---|---|---|---|---|---|---|
| ran_pars | cond | obs | sd___(Intercept) | 0.10 | 0.02 | 0.07 | 0.14 | 1 |
| ran_pars | cond | phylo | sd___(Intercept) | 0.02 | 0.01 | 0.00 | 0.05 | 1 |
| ran_pars | cond | species | sd___(Intercept) | 0.03 | 0.03 | 0.00 | 0.09 | 1 |

**Posterior predictive check TU-SH-SSH; 100 simulated datasets**

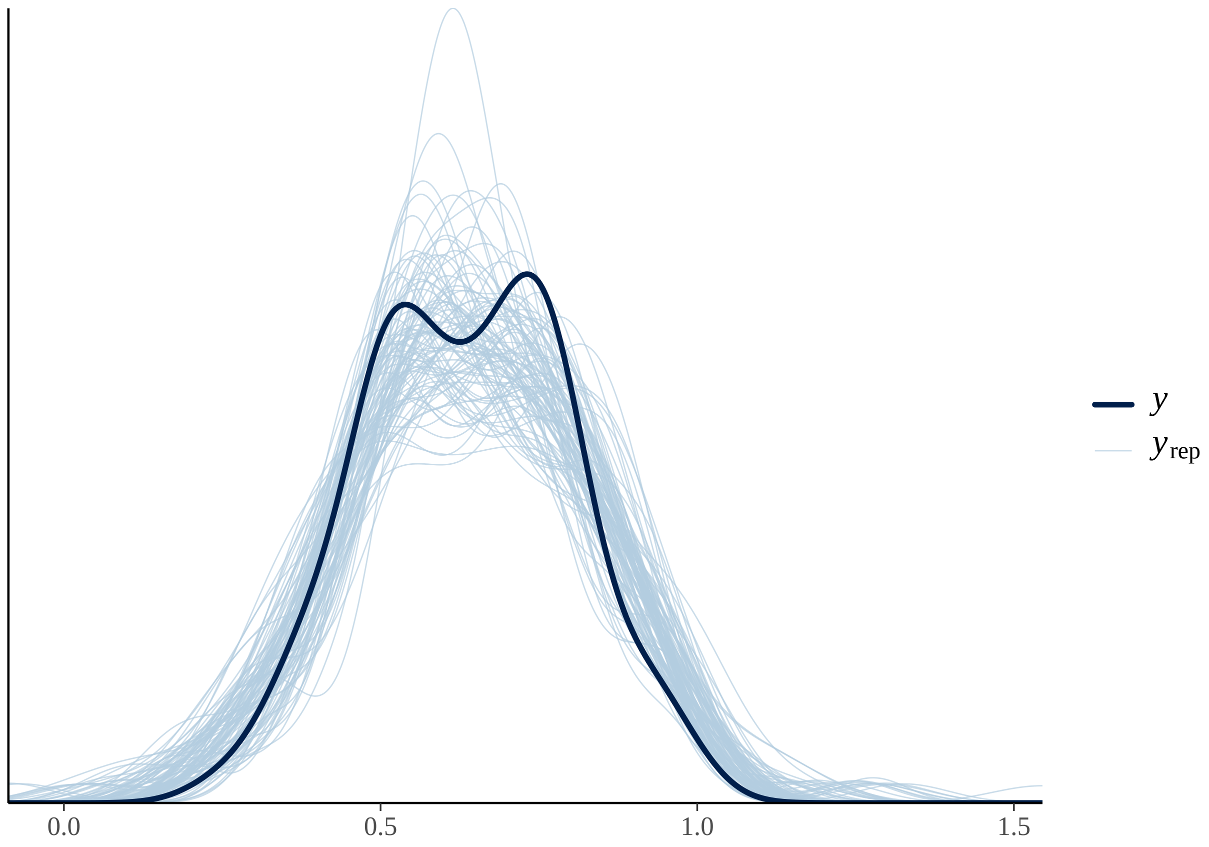

#### Fixed effects results TU-SH-SSH

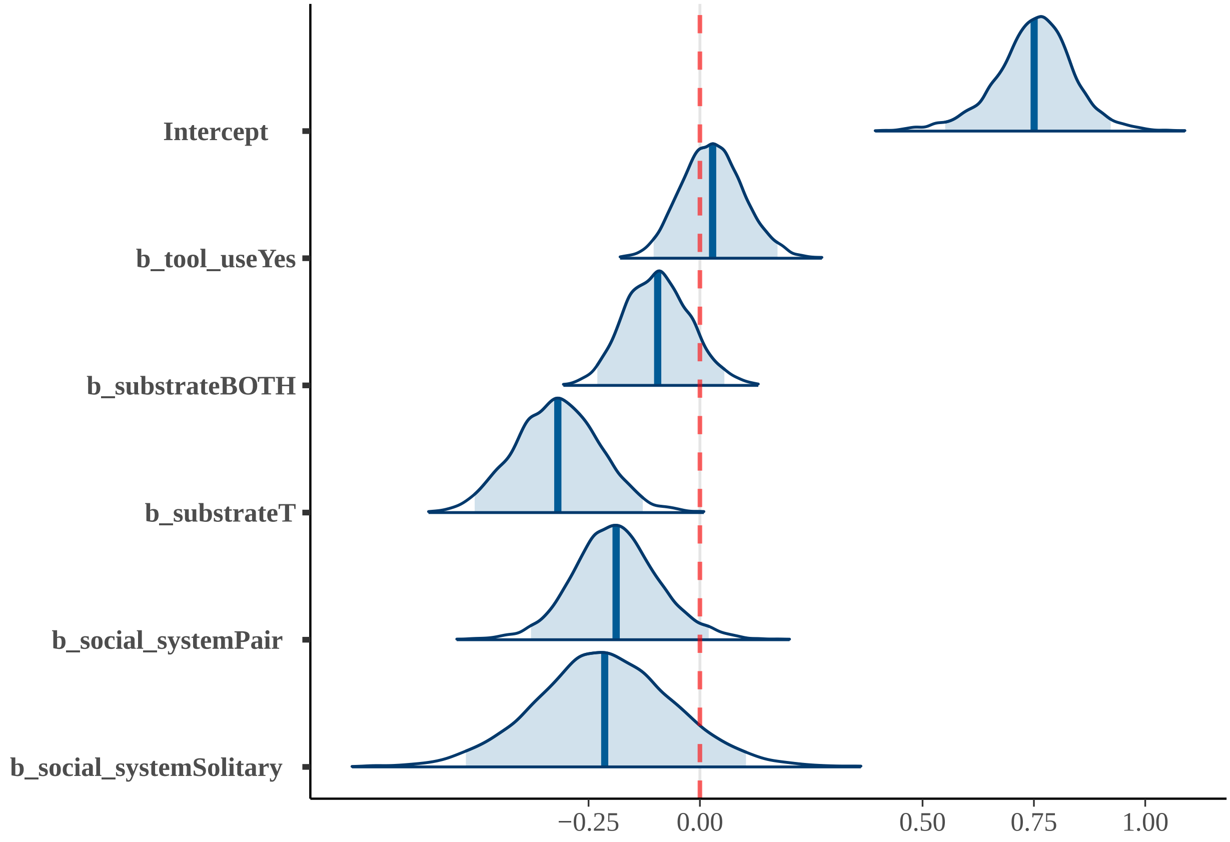


##### 8) SP-SS-B-TUH

#### Summary SP-SS-B-TUH

| effect | component | term | estimate | std.error | conf.low | conf.high | rhat |
|---|---|---|---|---|---|---|---|
| fixed | cond | (Intercept) | 0.75 | 0.10 | 0.54 | 0.95 | 1 |
| fixed | cond | tool_useYes | 0.04 | 0.07 | -0.10 | 0.18 | 1 |
| fixed | cond | substrateBOTH | -0.06 | 0.08 | -0.21 | 0.10 | 1 |
| fixed | cond | substrateT | -0.26 | 0.11 | -0.47 | -0.05 | 1 |
| fixed | cond | social_systemPair | -0.29 | 0.17 | -0.63 | 0.05 | 1 |
| fixed | cond | social_systemSolitary | -0.24 | 0.21 | -0.66 | 0.17 | 1 |
| fixed | cond | bm | -0.06 | 0.07 | -0.20 | 0.07 | 1 |
| fixed | cond | imi | 0.04 | 0.08 | -0.11 | 0.20 | 1 |

| effect | component | group | term | estimate | std.error | conf.low | conf.high | rhat |
|---|---|---|---|---|---|---|---|---|
| ran_pars | cond | obs | sd__(Intercept) | 0.09 | 0.02 | 0.06 | 0.13 | 1 |
| ran_pars | cond | phylo | sd__(Intercept) | 0.03 | 0.01 | 0.01 | 0.05 | 1 |
| ran_pars | cond | species | sd__(Intercept) | 0.03 | 0.02 | 0.00 | 0.09 | 1 |

**Posterior predictive check SP-SS-B-TUH; 100 simulated datasets**

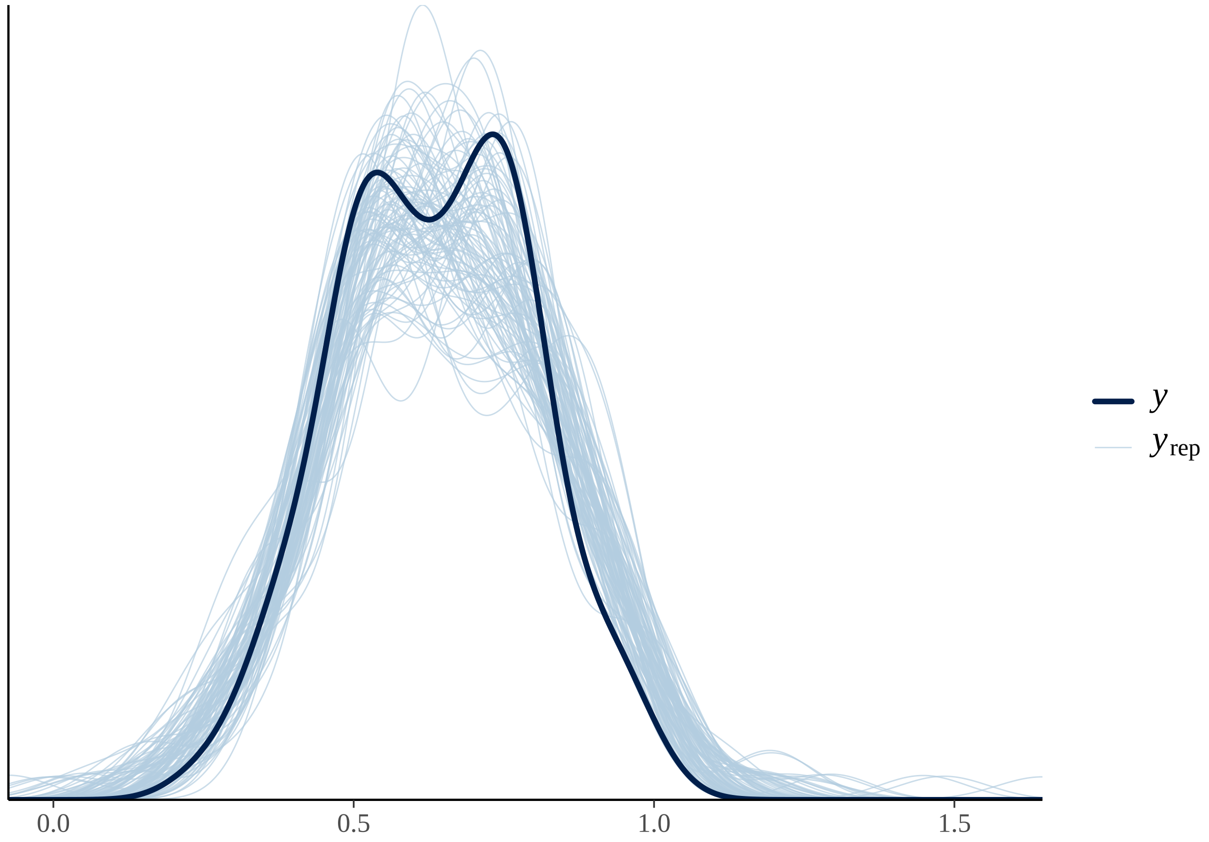

#### Fixed effects results SP-SS-B-TUH

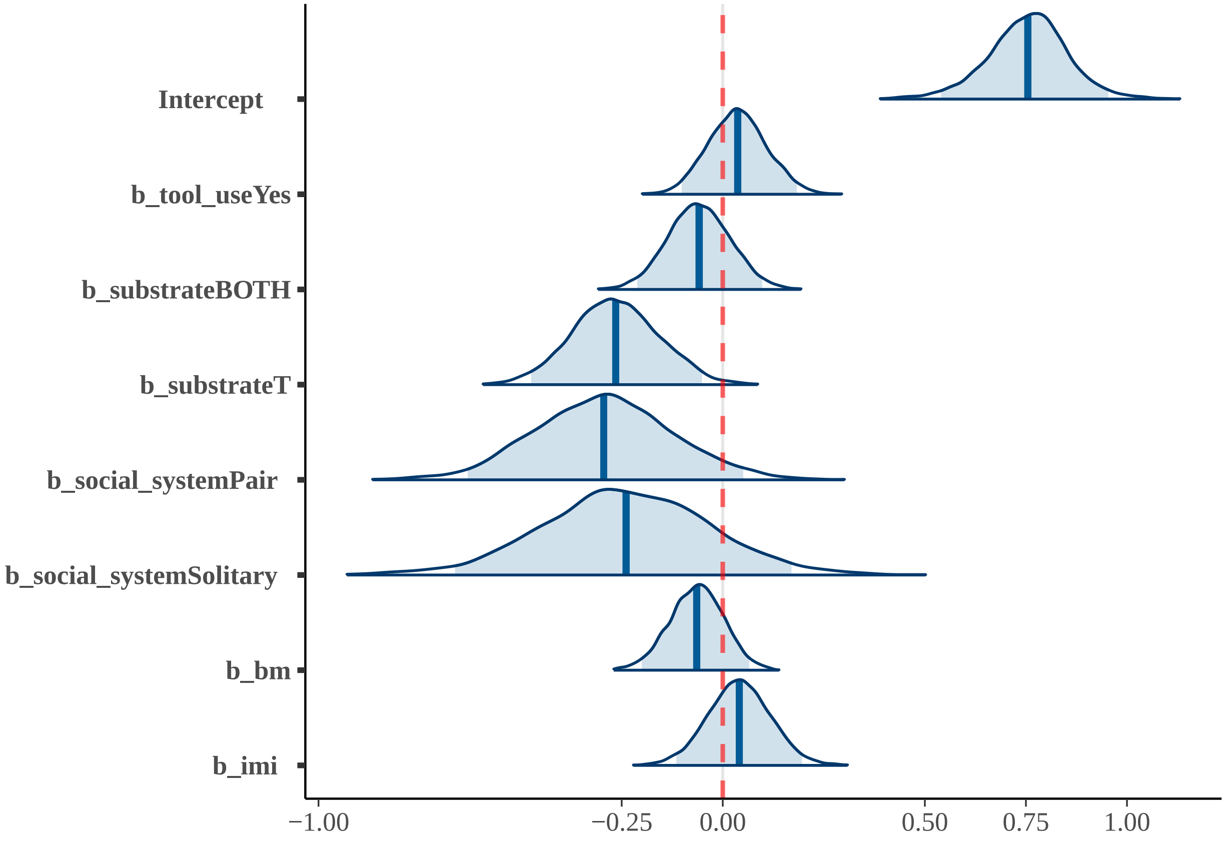


### 9) FH

#### Summary FH

| effect | component | term | estimate | std.error | conf.low | conf.high | rhat |
|---|---|---|---|---|---|---|---|
| fixed | cond | (Intercept) | 0.67 | 0.14 | 0.40 | 0.96 | 1 |
| fixed | cond | dimo | -0.07 | 0.04 | -0.14 | 0.00 | 1 |
| fixed | cond | cl2 | -0.09 | 0.13 | -0.35 | 0.15 | 1 |
| fixed | cond | cl3 | 0.08 | 0.14 | -0.20 | 0.35 | 1 |
| fixed | cond | cl4 | -0.04 | 0.15 | -0.35 | 0.24 | 1 |

| effect | component | group | term | estimate | std.error | conf.low | conf.high | rhat |
|---|---|---|---|---|---|---|---|---|
| ran_pars | cond | obs | sd__(Intercept) | 0.10 | 0.02 | 0.06 | 0.14 | 1 |
| ran_pars | cond | phylo | sd__(Intercept) | 0.03 | 0.01 | 0.01 | 0.05 | 1 |
| ran_pars | cond | species | sd__(Intercept) | 0.04 | 0.03 | 0.00 | 0.10 | 1 |

**Posterior predictive check FH; 100 simulated datasets**

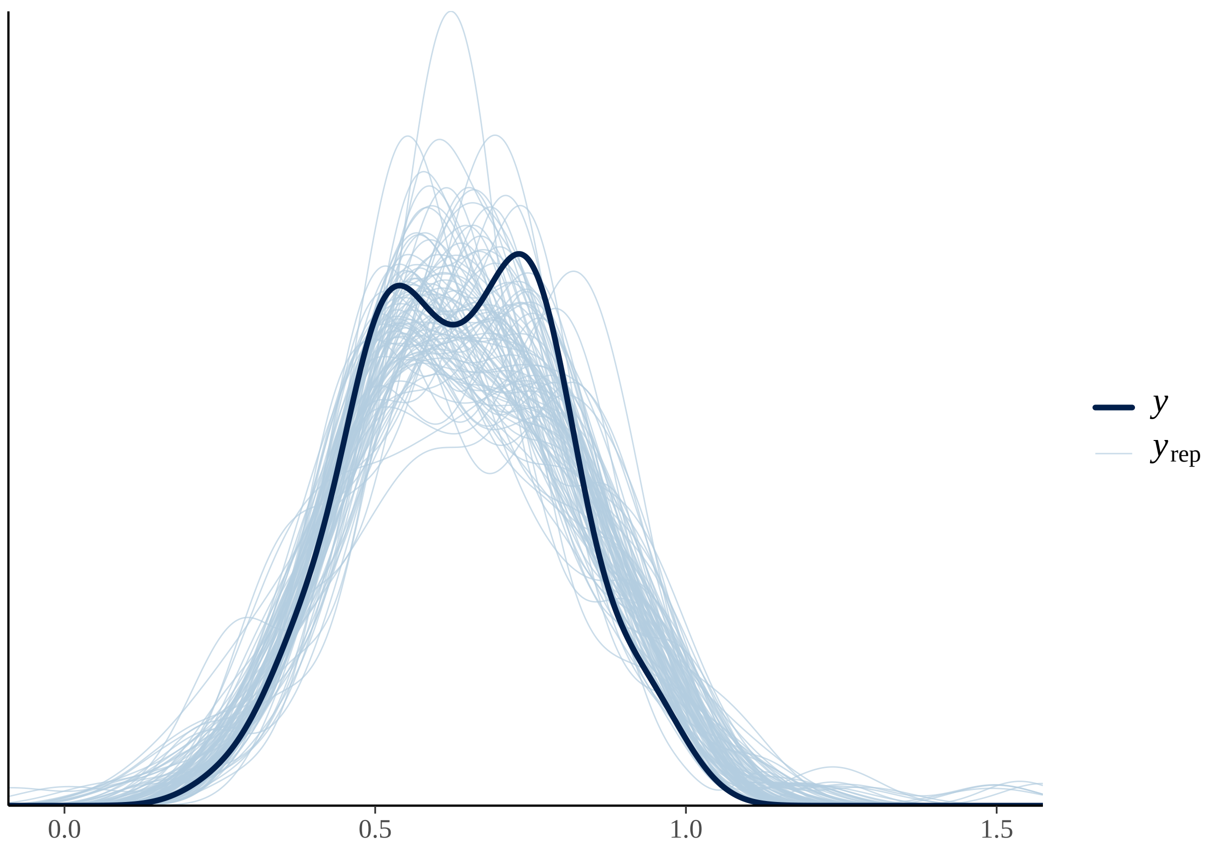

#### Fixed effects results FH

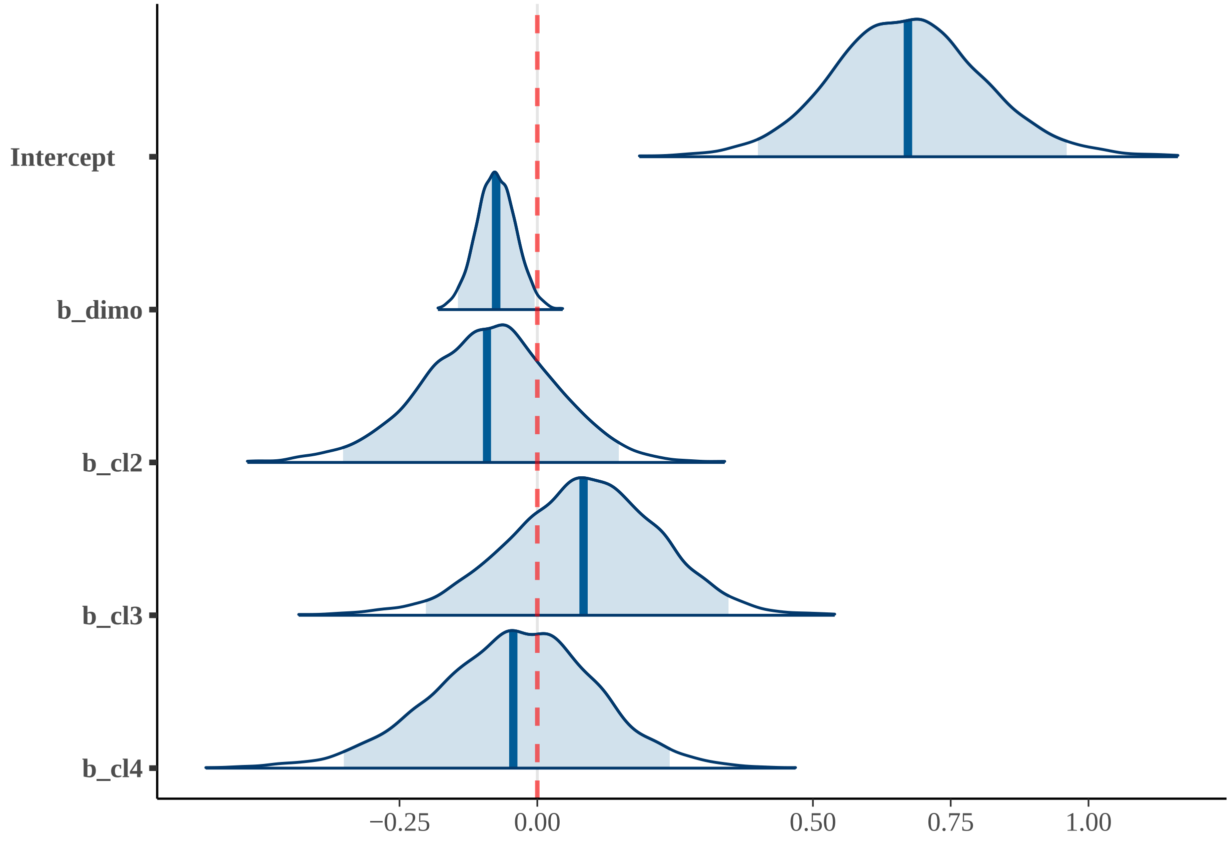


##### 10) EFH

#### Summary EFH

| effect | component | term | estimate | std.error | conf.low | conf.high | rhat |
|---|---|---|---|---|---|---|---|
| fixed | cond | (Intercept) | 0.68 | 0.11 | 0.45 | 0.90 | 1 |
| fixed | cond | extractive | -0.05 | 0.08 | -0.21 | 0.12 | 1 |
| fixed | cond | social_learning | -0.02 | 0.07 | -0.16 | 0.12 | 1 |
| fixed | cond | tool_useYes | 0.08 | 0.08 | -0.07 | 0.23 | 1 |

| effect | component | group | term | estimate | std.error | conf.low | conf.high | rhat |
|---|---|---|---|---|---|---|---|---|
| ran_pars | cond | obs | sd__(Intercept) | 0.10 | 0.02 | 0.07 | 0.14 | 1 |
| ran_pars | cond | phylo | sd__(Intercept) | 0.03 | 0.01 | 0.01 | 0.06 | 1 |
| ran_pars | cond | species | sd__(Intercept) | 0.04 | 0.03 | 0.00 | 0.11 | 1 |

**Posterior predictive check EFH; 100 simulated datasets**

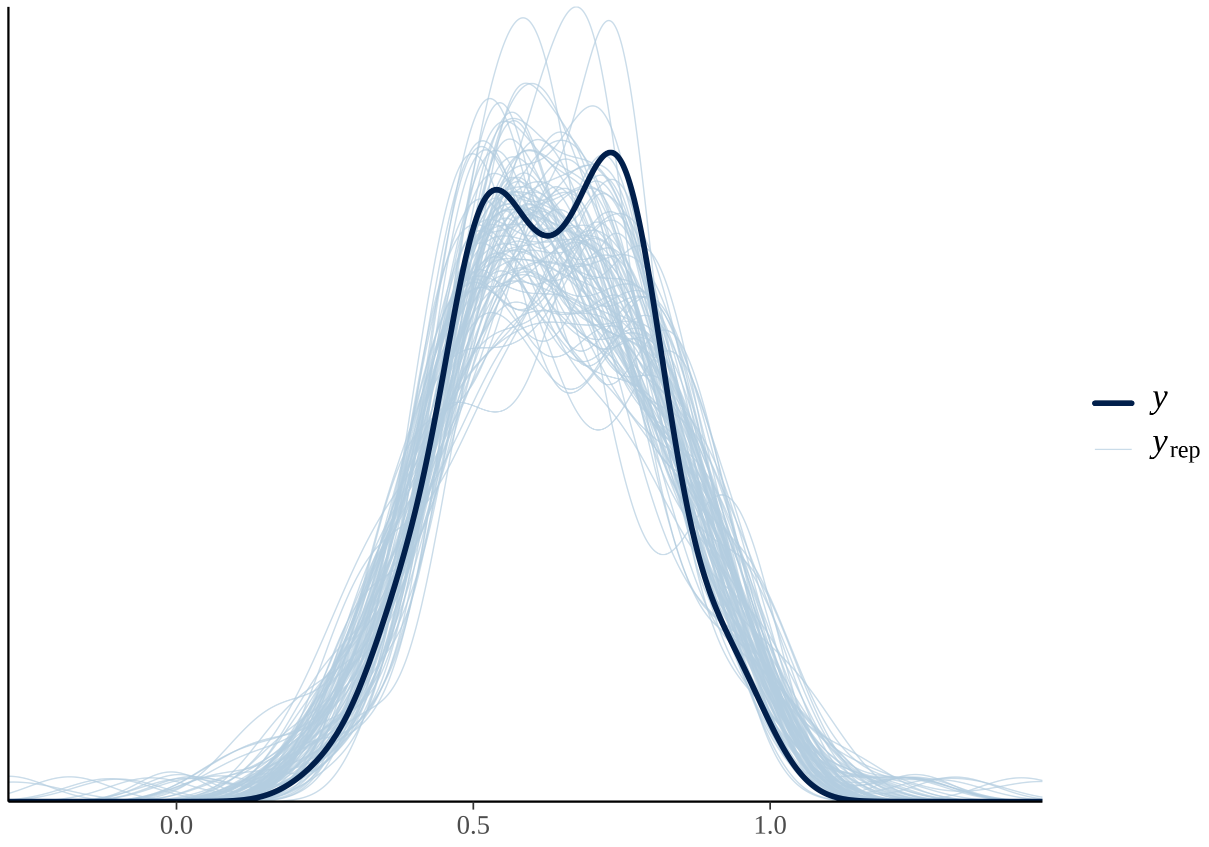

#### Fixed effects results EFH

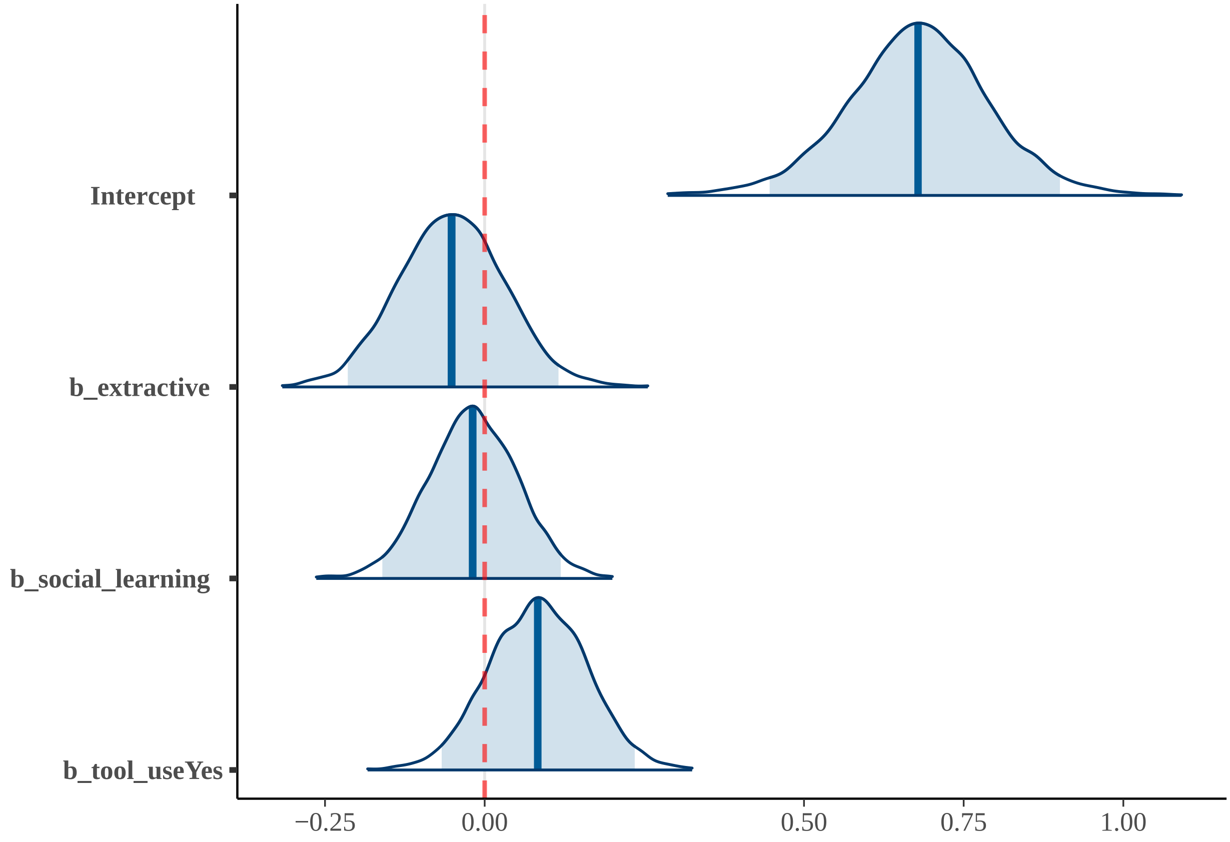


##### 11) Reduced model

#### Summary Reduced model

| effect | component | term | estimate | std.error | conf.low | conf.high | rhat |
|---|---|---|---|---|---|---|---|
| fixed | cond | (Intercept) | 0.69 | 0.10 | 0.50 | 0.88 | 1 |
| fixed | cond | substrateBOTH | -0.02 | 0.06 | -0.15 | 0.10 | 1 |
| fixed | cond | substrateT | -0.26 | 0.10 | -0.44 | -0.07 | 1 |

| effect | component | group | term | estimate | std.error | conf.low | conf.high | rhat |
|---|---|---|---|---|---|---|---|---|
| ran_pars | cond | obs | sd__(Intercept) | 0.10 | 0.02 | 0.06 | 0.13 | 1 |
| ran_pars | cond | phylo | sd__(Intercept) | 0.03 | 0.01 | 0.01 | 0.05 | 1 |
| ran_pars | cond | species | sd__(Intercept) | 0.03 | 0.02 | 0.00 | 0.09 | 1 |

**Posterior predictive check Reduced model; 100 simulated datasets**

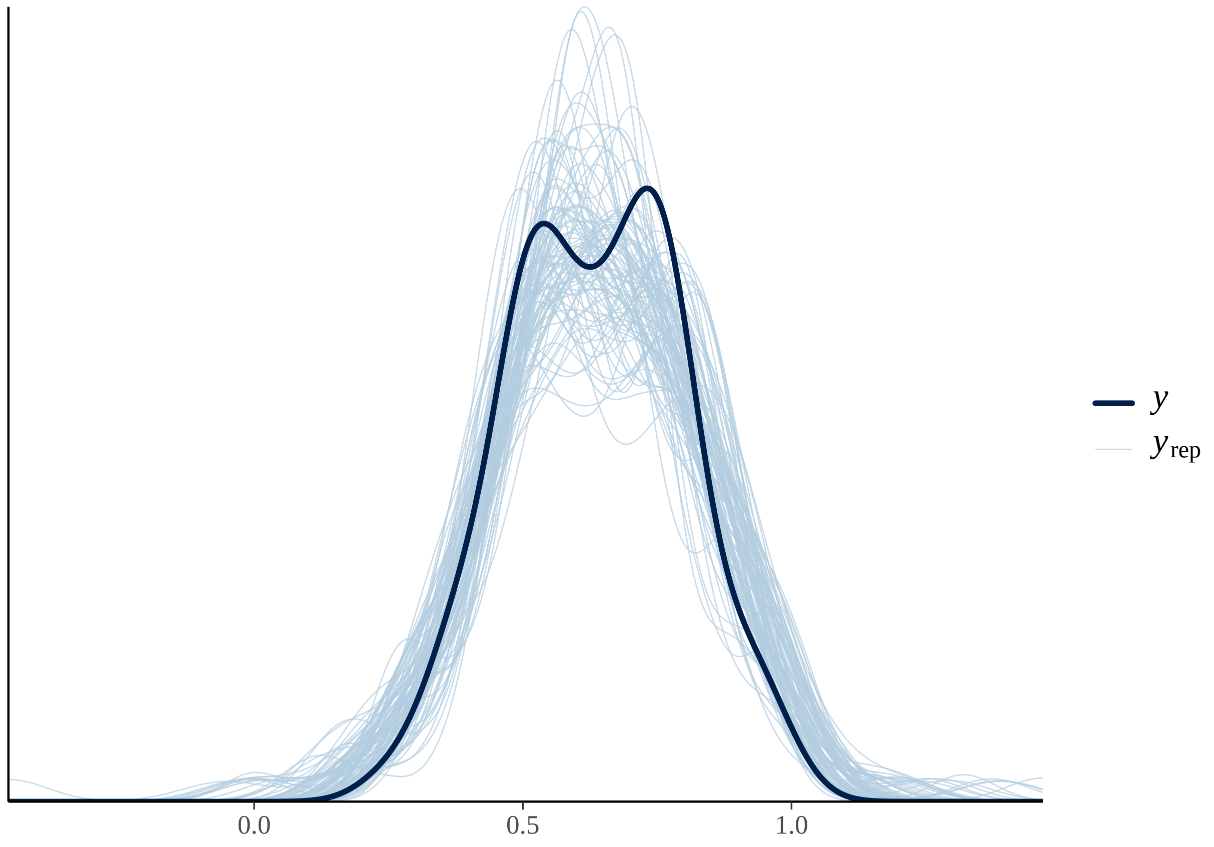

#### Fixed effects results Reduced model

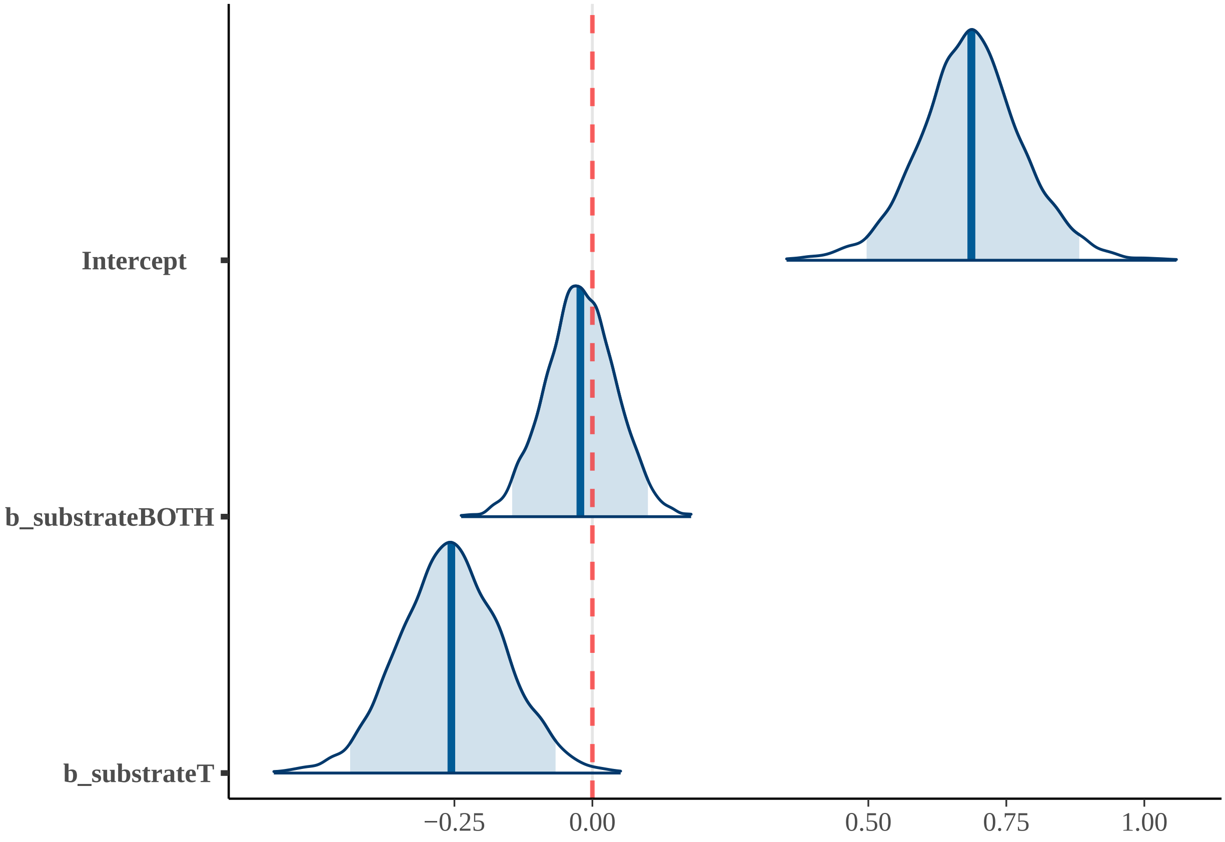

**MABSHI results including *Homo sapiens***

Numerical results for all the hypotheses tested, using MABSHI as the dependent variable, are presented here (see Supplementary Table 1 for further details). *Homo sapiens* was included in these analyses. In addition, the final reduced model is also presented after all the hypotheses tested. Posterior predictive checks for each of the tested hypotheses are shown, along with plots illustrating the estimates for the fixed effects included in each of the tested models.

#### 0) Intercepts-only model

### Summary

| effect | component | term | estimate | std.error | conf.low | conf.high | rhat |
|---|---|---|---|---|---|---|---|
| fixed | cond | (Intercept) | 0.67 | 0.11 | 0.44 | 0.91 | 1 |

| effect | component | group | term | estimate | std.error | conf.low | conf.high | rhat |
|---|---|---|---|---|---|---|---|---|
| ran_pars | cond | obs | sd__(Intercept) | 0.11 | 0.02 | 0.07 | 0.15 | 1 |
| ran_pars | cond | phylo | sd__(Intercept) | 0.03 | 0.01 | 0.01 | 0.06 | 1 |
| ran_pars | cond | species | sd__(Intercept) | 0.04 | 0.03 | 0.00 | 0.12 | 1 |

#### Posterior predictive check; 100 simulated datasets

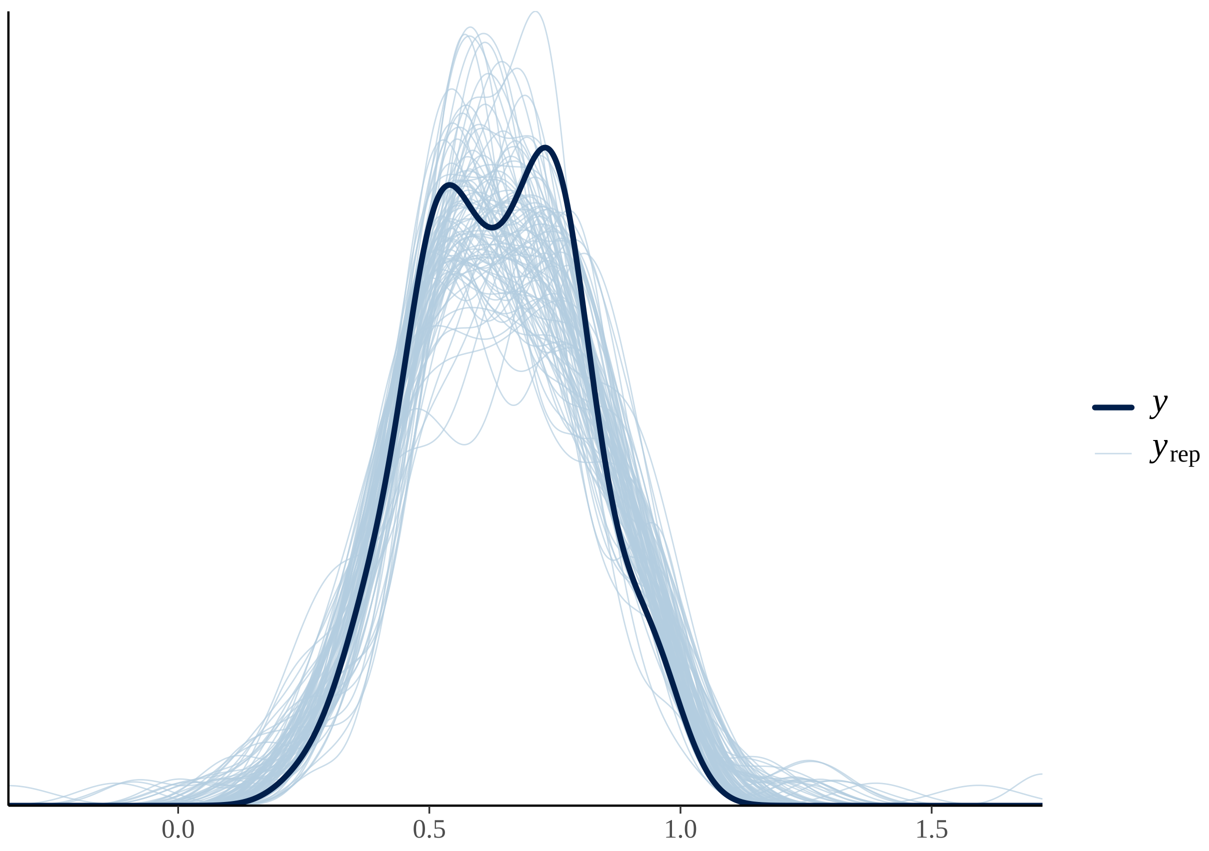

#### Fixed effects results

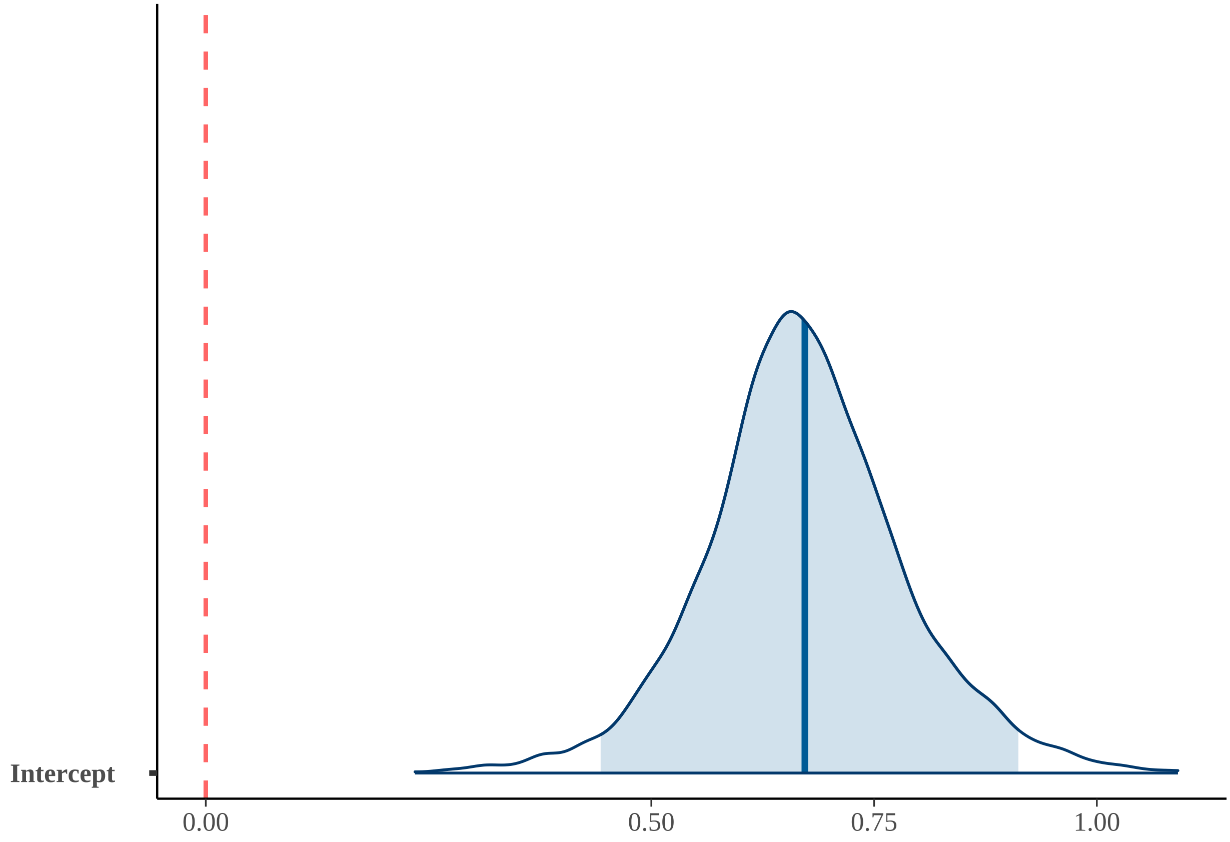


##### 1) POH1

#### Summary POH1

| effect | component | term | estimate | std.error | conf.low | conf.high | rhat |
|---|---|---|---|---|---|---|---|
| fixed | cond | (Intercept) | 0.67 | 0.12 | 0.44 | 0.92 | 1 |
| fixed | cond | fru | 0.02 | 0.02 | -0.02 | 0.07 | 1 |
| fixed | cond | substrateBOTH | -0.02 | 0.07 | -0.16 | 0.13 | 1 |
| fixed | cond | substrateT | -0.08 | 0.10 | -0.28 | 0.12 | 1 |

| effect | component | group | term | estimate | std.error | conf.low | conf.high | rhat |
|---|---|---|---|---|---|---|---|---|
| ran_pars | cond | obs | sd__(Intercept) | 0.11 | 0.02 | 0.07 | 0.15 | 1 |
| ran_pars | cond | phylo | sd__(Intercept) | 0.03 | 0.01 | 0.01 | 0.06 | 1 |
| ran_pars | cond | species | sd__(Intercept) | 0.05 | 0.04 | 0.00 | 0.13 | 1 |

#### Posterior predictive check POH1; 100 simulated datasets

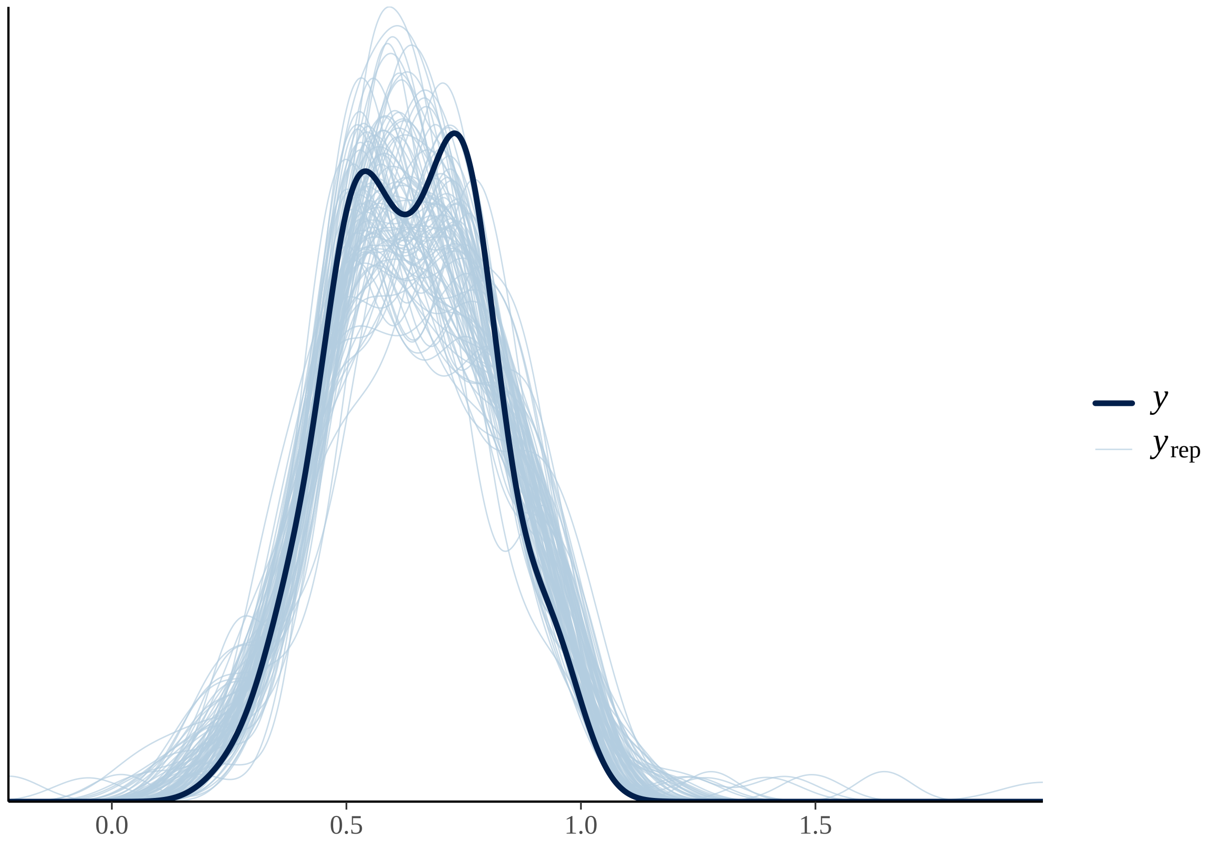

#### Fixed effects results POH1

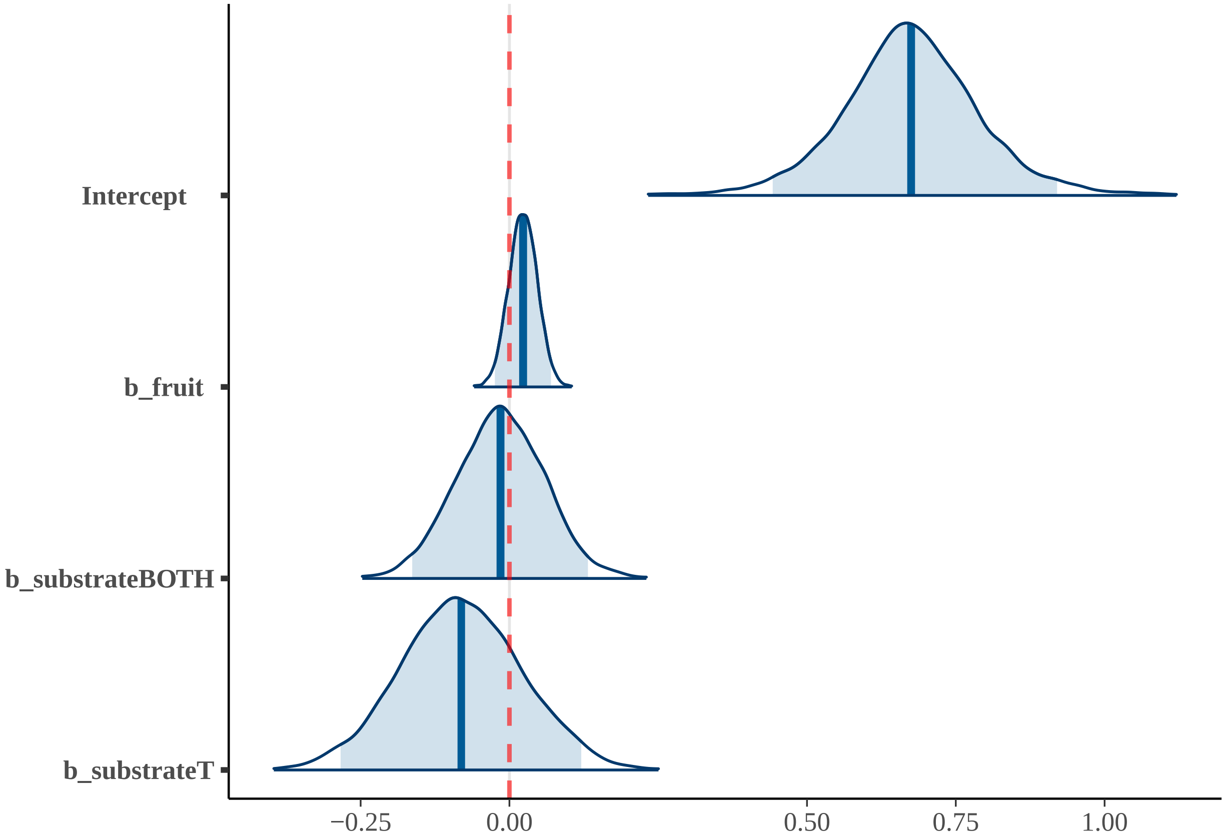


##### 2) POH2

#### Summary POH2

| effect | component | term | estimate | std.error | conf.low | conf.high | rhat |
|---|---|---|---|---|---|---|---|
| fixed | cond | (Intercept) | 0.63 | 0.13 | 0.38 | 0.87 | 1 |
| fixed | cond | diet | 0.04 | 0.02 | 0.00 | 0.08 | 1 |
| fixed | cond | substrateBOTH | -0.01 | 0.07 | -0.15 | 0.14 | 1 |
| fixed | cond | substrateT | -0.10 | 0.10 | -0.29 | 0.09 | 1 |

| effect | component | group | term | estimate | std.error | conf.low | conf.high | rhat |
|---|---|---|---|---|---|---|---|---|
| ran_pars | cond | obs | sd__(Intercept) | 0.10 | 0.02 | 0.07 | 0.14 | 1 |
| ran_pars | cond | phylo | sd__(Intercept) | 0.04 | 0.01 | 0.02 | 0.06 | 1 |
| ran_pars | cond | species | sd__(Intercept) | 0.04 | 0.03 | 0.00 | 0.11 | 1 |

#### Posterior predictive check POH2; 100 simulated datasets

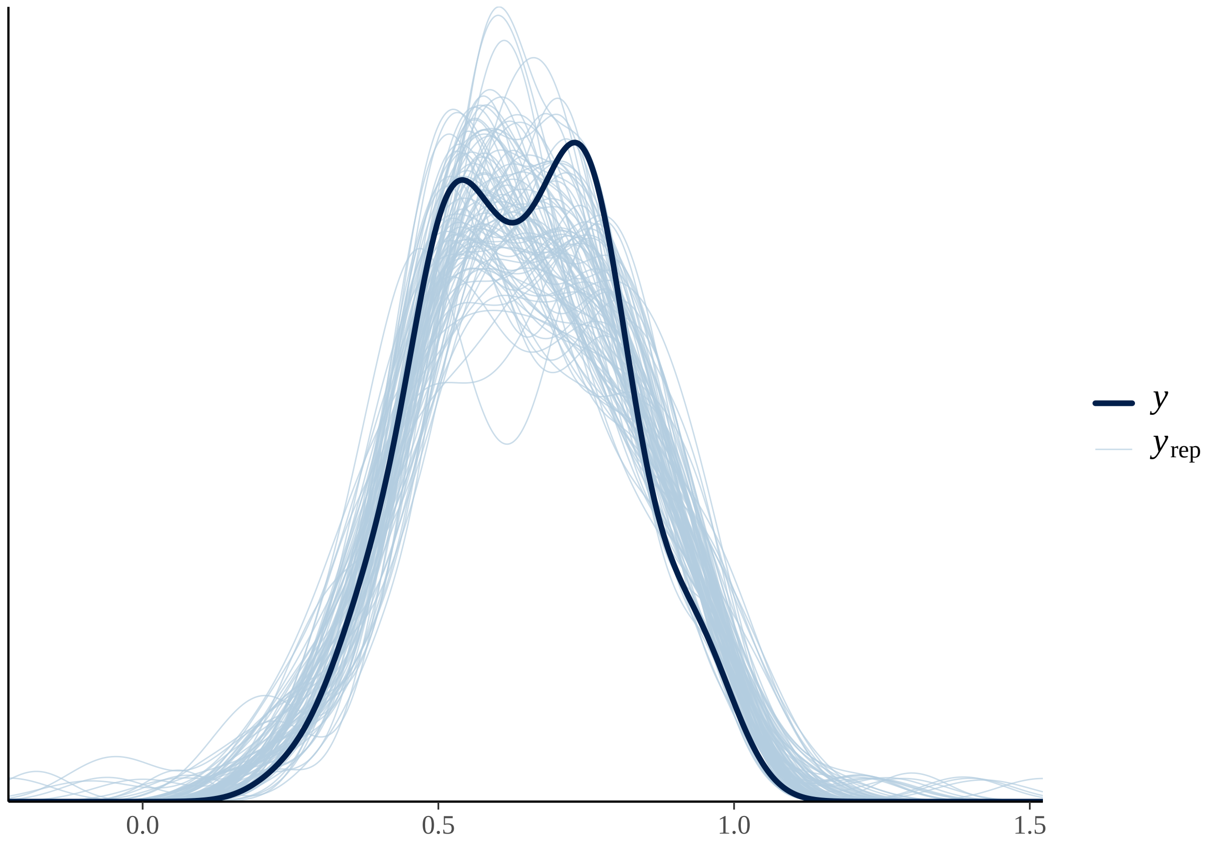

#### Fixed effects results POH2

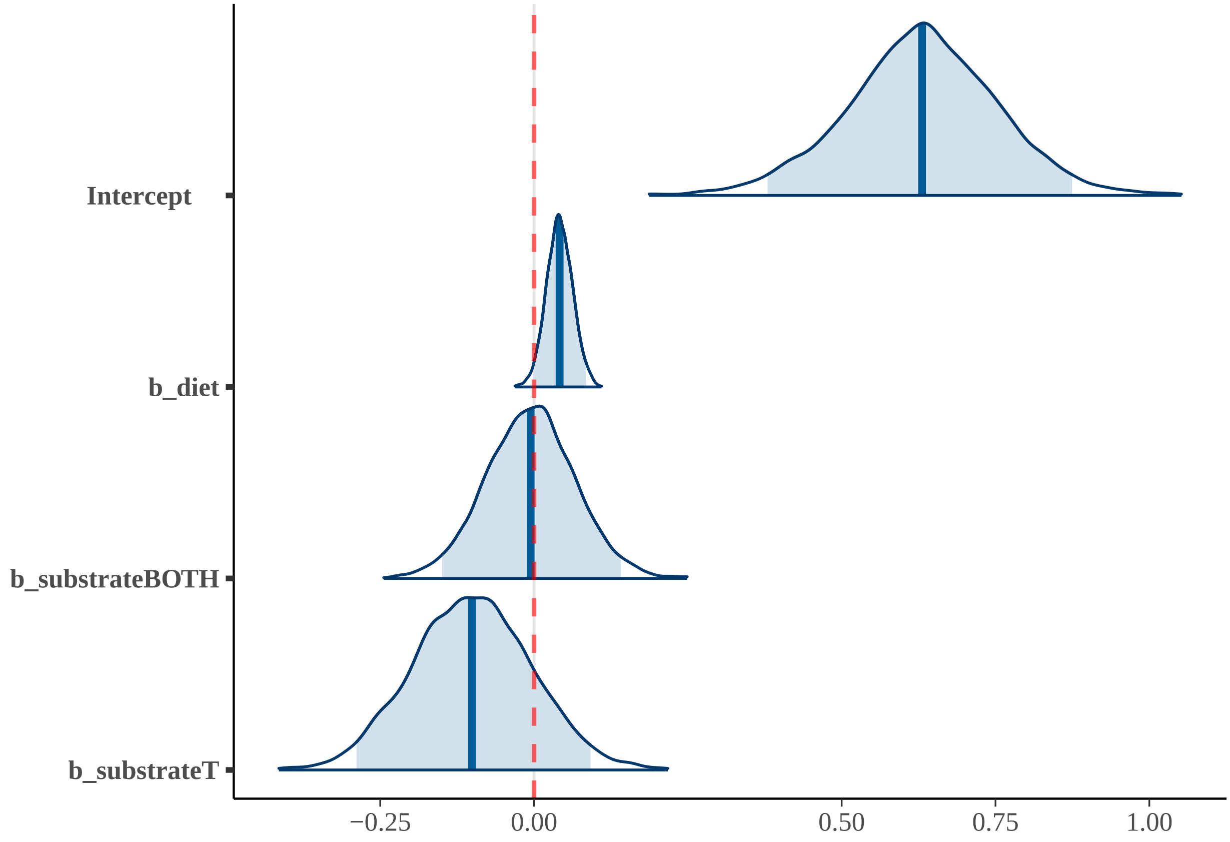


### 3) BH

#### Summary BH

| effect | component | term | estimate | std.error | conf.low | conf.high | rhat |
|---|---|---|---|---|---|---|---|
| fixed | cond | (Intercept) | 0.72 | 0.09 | 0.54 | 0.91 | 1 |
| fixed | cond | imi2 | -0.11 | 0.04 | -0.19 | -0.02 | 1 |
| fixed | cond | cc | 0.18 | 0.07 | 0.03 | 0.32 | 1 |
| fixed | cond | bm | -0.12 | 0.08 | -0.28 | 0.05 | 1 |
| fixed | cond | substrateBOTH | -0.10 | 0.08 | -0.25 | 0.05 | 1 |
| fixed | cond | substrateT | -0.29 | 0.11 | -0.50 | -0.08 | 1 |

| effect | component | group | term | estimate | std.error | conf.low | conf.high | rhat |
|---|---|---|---|---|---|---|---|---|
| ran_pars | cond | obs | sd__(Intercept) | 0.10 | 0.02 | 0.07 | 0.13 | 1 |
| ran_pars | cond | phylo | sd__(Intercept) | 0.03 | 0.01 | 0.01 | 0.05 | 1 |
| ran_pars | cond | species | sd__(Intercept) | 0.04 | 0.03 | 0.00 | 0.10 | 1 |

**Posterior predictive check BH; 100 simulated datasets**

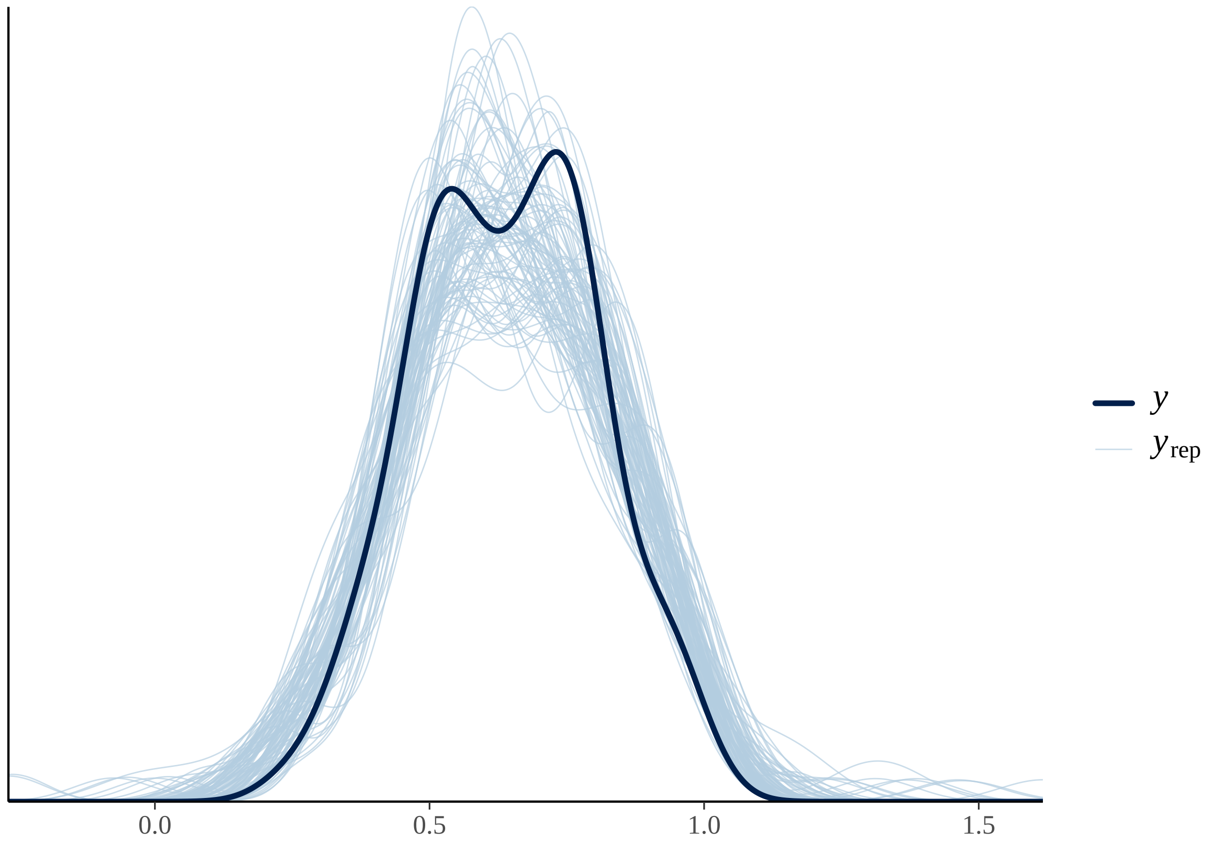

#### Fixed effects results BH

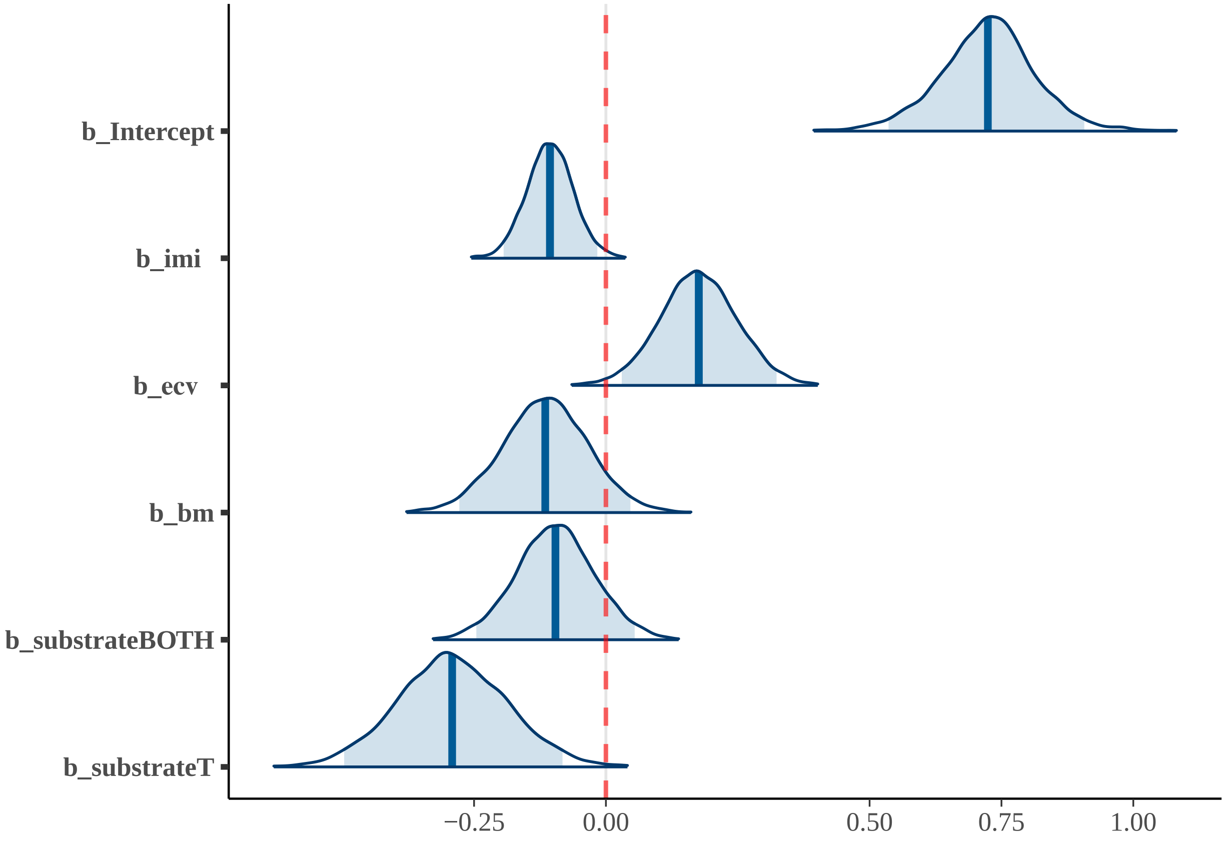


##### 4) TUH

#### Summary TUH

| effect | component | term | estimate | std.error | conf.low | conf.high | rhat |
|---|---|---|---|---|---|---|---|
| fixed | cond | (Intercept) | 0.67 | 0.12 | 0.43 | 0.91 | 1 |
| fixed | cond | tool_useYes | 0.06 | 0.08 | -0.09 | 0.21 | 1 |
| fixed | cond | cc | 0.15 | 0.08 | 0.00 | 0.31 | 1 |
| fixed | cond | bm | -0.17 | 0.08 | -0.34 | -0.01 | 1 |

| effect | component | group | term | estimate | std.error | conf.low | conf.high | rhat |
|---|---|---|---|---|---|---|---|---|
| ran_pars | cond | obs | sd__(Intercept) | 0.10 | 0.02 | 0.07 | 0.14 | 1 |
| ran_pars | cond | phylo | sd__(Intercept) | 0.04 | 0.01 | 0.02 | 0.06 | 1 |
| ran_pars | cond | species | sd__(Intercept) | 0.04 | 0.03 | 0.00 | 0.10 | 1 |

**Posterior predictive check TUH; 100 simulated datasets**

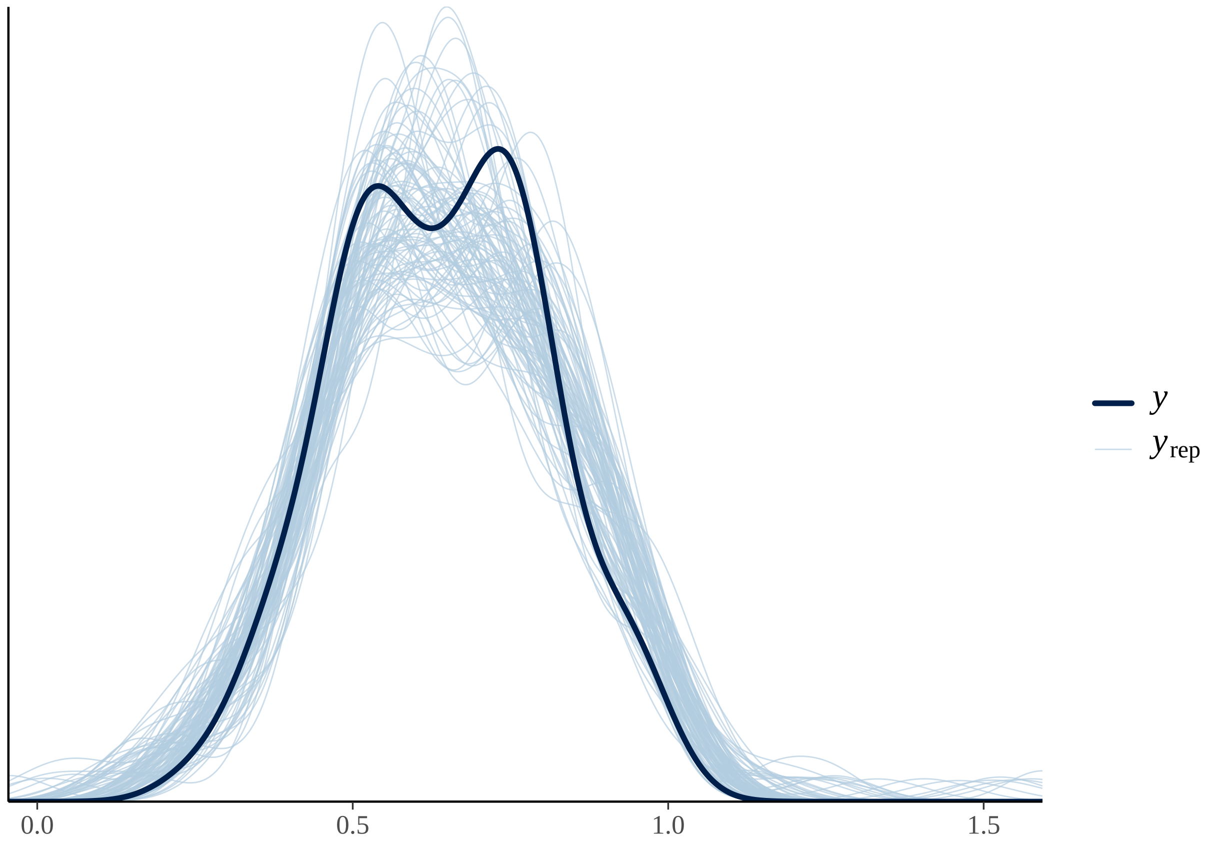

#### Fixed effects results TUH

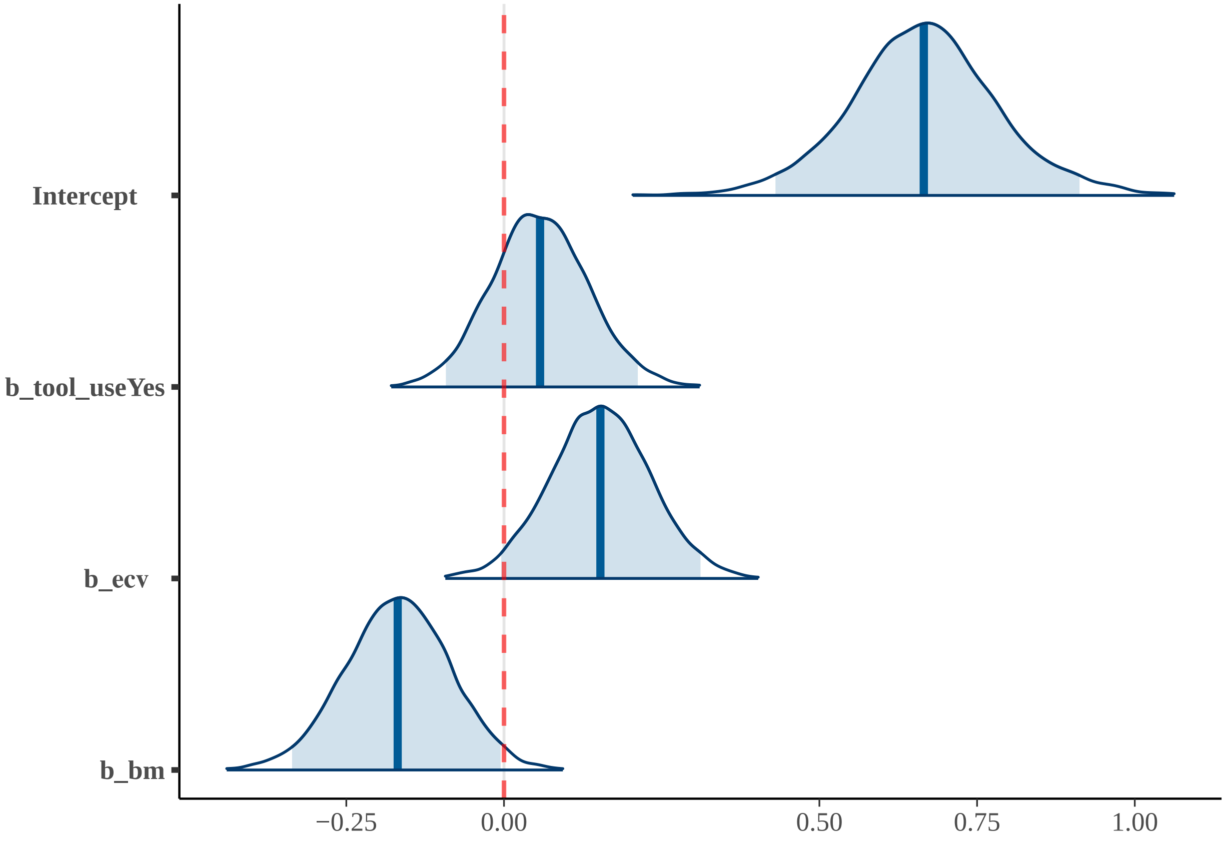


### 5) TU-SH

#### Summary TU-SH

| effect | component | term | estimate | std.error | conf.low | conf.high | rhat |
|---|---|---|---|---|---|---|---|
| fixed | cond | (Intercept) | 0.70 | 0.12 | 0.46 | 0.95 | 1 |
| fixed | cond | tool_useYes | 0.05 | 0.08 | -0.10 | 0.20 | 1 |
| fixed | cond | ecv | 0.21 | 0.08 | 0.05 | 0.38 | 1 |
| fixed | cond | bm | -0.20 | 0.08 | -0.36 | -0.03 | 1 |
| fixed | cond | substrateBOTH | -0.03 | 0.07 | -0.17 | 0.11 | 1 |
| fixed | cond | substrateT | -0.18 | 0.11 | -0.39 | 0.02 | 1 |

| effect | component | group | term | estimate | std.error | conf.low | conf.high | rhat |
|---|---|---|---|---|---|---|---|---|
| ran_pars | cond | obs | sd__(Intercept) | 0.09 | 0.02 | 0.06 | 0.13 | 1 |
| ran_pars | cond | phylo | sd__(Intercept) | 0.04 | 0.01 | 0.02 | 0.06 | 1 |
| ran_pars | cond | species | sd__(Intercept) | 0.04 | 0.03 | 0.00 | 0.10 | 1 |

#### Posterior predictive check TU-SH; 100 simulated datasets

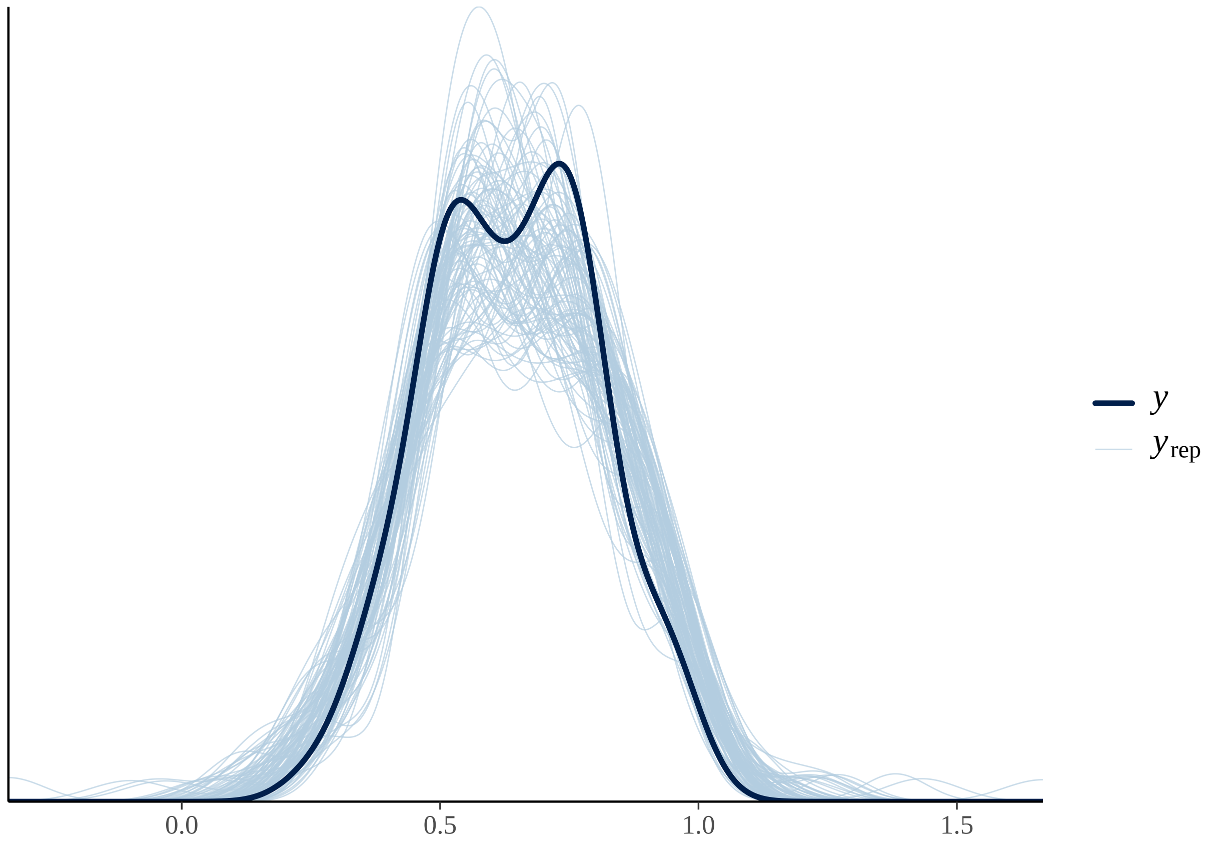

#### Fixed effects results TU-SH

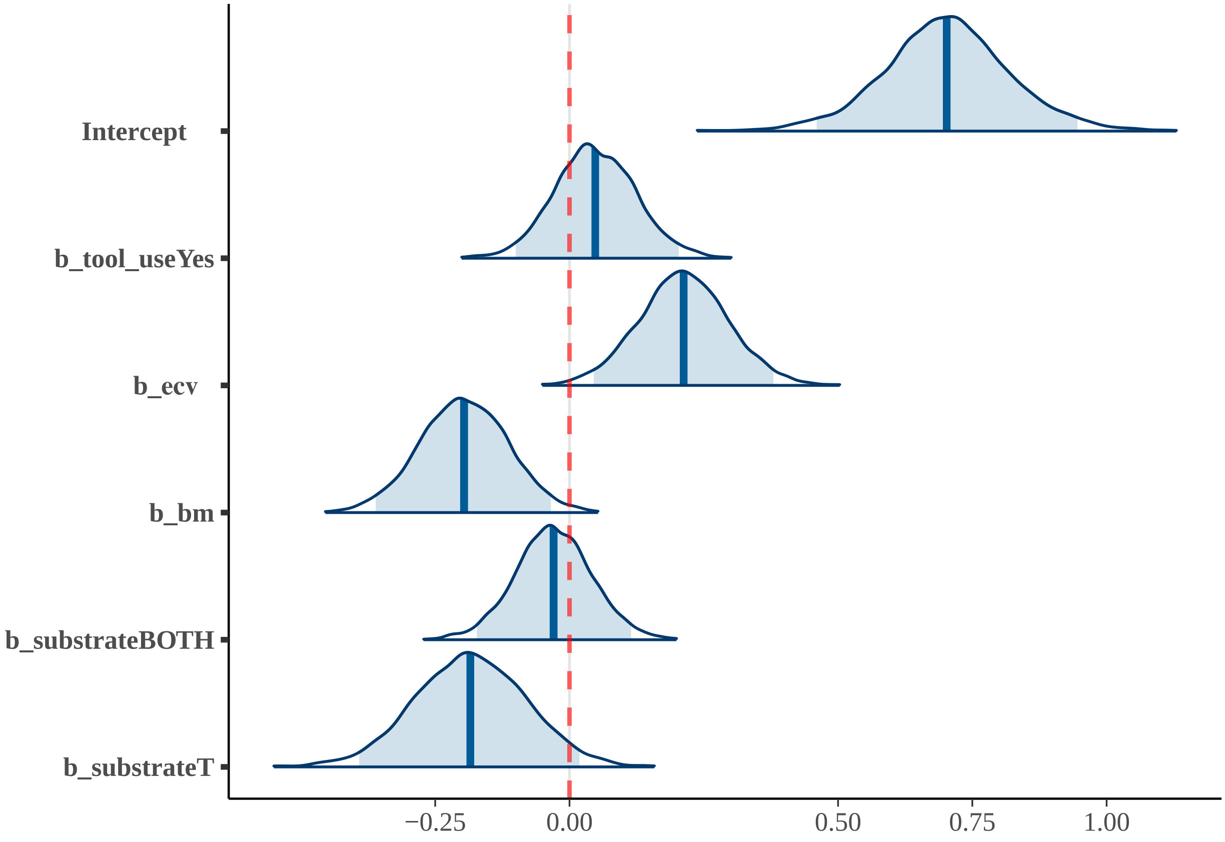


### 6) TU-BH

#### Summary TU-BH

| effect | component | term | estimate | std.error | conf.low | conf.high | rhat |
|---|---|---|---|---|---|---|---|
| fixed | cond | (Intercept) | 0.72 | 0.10 | 0.52 | 0.91 | 1 |
| fixed | cond | tool_useYes | 0.00 | 0.07 | -0.14 | 0.15 | 1 |
| fixed | cond | ecv | 0.18 | 0.08 | 0.02 | 0.33 | 1 |
| fixed | cond | bm | -0.12 | 0.08 | -0.29 | 0.05 | 1 |
| fixed | cond | substrateBOTH | -0.10 | 0.08 | -0.25 | 0.06 | 1 |
| fixed | cond | substrateT | -0.29 | 0.11 | -0.50 | -0.07 | 1 |
| fixed | cond | imi | -0.11 | 0.05 | -0.19 | -0.01 | 1 |

| effect | component | group | term | estimate | std.error | conf.low | conf.high | rhat |
|---|---|---|---|---|---|---|---|---|
| ran_pars | cond | obs | sd___(Intercept) | 0.10 | 0.02 | 0.06 | 0.14 | 1 |
| ran_pars | cond | phylo | sd___(Intercept) | 0.03 | 0.01 | 0.00 | 0.05 | 1 |
| ran_pars | cond | species | sd___(Intercept) | 0.04 | 0.03 | 0.00 | 0.11 | 1 |

### Posterior predictive check TU-BH; 100 simulated datasets

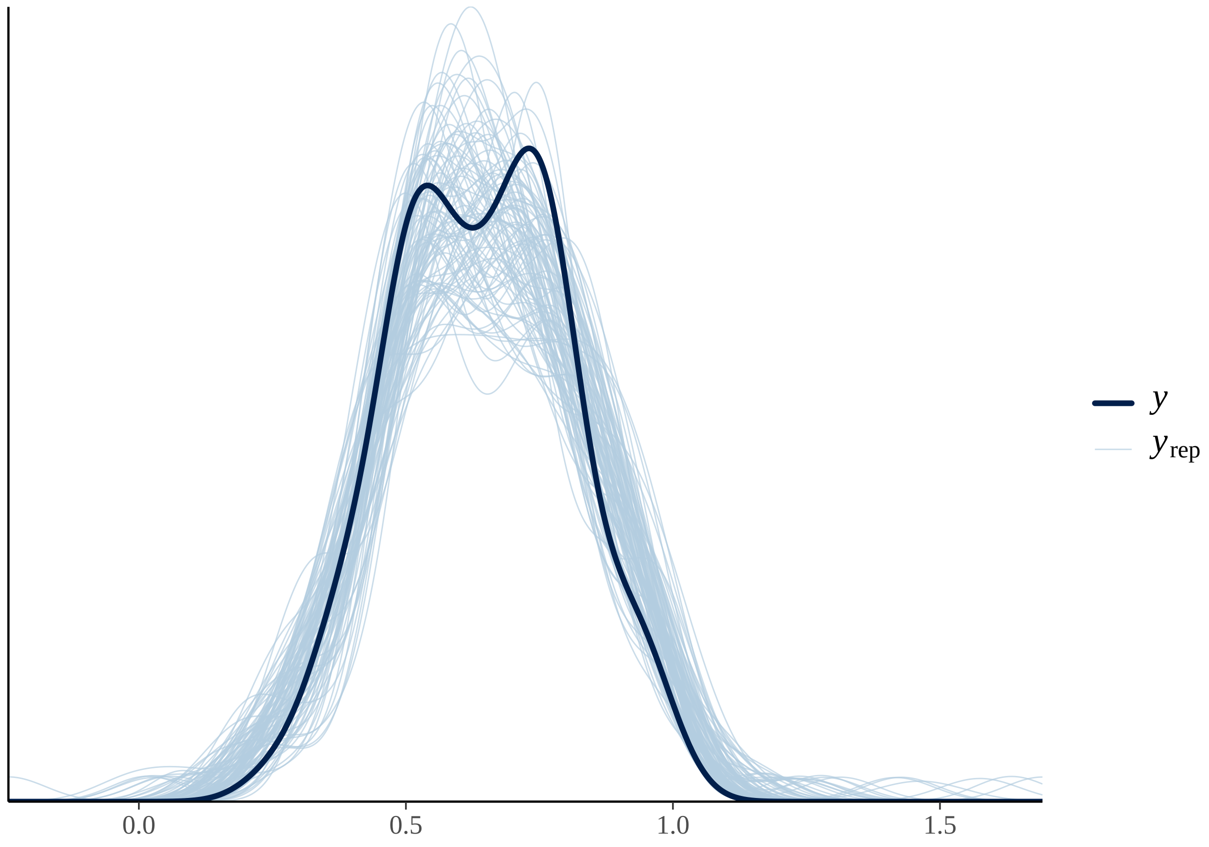

#### Fixed effects results TU-BH

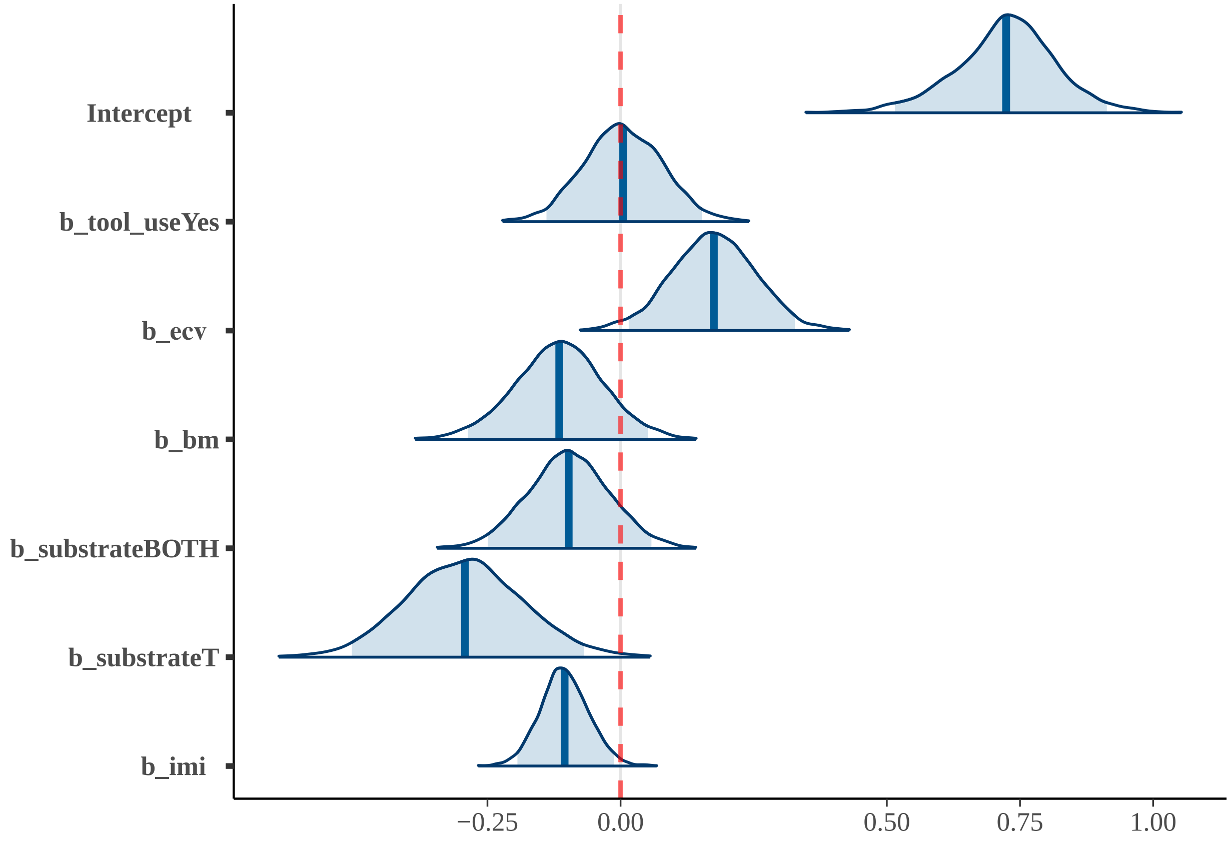


##### 7) TU-SH-SSH

#### Summary TU-SH-SSH

| effect | component | term | estimate | std.error | conf.low | conf.high | rhat |
|---|---|---|---|---|---|---|---|
| fixed | cond | (Intercept) | 0.74 | 0.11 | 0.51 | 0.96 | 1 |
| fixed | cond | tool_useYes | 0.11 | 0.08 | -0.04 | 0.27 | 1 |
| fixed | cond | substrateBOTH | -0.09 | 0.08 | -0.25 | 0.08 | 1 |
| fixed | cond | substrateT | -0.19 | 0.10 | -0.39 | 0.02 | 1 |
| fixed | cond | social_systemPair | -0.19 | 0.12 | -0.42 | 0.05 | 1 |
| fixed | cond | social_systemSolitary | -0.30 | 0.19 | -0.68 | 0.07 | 1 |

| effect | component | group | term | estimate | std.error | conf.low | conf.high | rhat |
|---|---|---|---|---|---|---|---|---|
| ran_pars | cond | obs | sd__(Intercept) | 0.10 | 0.02 | 0.07 | 0.15 | 1 |
| ran_pars | cond | phylo | sd__(Intercept) | 0.03 | 0.01 | 0.00 | 0.06 | 1 |
| ran_pars | cond | species | sd__(Intercept) | 0.05 | 0.03 | 0.00 | 0.12 | 1 |

**Posterior predictive check TU-SH-SSH; 100 simulated datasets**

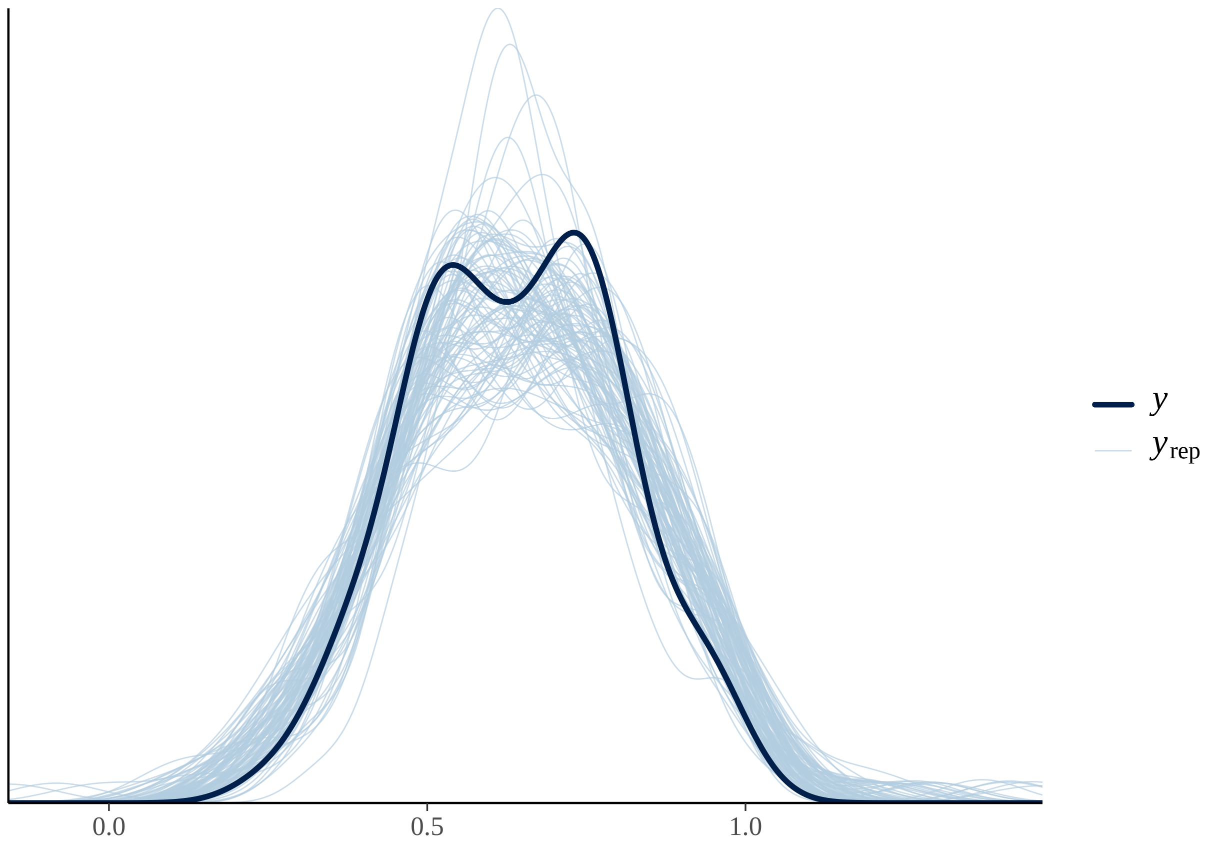

#### Fixed effects results TU-SH-SSH

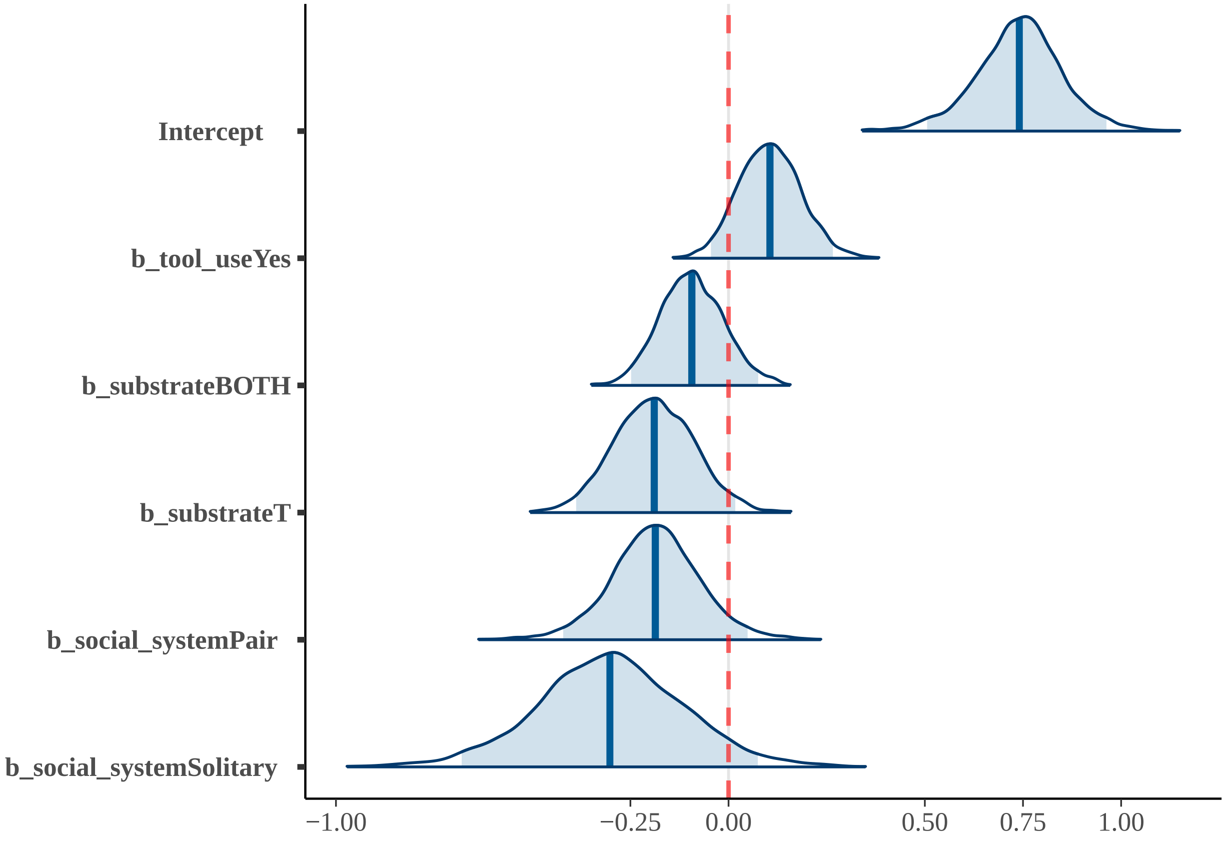


##### 8) SP-SS-B-TUH

#### Summary SP-SS-B-TUH

| effect | component | term | estimate | std.error | conf.low | conf.high | rhat |
|---|---|---|---|---|---|---|---|
| fixed | cond | (Intercept) | 0.71 | 0.12 | 0.45 | 0.96 | 1 |
| fixed | cond | tool_useYes | 0.08 | 0.08 | -0.08 | 0.24 | 1 |
| fixed | cond | substrateBOTH | -0.09 | 0.09 | -0.26 | 0.08 | 1 |
| fixed | cond | substrateT | -0.22 | 0.12 | -0.46 | 0.02 | 1 |
| fixed | cond | social_systemPair | -0.09 | 0.18 | -0.46 | 0.27 | 1 |
| fixed | cond | social_systemSolitary | -0.10 | 0.24 | -0.58 | 0.37 | 1 |
| fixed | cond | bm | 0.01 | 0.07 | -0.13 | 0.15 | 1 |
| fixed | cond | imi | -0.10 | 0.07 | -0.24 | 0.05 | 1 |

| effect | component | group | term | estimate | std.error | conf.low | conf.high | rhat |
|---|---|---|---|---|---|---|---|---|
| ran_pars | cond | obs | sd__(Intercept) | 0.10 | 0.02 | 0.07 | 0.14 | 1 |
| ran_pars | cond | phylo | sd__(Intercept) | 0.03 | 0.01 | 0.01 | 0.06 | 1 |
| ran_pars | cond | species | sd__(Intercept) | 0.04 | 0.03 | 0.00 | 0.11 | 1 |

**Posterior predictive check SP-SS-B-TUH; 100 simulated datasets**

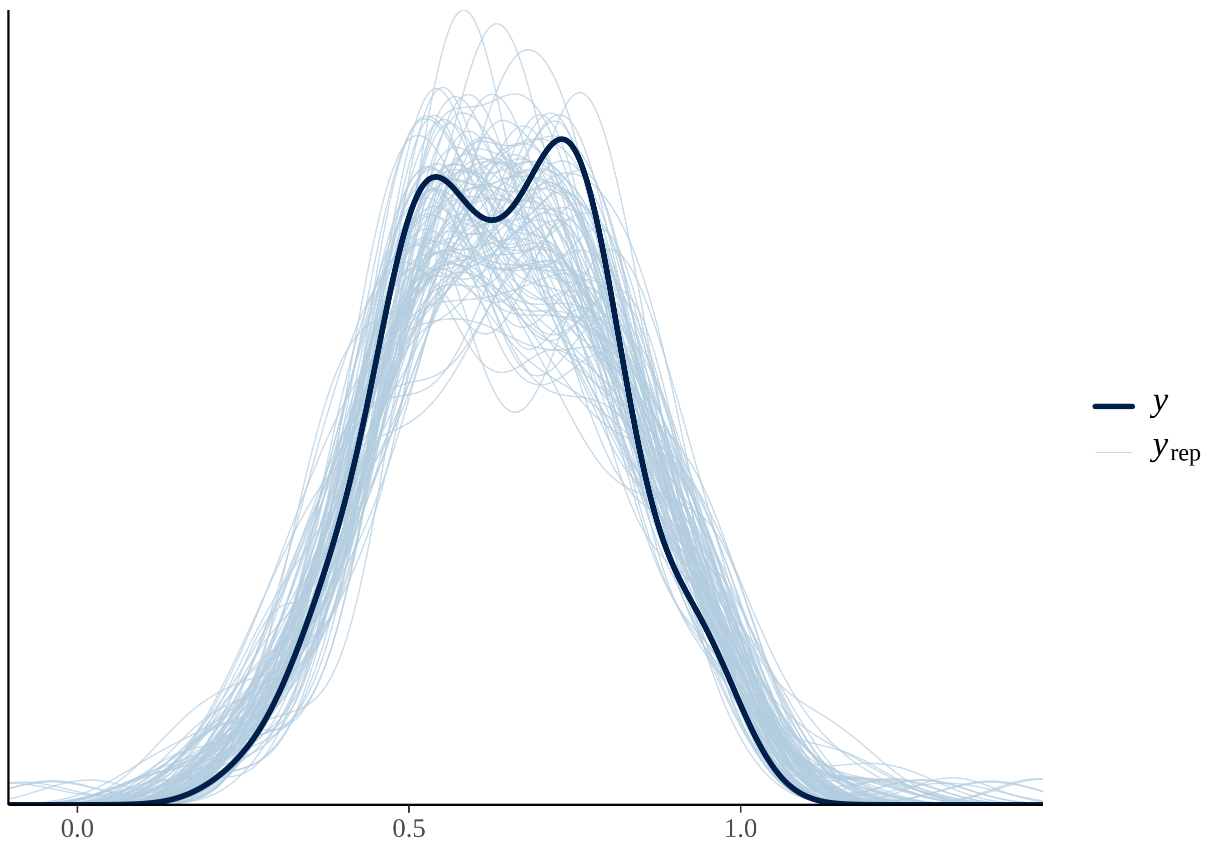

#### Fixed effects results SP-SS-B-TUH

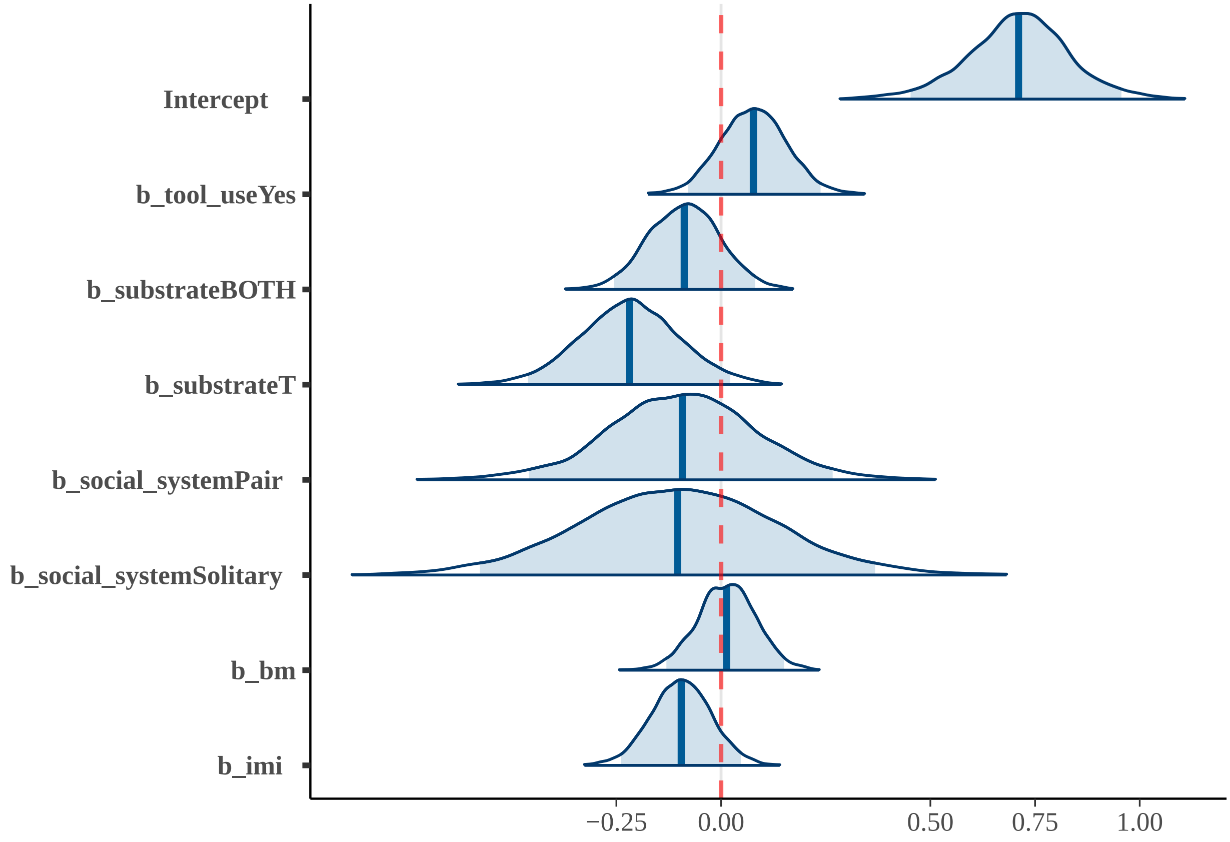


### 9) FH

#### Summary FH

| effect | component | term | estimate | std.error | conf.low | conf.high | rhat |
|---|---|---|---|---|---|---|---|
| fixed | cond | (Intercept) | 0.65 | 0.15 | 0.37 | 0.97 | 1 |
| fixed | cond | dimo | -0.08 | 0.04 | -0.16 | -0.01 | 1 |
| fixed | cond | cl2 | -0.03 | 0.13 | -0.31 | 0.23 | 1 |
| fixed | cond | cl3 | 0.10 | 0.15 | -0.21 | 0.38 | 1 |
| fixed | cond | cl4 | -0.02 | 0.16 | -0.36 | 0.28 | 1 |

| effect | component | group | term | estimate | std.error | conf.low | conf.high | rhat |
|---|---|---|---|---|---|---|---|---|
| ran_pars | cond | obs | sd__(Intercept) | 0.10 | 0.02 | 0.07 | 0.14 | 1 |
| ran_pars | cond | phylo | sd__(Intercept) | 0.03 | 0.01 | 0.01 | 0.06 | 1 |
| ran_pars | cond | species | sd__(Intercept) | 0.04 | 0.03 | 0.00 | 0.11 | 1 |

#### Posterior predictive check FH; 100 simulated datasets

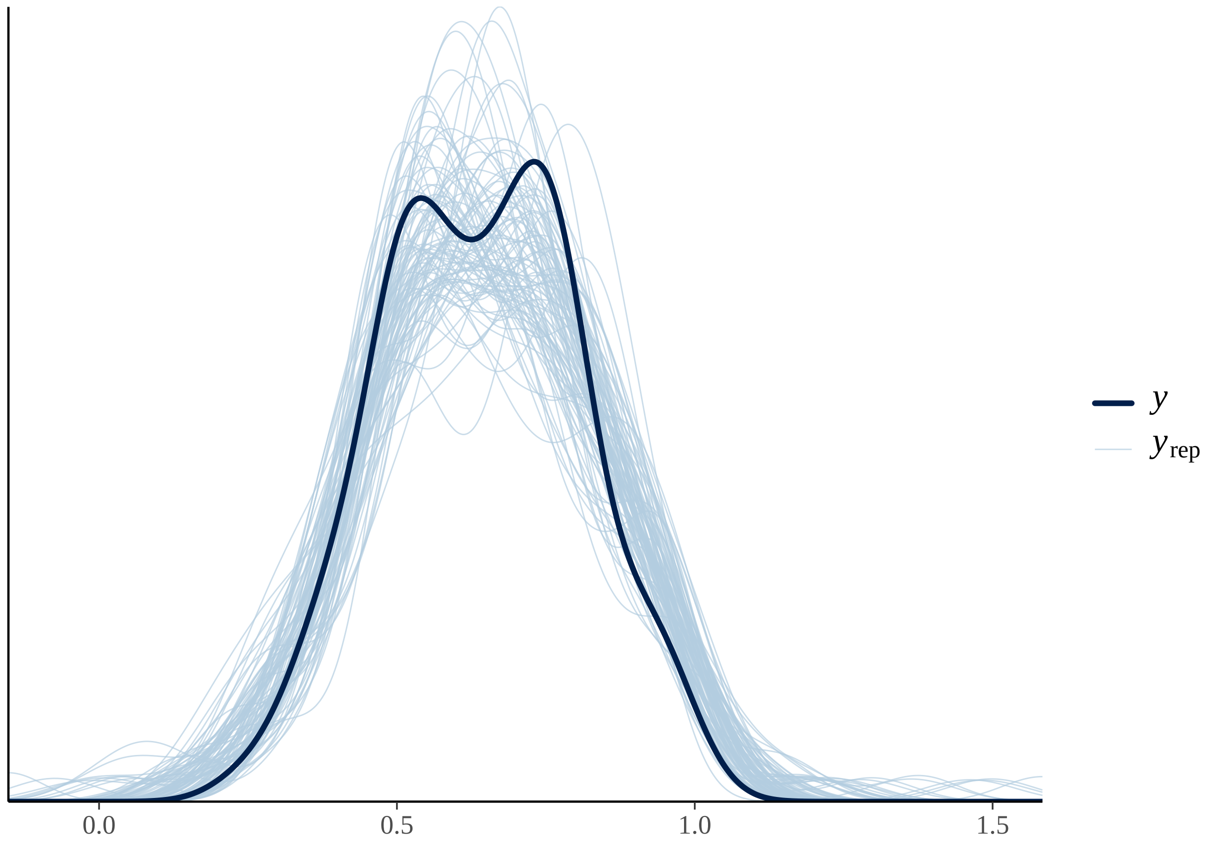

#### Fixed effects results FH

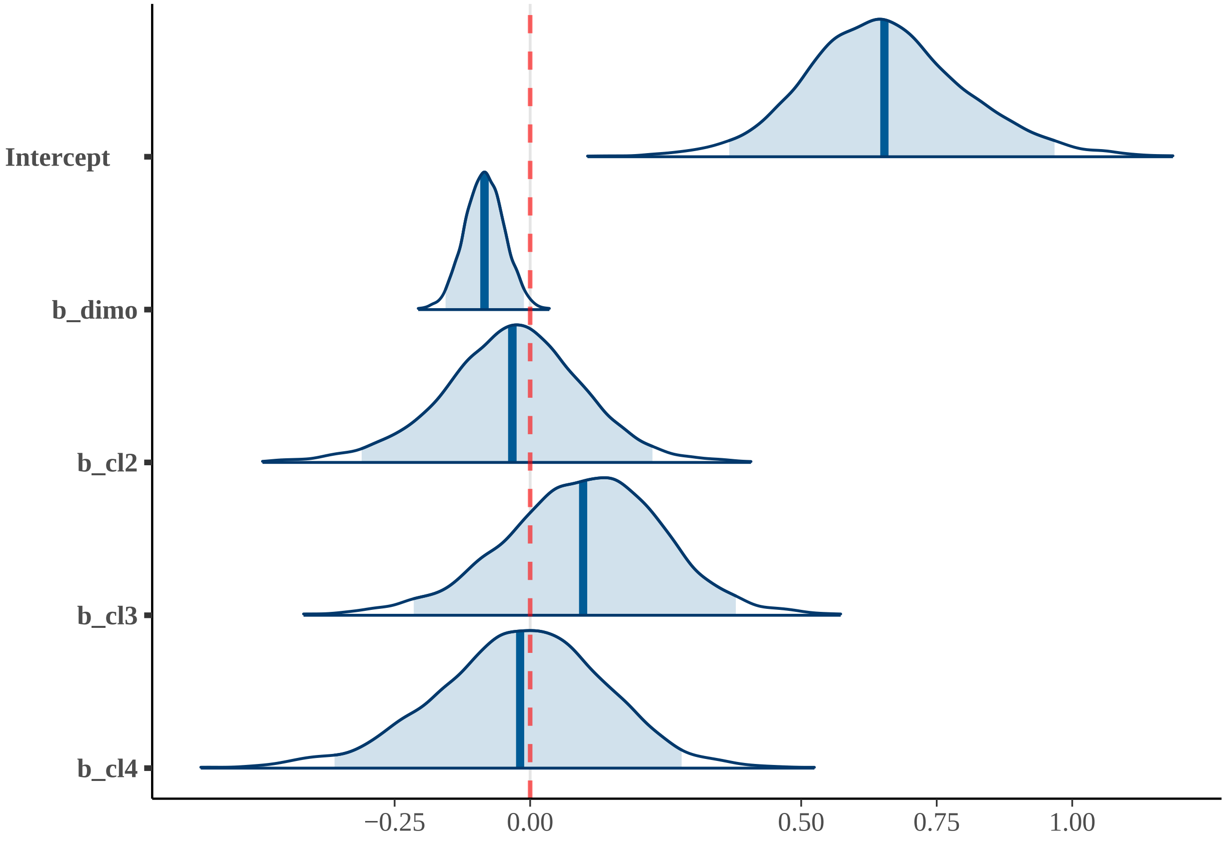


##### 10) EFH

#### Summary EFH

| effect | component | term | estimate | std.error | conf.low | conf.high | rhat |
|---|---|---|---|---|---|---|---|
| fixed | cond | (Intercept) | 0.68 | 0.12 | 0.43 | 0.92 | 1 |
| fixed | cond | extractive | -0.06 | 0.09 | -0.23 | 0.12 | 1 |
| fixed | cond | social_learning | -0.02 | 0.07 | -0.17 | 0.12 | 1 |
| fixed | cond | tool_useYes | 0.13 | 0.08 | -0.03 | 0.28 | 1 |

| effect | component | group | term | estimate | std.error | conf.low | conf.high | rhat |
|---|---|---|---|---|---|---|---|---|
| ran_pars | cond | obs | sd__(Intercept) | 0.11 | 0.02 | 0.07 | 0.15 | 1 |
| ran_pars | cond | phylo | sd__(Intercept) | 0.03 | 0.01 | 0.01 | 0.06 | 1 |
| ran_pars | cond | species | sd__(Intercept) | 0.05 | 0.03 | 0.00 | 0.13 | 1 |

### Posterior predictive check EFH; 100 simulated datasets

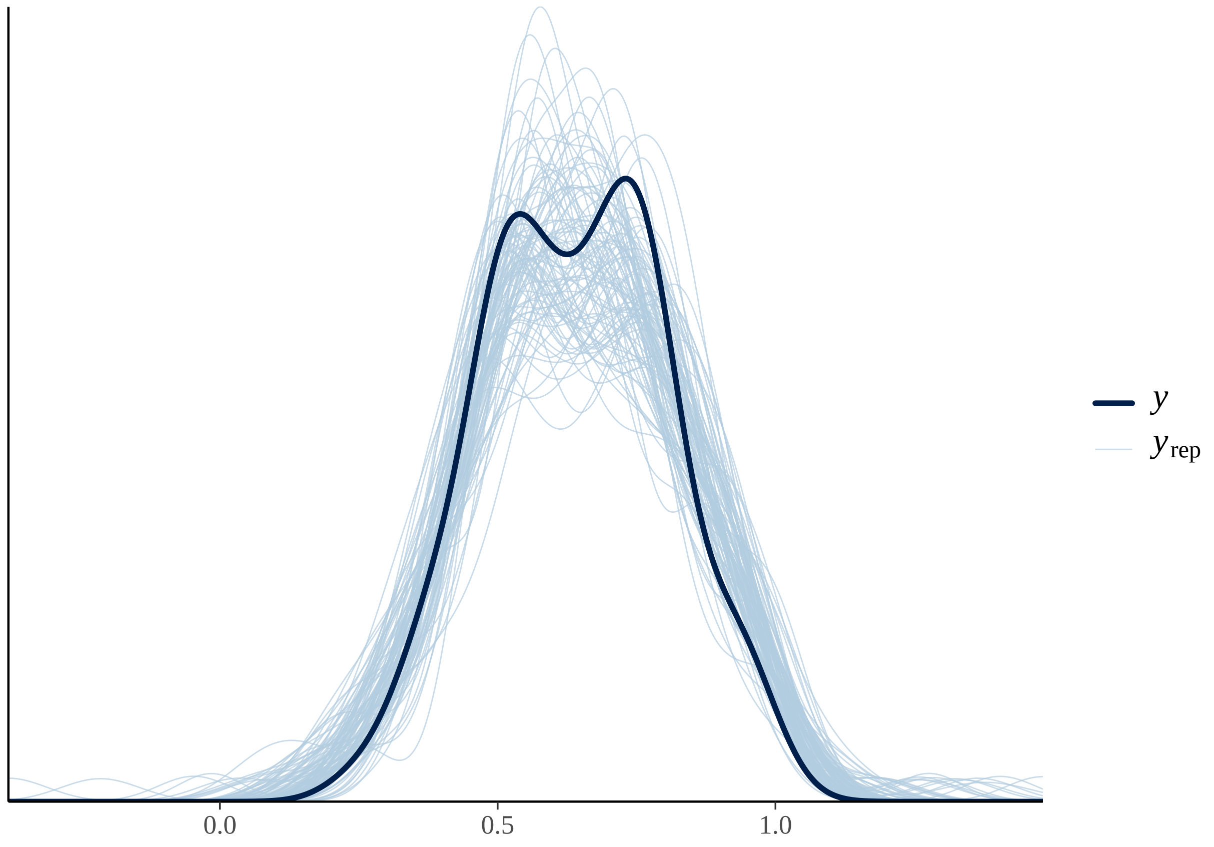

#### Fixed effects results EFH

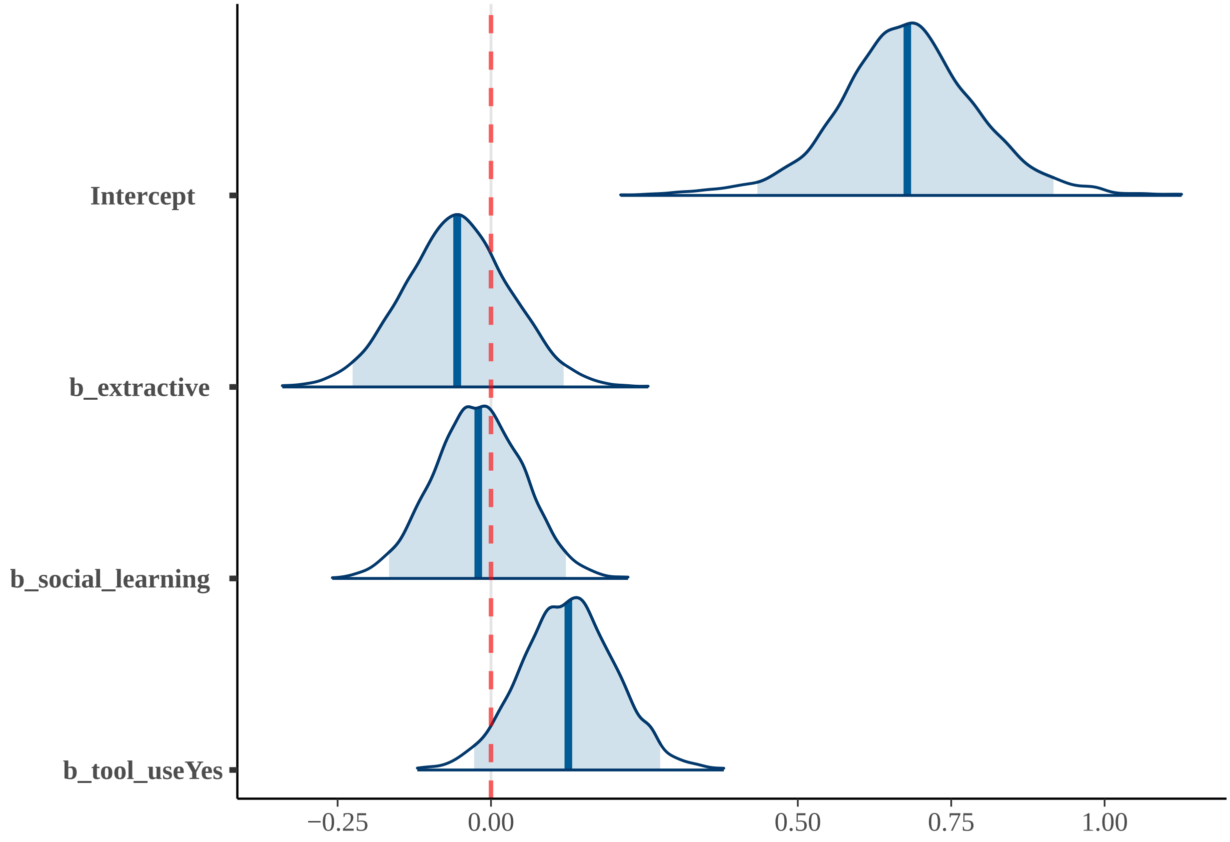


##### 11) Reduced model

#### Summary Reduced model

| effect | component | term | estimate | std.error | conf.low | conf.high | rhat |
|---|---|---|---|---|---|---|---|
| fixed | cond | (Intercept) | 0.73 | 0.09 | 0.54 | 0.91 | 1 |
| fixed | cond | ecv | 0.09 | 0.04 | 0.01 | 0.16 | 1 |
| fixed | cond | imi | -0.13 | 0.04 | -0.21 | -0.05 | 1 |
| fixed | cond | substrateBOTH | -0.13 | 0.07 | -0.27 | 0.02 | 1 |
| fixed | cond | substrateT | -0.31 | 0.11 | -0.52 | -0.10 | 1 |

| effect | component | group | term | estimate | std.error | conf.low | conf.high | rhat |
|---|---|---|---|---|---|---|---|---|
| ran_pars | cond | obs | sd__(Intercept) | 0.10 | 0.02 | 0.07 | 0.14 | 1 |
| ran_pars | cond | phylo | sd__(Intercept) | 0.03 | 0.01 | 0.00 | 0.05 | 1 |
| ran_pars | cond | species | sd__(Intercept) | 0.04 | 0.03 | 0.00 | 0.10 | 1 |

#### Posterior predictive check Reduced model; 100 simulated datasets

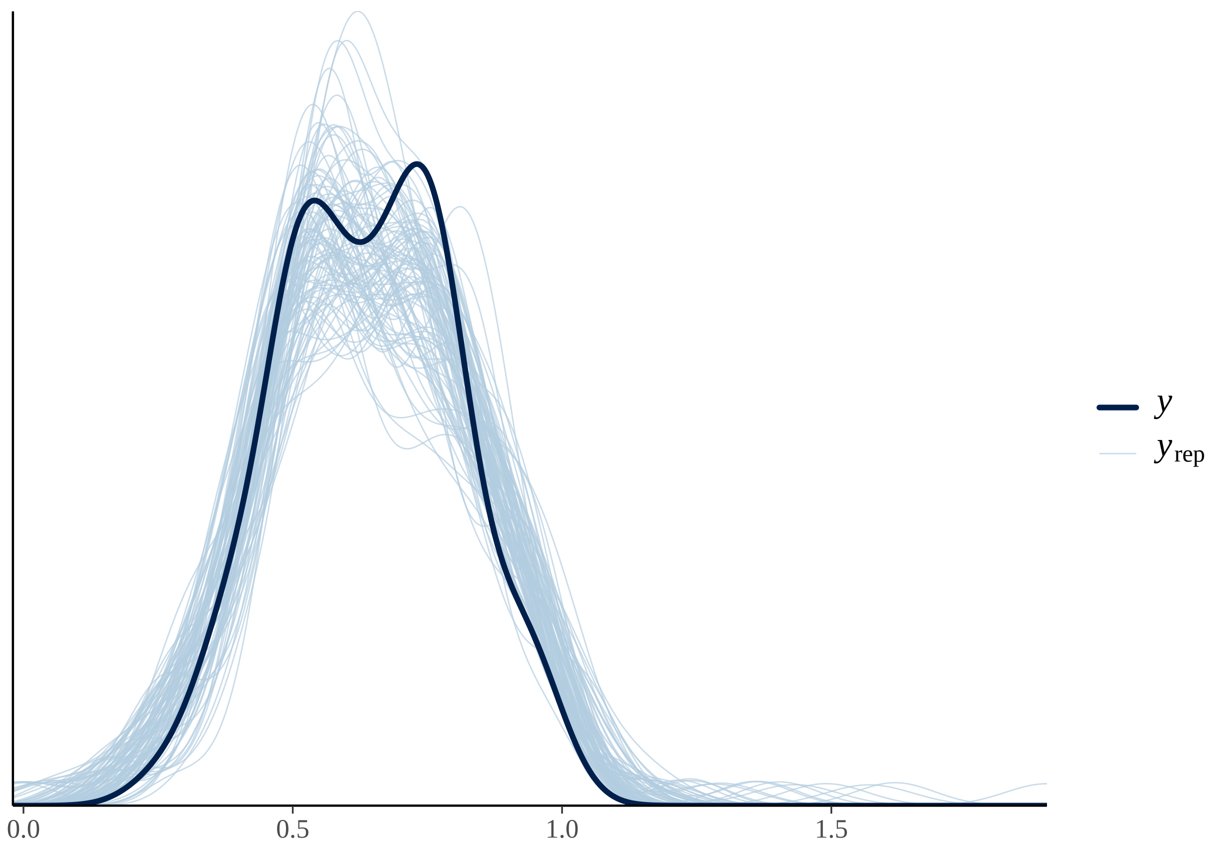

#### Fixed effects results Reduced model

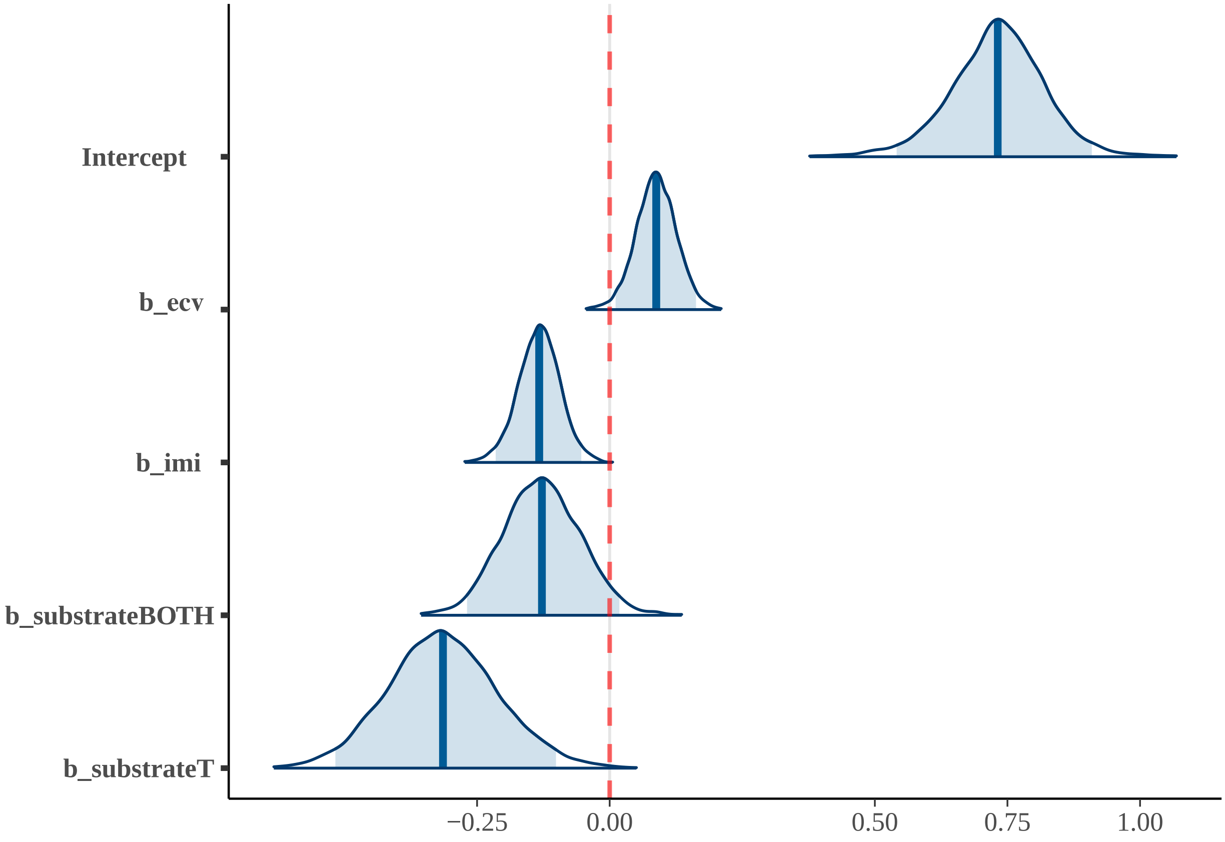
